# *APOE4* is Associated with Differential Regional Vulnerability to Bioenergetic Deficits in Aged *APOE* Mice

**DOI:** 10.1101/482422

**Authors:** Tal Nuriel, Delfina Larrea, David N. Guilfoyle, Leila Pirhaji, Kathleen Shannon, Hirra Arain, Archana Ashok, Marta Pera, Qiuying Chen, Allissa A. Dillman, Helen Y. Figueroa, Mark R. Cookson, Steven S. Gross, Ernest Fraenkel, Karen E. Duff, Estela Area-Gomez

**Affiliations:** Taub Institute for Research on Alzheimer’s Disease and the Aging Brain, Columbia University, 630 West 168th Street, New York, NY 10032, USA.; Department of Pathology and Cell Biology, Columbia University, 630 West 168th Street, New York, NY 10032, USA.; Department of Neurology, Columbia University, 630 West 168th Street, New York, NY 10032, USA.; Center for Biomedical Imaging and Neuromodulation, Nathan S. Kline Institute, Orangeburg, NY 10962, USA.; Department of Biological Engineering, Massachusetts Institute of Technology, 350 Brookline Street, Cambridge, MA 02139, USA.; Animal Facility, Nathan S. Kline Institute, Orangeburg, NY 10962, USA.; Department of Pharmacology, Weill Cornell Medical College, 1300 York Avenue, New York, NY 10065, USA.; Cell Biology and Gene Expression Section, Laboratory of Neurogenetics, National Institute on Aging, National Institutes of Health, Bethesda, Maryland, 20892; Broad Institute, 415 Main Street, Cambridge, MA 02142, Cambridge, Massachusetts, 02139, USA.; Division of Integrative Neuroscience in the Department of Psychiatry, New York State Psychiatric Institute, 630 West 168th Street, New York, NY 10032, USA.

## Abstract

The ε4 allele of apolipoprotein E (*APOE*) is the dominant genetic risk factor for late-onset Alzheimer’s disease (AD). However, the reason for the association between *APOE4* and AD remains unclear. While much of the research has focused on the ability of the apoE4 protein to increase the aggregation and decrease the clearance of Aβ, there is also an abundance of data showing that *APOE4* negatively impacts many additional processes in the brain, including bioenergetics. In order to gain a more comprehensive understanding of the *APOE4*’s role in AD pathogenesis, we performed a multi-omic analysis of *APOE4* vs. *APOE3* expression in the entorhinal cortex (EC) and primary visual cortex (PVC) of aged *APOE* mice. These studies revealed region-specific alterations in several bioenergetic pathways, including oxidative phosphorylation (OxPhos), the TCA-cycle and fatty acid metabolism. Follow-up analysis utilizing the Seahorse platform revealed decreased mitochondrial respiration in the hippocampus and cortex of aged *APOE4* vs. *APOE3* mice, but not in the EC of these mice. Additional studies, as well as the original multi-omic data suggest that bioernergetic pathways in the EC of aged *APOE* mice may be differentially regulated by *APOE4* expression. Given the importance of the EC as one of the first regions to be affected by AD pathology in humans, this differential bionenergetic regulation observed in the EC vs. other brain regions of aged *APOE4* mice may play an important role in the pathogenesis of AD, particularly among *APOE4* carriers.

## INTRODUCTION

Possession of the ε4 allele of apolipoprotein E (*APOE*) is the major genetic risk factor for late-onset Alzheimer’s disease (AD). In normal physiology, the apoE protein plays a vital role in the transport of cholesterol and other lipids through the bloodstream, as well as within the brain (*1-3*). Although the three common isoforms of apoE (E2, E3 and E4) differ from each other by only two amino acids—apoE2 (Cys112, Cys158), apoE3 (Cys112, Arg158), apoE4 (Arg112, Arg158)—this small change in amino acid sequence has a large effect on protein structure, causing the apoE4 protein to possess a more globular structure due to increased interaction between its N- and C-terminal domains. This difference in structure, in turn, affects the type of lipids that each isoform binds, as well as the affinity of each isoform for the numerous receptors that mediate the uptake of apoE and its cargo into cells (see reviews by (1, 4)).

Importantly, these isoform differences also have a major impact on the pathogenesis of late-onset AD. Although the *APOE2, APOE3* and *APOE4* alleles are present at a relative frequency of about 8%, 77% and 15% in the normal population (5, 6), the *APOE4* allele is present at a relative frequency of 36-64% in AD patients (7-10), with individuals who possess one or two *APOE4* alleles having a 3- or 12-fold increased risk of developing AD, respectively (11). Although a number of mechanisms have been proposed to help explain this *APOE4*-associated susceptibility to AD, the precise cause remains a source of debate. The prominent hypothesis is that this increase in AD risk is due to the ability of apoE4 to increase the aggregation and decrease the clearance of Aβ (12-18). However, *APOE4* expression has also been shown to have deleterious effects on numerous Aβ-independent pathways, including lipid metabolism, tau pathology, bioenergetics, neuronal development, synaptic plasticity, the neuro-vasculature, and neuro-inflammation [see reviews by (19) and (20)], any number of which could play a pivotal role in the pathogenesis of AD among *APOE4* carriers.

In terms of *APOE4*’s effects on bioenergetics, a number of pivotal reports have demonstrated that *APOE4* expression leads to widespread dysregulation of the brain’s bioenergetic capacity. For example, early reports by Eric Reiman and colleagues demonstrated that both young and old *APOE4* carriers display decreased glucose utilization, as measured by fluorodeoxyglucose positron emission tomography (FDG-PET), in similar brain regions to that seen with AD patients (21-23). Additional reports detail the wide-range of bioenergetic insults that *APOE4* expression can cause in the brain, including impaired insulin signaling (24, 25), reduced cerebral blood volume and cognitive function in response to a high fat diet (HFD) (26, 27), altered genetic expression of glucose-regulating enzymes and transporters (28, 29), and the generation of a toxic C-terminal fragment of apoE4 that can directly target electron transport chain (ETC) complexes in the mitochondria (30-32).

In order to study the diverse effects that *APOE4* expression has in the brain, we performed an initial multi-omic study of differential *APOE* isoform expression in an AD-vulnerable vs. an AD-resistant brain region from aged *APOE* targeted replacement mice, which express human *APOE* in place of their mouse *Apoe* gene and do not develop overt AD pathology. The EC was chosen as our AD-vulnerable brain region because it is one of the first brain regions to develop AD-related tau pathology, in humans (33, 34), and also because the morphology of the EC has been shown to be specifically affected by *APOE4* gene expression. (35, 36). The PVC was chosen as our AD-resistant region because it is relatively spared in AD (33, 37, 38). As described below, these studies revealed *APOE4*-specific alterations in the levels of numerous genes and small-molecules related to energy metabolism, which were explored further using the Seahorse platform. In agreement with previous data, this analysis revealed *APOE4*-associated deficits in mitochondrial respiration in several brain regions. However, contrary to what we observed in other brain regions, the EC displayed increases in oxidative metabolism in the context of *APOE4* expression, suggesting that, under metabolic stress, brain region-specific counterbalancing mechanisms are in play in the EC of aged *APOE4* mice. Additional studies and further mining of the multi-omic data was then used to elucidate this differential bioenergetic regulation in the EC of aged *APOE4* mice, which may play a causative role in the pathogenesis of AD and may point to novel therapeutic interventions for preventing or slowing the disease.

## RESULTS

### Multi-omic analysis reveals differential expression of energy-related genes and metabolites in the EC of aged *APOE4* mice

In order to investigate the effects of *APOE4* expression in an untargeted manner, we performed a multi-omic analysis (transcriptomics and metabolomics) on RNA and small-molecules extracted from an AD-vulnerable brain region (the EC) vs. an AD-resistant brain region (the PVC) of 14-15 month-old *APOE* targeted replacement mice. For the transcriptomic analysis, RNA was Trizol-extracted from the EC and PVC of aged *APOE* mice (10 *APOE3/3* and 19 *APOE3/*4 males). For the metabolomics analysis, small-molecules were extracted from the EC and PVC of aged *APOE* mice (8 *APOE3/3*, 9 *APOE3/4* and 7 *APOE4/4* males) using a methyl tert-butyl ether (MTBE)/methanol extraction protocol modified from previous reports (39, 40). The RNA and small-molecules were then analyzed using RNA-sequencing and untargeted metabolomics, respectively.

The data generated from these transcriptomics and metabolomics analyses (summarized in Tables 1 and 2 and Supplementary Tables S1 and S2) have recently been published as parts of larger studies on *APOE4*’s impact on endosomal-lysosomal dysregulation (41) and neuronal hyperactivity (42), respectively. However, as is often the case with omics and other systems biology approaches, the large amounts of data generated from these analyses inform us about changes in multiple biological pathways that cannot all be investigated in one study. In the case of the transcriptomics analysis, in addition to the dysregulation of endosomal-lysosomal genes that were revealed in our pathway analysis (and which we investigated further for our previously published study), another major KEGG pathway that was observed to be significantly enriched for differentially expressed genes was “Oxidative Phosphorylation” (Fig. 1A). This was driven by a large number of differentially expressed genes encoding for subunits of ETC complexes I-V, with each of these genes being upregulated in the EC of aged *APOE3/4* mice, as compared to aged *APOE3/3* mice (Fig. 1B). This increased transcription of ETC genes may also be related to the increases in energy-related metabolites that we observed in the EC of aged *APOE4/4* mice (which we reported as part of our neuronal hyperactivity study). These metabolites include several TCA cycle metabolites (malate, citrate and isocitrate), as well as fructose-6-phosphate, carnitine, and ATP, each of which had higher levels in the EC of aged *APOE4/4* vs. *APOE3/3* mice (Fig. 2A). Furthermore, the metabolomics data indicates a higher ATP:ADP ratio in the EC of aged *APOE4/4* vs. *APOE3/3* mice (Fig. 2C), which suggests an increase in the rate of oxidative metabolism in this region. Thus, in addition to the *APOE4*-associated differences in endosomal-lysosomal regulation and neuronal activity that these original studies highlighted, the data from both the transcriptomics and metabolomics studies also pointed to specific differences in several bioenergetic pathways.

**Table 1.** Differentially expressed targeted metabolites from the EC of APOE3 vs. E4 mice

| Metabolite | Regulation in E4/4 | Fold Change | p-value | FDR | Detection Mode | CAS Number |
| --- | --- | --- | --- | --- | --- | --- |
| **Fatty Acids** |  |  |  |  |  |  |
| gamma-Linolenic Acid | down | 1.31 | 1.19E-03 | 0.093 | positive | 506-26-3 |
| Myristic Acid | down | 3.92 | 0.004 | 0.118 | positive | 544-63-8 |
| Docosahexaenoic Acid | up | 1.36 | 0.011 | 0.154 | positive | 6217-54-5 |
| Stearic Acid | down | 1.69 | 0.011 | 0.154 | positive | 57-11-4 |
| 12-Hydroxydodecanoic Acid | down | 1.36 | 0.021 | 0.213 | positive | 505-95-3 |
| Arachidic Acid | down | 1.40 | 0.037 | 0.230 | negative | 506-30-9 |
| Palmitic Acid | down | 1.32 | 0.049 | 0.243 | negative | 57-10-3 |
| **Oligosaccharides** |  |  |  |  |  |  |
| trisaccharide | up | 1.99 | 0.001 | 0.049 | negative | 512-69-6 |
| tetrasaccharide | up | 2.24 | 0.015 | 0.169 | negative | 10094-58-3 |
| disaccharide | up | 2.20 | 0.015 | 0.169 | negative | 63-42-3 |
| **Vitamin and Vitamin Derivatives** |  |  |  |  |  |  |
| Phylloquinone | up | 1.47 | 0.003 | 0.102 | positive | 84-80-0 |
| Tocopherol | up | 2.74 | 0.005 | 0.123 | negative | 59-02-9 |
| Dehydroascorbic acid | up | 1.76 | 0.008 | 0.134 | positive | 490-83-5 |
| **Energy-Related Metabolites** |  |  |  |  |  |  |
| Inosine 5'-monophosphate (IMP) | up | 2.26 | 0.003 | 0.072<br>0 | negative | 131-99-7 |
| D-Fructose 6-phosphate | up | 1.10 | 0.011 | 0.169 | negative | 643-13-0 |
| Succinoadenosine | up | 1.47 | 0.011 | 0.169 | negative | 4542-23-8 |
| Carnitine | up | 1.27 | 0.021 | 0.213 | positive | 541-15-1 |
| Citric Acid/Isocitric Acid | up | 1.24 | 0.028 | 0.230 | negative | 77-92-9/<br>1637-73-6 |
| Malic Acid | up | 1.16 | 0.037 | 0.230 | negative | 97-67-6 |
| ATP | up | 1.24 | 0.049 | 0.243 | negative | 56-65-5 |
| **Cholesterol Metabolites** |  |  |  |  |  |  |
| Lanosterol | up | 2.27 | 0.015 | 0.169 | negative | 79-63-0 |
| Cholesteryl Acetate | up | 2.29 | 0.015 | 0.169 | negative | 604-35-3 |
| **Amino Acids** |  |  |  |  |  |  |
| Leucine | up | 1.26 | 0.015 | 0.169 | negative | 61-90-5 |
| Proline | up | 1.12 | 0.037 | 0.230 | negative | 147-85-3 |
| Glycine | up | 1.10 | 0.037 | 0.230 | negative | 56-40-6 |
| **Tryptophan Metabolites** |  |  |  |  |  |  |
| Quinaldic Acid | up | 1.68 | 0.001 | 0.049 | negative | 93-10-7 |
| Kynurenine | up | 2.49 | 0.015 | 0.169 | negative | 2922-83-0 |
| Kynurenic Acid | up | 1.32 | 0.049 | 0.243 | negative | 492-27-3 |
| **Cysteine and Methionine Metabolites** |  |  |  |  |  |  |
| S-Adenosylhomocysteine | up | 1.15 | 0.021 | 0.213 | positive | 979-92-0 |
| **Histidine Metabolism** |  |  |  |  |  |  |
| Carnosine | up | 1.31 | 0.037 | 0.230 | negative | 305-84-0 |
| **Arginine and Proline Metabolites** |  |  |  |  |  |  |
| 4-Oxoproline | up | 1.30 | 0.003 | 0.102 | positive | 4347-18-6 |
| **Tyrosine Metabolites** |  |  |  |  |  |  |
| Vanylglycol (MHPG) | up | 1.64 | 0.011 | 0.169 | negative | 67423-45-4 |
| Tyramine | up | 1.06 | 0.028 | 0.216 | positive | 51-67-2 |
| Thymidine | down | 1.49 | 0.028 | 0.216 | positive | 50-89-5 |
| Uracil | down | 1.38 | 0.049 | 0.243 | negative | 66-22-8 |
| **Purine Metabolism** |  |  |  |  |  |  |
| GMP | up | 1.14 | 0.028 | 0.230 | negative | 85-32-5 |
| **Miscellaneous** |  |  |  |  |  |  |
| Hydroxybutyric acid | up | 1.35 | 0.002 | 0.055 | negative | 5094-24-6 |
| Methylglutaryl carnitine | up | 2.42 | 0.005 | 0.134 | positive | 102673-95-0 |
| Trimethylamine N-oxide | up | 2.65 | 0.021 | 0.213 | positive | 1184-78-7 |
| 2-Hydroxypyridine | up | 1.36 | 0.037 | 0.275 | positive | 142-08-5 |
| N-Acetylneuraminic Acid | up | 1.23 | 0.037 | 0.230 | negative | 131-48-6 |

**Table 2.** Differentially expressed targeted metabolites from the PVC of APOE3 vs. E4 mice

| Metabolite | Regulation<br>in E4/4 | Fold<br>Change | p-value | FDR | Detection<br>Mode | CAS<br>Number |
| --- | --- | --- | --- | --- | --- | --- |
| Fatty Acids |  |  |  |  |  |  |
| gamma-Linolenic Acid | down | 1.19 | 0.003 | 0.068 | positive | 506-26-3 |
| Palmitic Acid | down | 1.32 | 0.021 | 0.212 | negative | 57-10-3 |
| 10-Hydroxydecanoate | up | 2.22 | 0.015 | 0.195 | positive | 1679-53-4 |
| Docosahexaenoic Acid | down | 2.03 | 0.021 | 0.212 | negative | 6217-54-5 |
| Arachidonic acid | down | 1.45 | 0.021 | 0.212 | negative | 506-32-1 |
| Oligosaccharides |  |  |  |  |  |  |
| trisaccharide | up | 1.82 | 0.015 | 0.212 | negative |  |
| tetrasaccharide | up | 2.70 | 0.021 | 0.212 | negative |  |
| Vitamin and Vitamin<br>Derivatives |  |  |  |  |  |  |
| Dehydroascorbic acid | up | 2.03 | 0.001 | 0.062 | positive | 490-83-5 |
| Phylloquinone | up | 1.90 | 0.001 | 0.062 | positive | 84-80-0 |
| Ascorbic acid | up | 105.84 | 0.005 | 0.192 | negative | 50-81-7 |
| Tocopherol | up | 2.28 | 0.008 | 0.192 | negative | 59-02-9 |
| Energy-Related<br>Metabolites |  |  |  |  |  |  |
| Acetylcarnitine | up | 1.40 | 0.003 | 0.068 | positive | 3040-38-8 |
| Carnitine | up | 1.31 | 0.011 | 0.154 | positive | 541-15-1 |
| Coenzyme A (CoA) | up | 2.82 | 0.028 | 0.282 | positive | 85-61-0 |
| Pyruvate | up | 1.43 | 0.028 | 0.238 | negative | 127-17-3 |
| Acetoacetic Acid | up | 1.40 | 0.028 | 0.238 | negative | 541-50-4 |
| Cholesterol Metabolites |  |  |  |  |  |  |
| Taurodeoxycholate | up | 1.94 | 0.003 | 0.068 | positive | 516-50-7 |
| Cholesteryl Acetate | up | 2.24 | 0.008 | 0.192 | negative | 604-35-3 |
| Lanosterol | up | 2.50 | 0.011 | 0.192 | negative | 79-63-0 |
| Amino Acids |  |  |  |  |  |  |
| Tyrosine | up | 1.25 | 0.037 | 0.270 | negative | 60-18-4 |
| Serine | down | 1.18 | 0.049 | 0.270 | negative | 56-45-1 |
| Glutamate | down | 1.08 | 0.049 | 0.282 | positive | 56-86-0 |
| Cysteine and Methionine<br>Metabolites |  |  |  |  |  |  |
| Pterin | down | 3.91 | 0.002 | 0.192 | negative | 2236-60-4 |
| Histidine Metabolites |  |  |  |  |  |  |
| Urocanic acid | down | 1.34 | 0.021 | 0.212 | negative | 104-98-3 |
| Carnosine | up | 1.32 | 0.028 | 0.282 | positive | 305-84-0 |
| Arginine and Proline<br>Metabolites |  |  |  |  |  |  |
| Phosphocreatine | up | 1.62 | 0.037 | 0.282 | positive | 67-07-2 |
| 4-Guanidinobutyric Acid | down | 1.09 | 0.049 | 0.270 | negative | 463-00-3 |
| **Threonine Metabolites** |  |  |  |  |  |  |
| 2-Ketobutyric Acid | up | 1.43 | 0.021 | 0.212 | negative | 600-18-0 |
| **Tyrosine Metabolites** |  |  |  |  |  |  |
| Homovanillic Acid | down | 1.30 | 0.049 | 0.270 | negative | 306-08-1 |
| **Riboflavin Metabolites** |  |  |  |  |  |  |
| 2,4-Dihydroxypteridine (Lumazine) | up | 2.07 | 0.008 | 0.154 | positive | 487-21-8 |
| Flavin adenine dinucleotide (FAD) | up | 1.19 | 0.021 | 0.212 | negative | 146-14-5 |
| Lumichrome | down | 1.71 | 0.049 | 0.282 | positive | 1086-80-2 |
| **Pyrimidine Metabolites** |  |  |  |  |  |  |
| 2-Aminoisobutyric acid | up | 1.38 | 0.011 | 0.154 | positive | 62-57-7 |
| UMP | up | 1.22 | 0.049 | 0.270 | negative | 58-97-9 |
| CMP | up | 1.29 | 0.049 | 0.282 | positive | 63-37-6 |
| **Miscellaneous** |  |  |  |  |  |  |
| Lipoic Acid | up | 3.76 | 0.005 | 0.192 | negative | 1077-28-7 |
| Thiourea | down | 1.45 | 0.008 | 0.192 | negative | 62-56-6 |
| Butyrylcarnitine | up | 1.24 | 0.011 | 0.154 | positive | 25576-40-6 |
| Hydroxybutyric Acid | up | 1.49 | 0.011 | 0.192 | negative |  |
| Indoxyl Sulfate | down | 2.12 | 0.021 | 0.212 | negative | 2642-37-7 |
| 3-Hydroxymethylglutaric Acid | up | 1.27 | 0.028 | 0.238 | negative | 503-49-1 |
| 2-Hydroxypyridine | up | 1.45 | 0.037 | 0.282 | positive | 142-08-5 |
| Maleic acid | up | 1.92 | 0.037 | 0.270 | negative | 110-16-7 |
| N-Acetylneuraminic Acid | up | 1.17 | 0.037 | 0.282 | positive | 131-48-6 |
| N-Methylglutamic Acid | down | 1.19 | 0.049 | 0.270 | negative | 35989-16-3 |
| 2-Aminoadipic Acid | down | 1.17 | 0.049 | 0.270 | negative | 542-32-5 |
| Cytidine diphosphate choline (CDPcholine) | down | 1.17 | 0.049 | 0.282 | positive | 987-78-0 |
| Glutathione | up | 1.49 | 0.049 | 0.282 | positive | 70-18-8 |

**Fig. 1.**
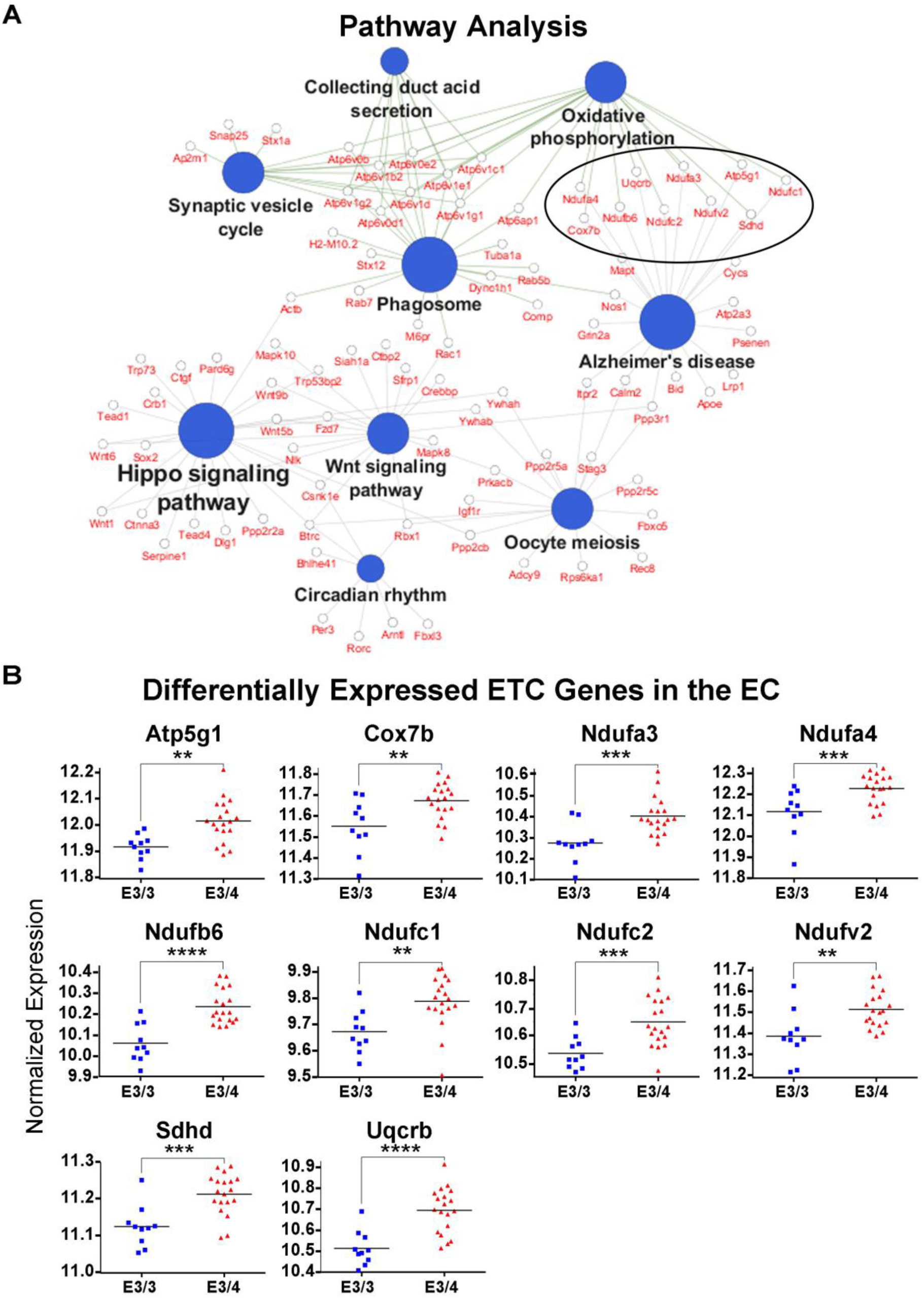
Transcriptomics analysis reveals an upregulation of electron transport chain genes in the EC of aged *APOE4* mice. RNA-sequencing was performed in order to analyze the effects of differential *APOE* isoform expression on RNA levels in the EC and PVC of aged *APOE* mice (19 *APOE3/4* males vs. 10 *APOE3/3* males). (A) Shown here are the significantly enriched KEGG pathways observed in the EC of *APOE3/4* vs. *APOE3/3* mice, using Cytoscape’s ClueGo application, with the ten differentially expressed electron transport chain (ETC) genes circled. (B) Graphs of the differentially regulated ETC genes in the EC of the *APOE3/4* vs. *APOE3/3* mice. (** denotes p < 0.01; *** denotes *p* < 0.001; **** denotes p < 0.0001)

**Fig. 2.**
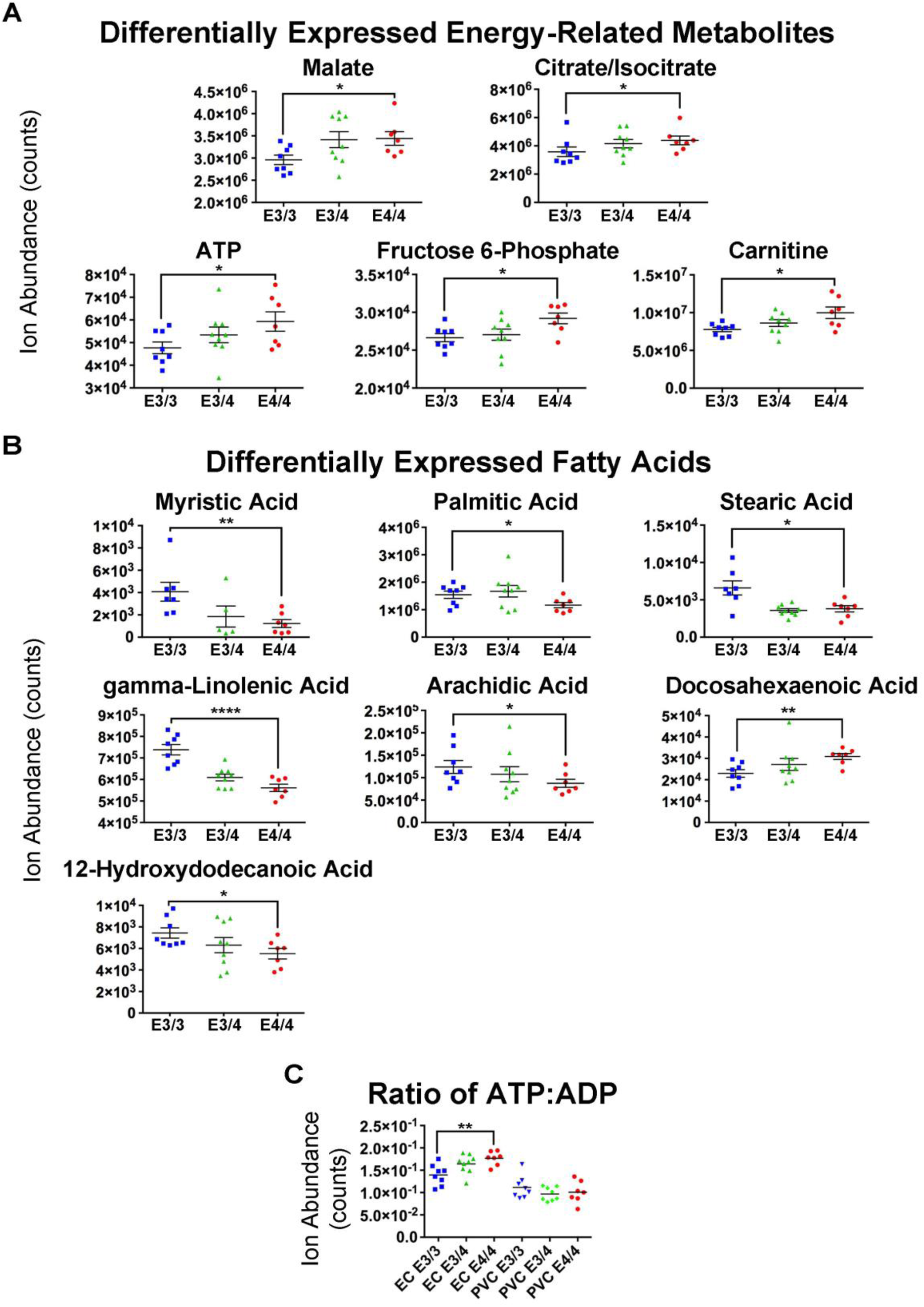
Metabolomics analysis reveals differential expression of energy-related metabolites and fatty acids in the EC of aged *APOE4* mice. An untargeted metabolomics analysis was performed in order to analyze the effects of differential *APOE* isoform expression on small-molecule metabolite levels in the EC and PVC of aged *APOE* mice (8 *APOE3/3*, 9 *APOE3/4* and 7 *APOE4/4* males). (A-C) Shown here are graphs of the bioenergetics-related metabolite groups that were shown to be differentially expressed in the EC of the *APOE4/4* vs. *APOE3/3* mice: (A) General energy-related metabolites and (B) Fatty acids. (C) We have also calculated the ratio of ATP:ADP in each genotype group from the EC and PVC of the aged *APOE* mice. (* denotes p < 0.05; ** denotes p < 0.01; **** denotes p < 0.0001)

Intriguingly, our metabolomics data also revealed an *APOE4*-associated decrease in the levels of numerous free fatty acids in aged *APOE4/4* vs. aged *APOE3/3* mice (Fig. 2B), which, unlike most of the previously mentioned energy-related metabolites, were similarly differentially altered in both the EC and the PVC. Although it is disputed whether or not free fatty acids can be utilized in the brain as an alternative energy source via β-oxidation (43-45), there is compelling evidence to suggest that it can (46-48). Thus, it is tempting to speculate that the decreased fatty acid levels observed in both the EC and PVC may be a result of increased β-oxidation in both of these brain regions, which may also be related to the observation that carnitine, a metabolite that is necessary for importing fatty acids into the mitochondria for energy utilization by the carnitine-palmitoyl transferase system, was the only energy-related metabolite in Fig. 2A whose levels were shown to be higher in both the EC and the PVC of the aged *APOE4/4* mice.

### Seahorse analysis reveals decreased mitochondrial respiration in the PVC, hippocampus and cortex of aged *APOE4* vs. *APOE3* mice, but not in the EC

In order to validate and expand upon the bioenergetic differences that we observed in our multi-omic analysis, we conducted a series of Seahorse experiments on mitochondria isolated from the EC and other brain regions of aged *APOE4/4* vs. *APOE3/3* mice. Utilizing the Seahorse XF24 platform (see methods section), we first measured the oxygen consumption rate (OCR; a measure of OxPhos efficiency) of mitochondria from the EC and PVC of 15-month-old *APOE* mice (4 *APOE3/3* and 4 *APOE4/4* males; samples pooled within each genotype and region). As shown in Supplementary Fig. S1, we observed a significant reduction in both the Complex I-mediated and Complex II-mediated OCR of mitochondria from the PVC of the *APOE4/4* vs. *APOE3/3* mice, but no difference in the OCR between genotypes in the mitochondria isolated from the EC region. In order to investigate whether this decreased OCR in the PVC was unique to the PVC or whether it represented a more global decrease in mitochondrial respiration within the aged *APOE4* brain, we performed an additional experiment measuring the OCR of mitochondria isolated from the EC, hippocampus (Hip) and cortex (Ctx) of 20-month-old *APOE* mice (2 *APOE3/3* and 2 *APOE4/4* males; samples pooled within each genotype and region). As shown in Fig. 3, we observed a significant reduction in the OCR of mitochondria from the Hip and the Ctx of *APOE4/4* vs. *APOE3/3* mice, with the most abundant differences observed in the Complex I-mediated OCR. However, in mitochondria from the EC of *APOE4/4* vs. *APOE3/3* mice, we did not observe any significant differences in Complex I-mediated basal mitochondrial respiration or in complex II-mediated OCR, and we observed significant increases in State 3 respiration in Complex I-mediated OCR in mitochondria from the EC (Fig. 3B). Furthermore, using the data generated from this Seahorse analysis, we determined the respiratory control ratio (RCR) (state3u/state4o), which is a measure of the coupling efficiency between OxPhos and ATP production (49, 50) (Fig. 3C). This analysis indicates that EC mitochondria from *APOE4* mice are highly coupled, resulting in increased production of ATP per unit of oxygen.

**Fig. 3.**
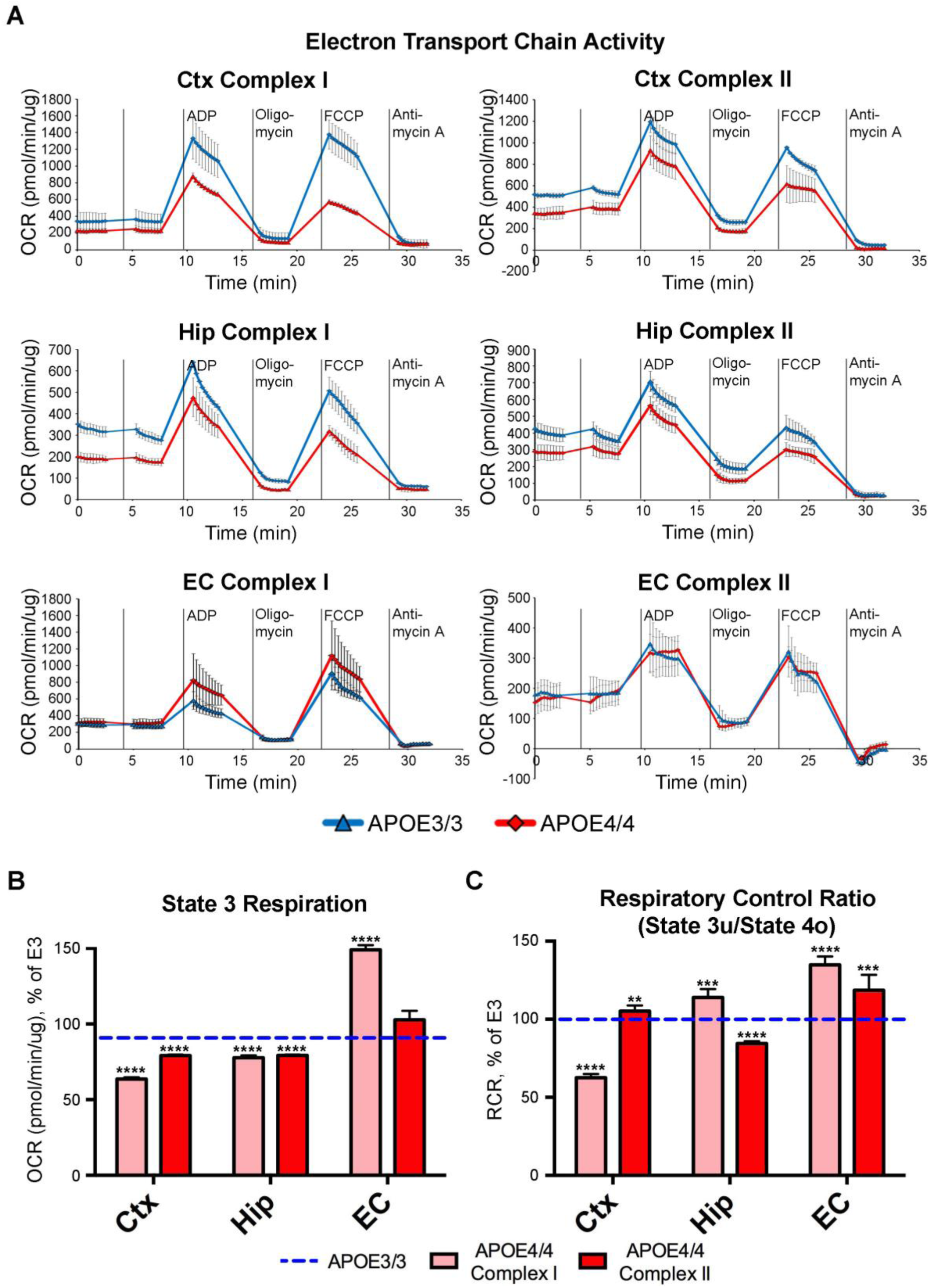
Seahorse analysis reveals decreased mitochondrial respiration in the cortex and hippocampus, but not in the EC, of aged *APOE4* mice. Seahorse analysis was performed in order to analyze the effects of differential *APOE* isoform expression on mitochondrial respiration in mitochondria that were purified from the cortex (Ctx), hippocampus (Hip) and EC of aged *APOE* mice (2 *APOE4/4* males, tissues pooled vs. 2 *APOE3/3* males, tissues pooled). (A) The complex I- and complex II-driven ETC activity from each region shows decreased mitochondrial respiration in the mitochondria purified from the cortex and hippocampus, but not the EC of the aged *APOE4/4* mice. (B-C) Bar graphs showing the average oxygen consumption rate (OCR) from (B) State 3 and for (C) the Respiration Control Ratio (RCR; state 3u/state 4o) in mitochondria purified from each region of the *APOE4/4* mice, as a percentage of the *APOE3/3* OCR from the equivalent tissues. The dotted blue line represents the normalized levels in the *APOE3/3* tissues. (* denotes p < 0.05; ** denotes p < 0.01; *** denotes *p* < 0.001; **** denotes p < 0.0001)

These observations suggest that, while *APOE4* expression leads to an overall reduction in mitochondrial function in the brains of aged *APOE4* mice, a result consistent with the *APOE4*-associated bioenergetic deficits observed using alternative approaches (24-29), mitochondria in the EC seem to undergo differential bioenergetic regulation, characterized by increased coupling efficiency. Importantly, these region-specific differences in the effects of differential *APOE* isoform expression on mitochondrial respiration did not appear to be the result of changes in mitochondrial mass, as the levels of Tom20 protein, as well as the levels of PGC1α RNA and the ratio of Mitochondrial:Nuclear DNA, were unchanged between *APOE* genotypes in the EC, Hip, and Ctx of 21-month-old *APOE* mice (Supplementary Fig. S2A, C & D). Differential *APOE* isoform expression also did not result in changes in the levels of specific ETC complex proteins in the EC, Hip or Ctx (Supplementary Fig. S2B).

### Computational analysis of untargeted metabolomics data reveals differential expression of ketones in the EC of aged *APOE4* mice

In order to investigate the source of these differential bioenergetic effects in the EC, we conducted a series of additional analyses on the EC of aged *APOE4/4* vs, *APOE3/3* mice. First, we utilized the PIUMet (Prize-collecting Steiner forest algorithm for Integrative Analysis of Untargeted Metabolomics) method (51) to predict identities of the untargeted metabolites that we uncovered in the EC of aged *APOE4/4* vs. *APOE3/3* during our original metabolomics analysis. PIUMet uses network optimization to analyze the data in the context of a vast database of protein-protein and protein-metabolite interactions, revealing both known and uncharacterized pathways that contain the putative metabolites and other molecules. PIUMet also allows for the incorporation of additional datasets, allowing us to utilize our targeted metabolomics results in the analysis.

We used PIUMet to analyze the 304 untargeted metabolite features whose levels were differentially altered between *APOE4/4* and *APOE3/3* in the EC of aged *APOE* mice (corrected p-value < 0.05). Among these features, 124 had matches in PIUMet’s underlying database of metabolites (Supplementary Table S5). PIUMet was able to infer the identity of 32 metabolite features, and revealed a network of protein-protein and protein-metabolite interactions associated with their dysregulation. The resulting network is enriched in 18 putative GO biological processes (Table 3) and 124 total metabolites (Supplementary Fig. S3). Among these, the most intriguing GO biological processes were fatty acid metabolic process, IMP metabolic process, ketone catabolic process, steroid metabolic process, and vitamin metabolic process. The most interesting putative identifications were acetone, adenylosuccinate, alpha-tocotrienol, and oleamide. Alpha-tocotrienol is a vitamin E metabolite, which bolsters the observation in the targeted metabolomics result that vitamins and vitamin-derivatives (including tocopherol, another vitamin E metabolite) were found to be increased in the EC and PVC of aged *APOE4/4* mice in our targeted metabolomics analysis (Tables 1 and 2). Similarly, the changes in the levels of adenylosuccinate support the idea that the purine nucleotide cycle is affected in *APOE4/4* EC, as this metabolite is downstream of IMP and upstream of succinoadenosine, both of which were identified in our targeted analysis (Table 1). Importantly, the identification of acetone and of the ketone catabolic process GO term may also represent a substantial finding. Ketone bodies (acetone, acetoacetone and β-hydroxybutyrate) are primarily generated in the liver from fatty acid breakdown via β-oxidation, and are used as a supplemental energy source when glucose levels are low. These ketone bodies readily cross the blood brain barrier and can act as a potent fuel for the brain (52-54). Alternatively, there is evidence that astrocytes also have the ability to produce ketones (55-57), both through the breakdown of fatty acids, as well as through the breakdown of ketogenic amino acids, most notably leucine (58, 59).

**Table 3.** GO biological processes observed in the PIUMet analysis of differentially expressed untargeted metabolites from the EC of *APOE4/4* vs. *APOE3/3* mice

| GO Term | Enrichment | p-value | FDR |
| --- | --- | --- | --- |
| carboxylic acid transport | 12.35 | 9.80E-20 | 1.43E-16 |
| carboxylic acid transmembrane transport | 22.34 | 1.36E-16 | 1.04E-13 |
| long-chain fatty acid metabolic process | 18.08 | 2.77E-13 | 1.50E-10 |
| fatty acid metabolic process | 7.8 | 1.45E-11 | 5.40E-09 |
| indolalkylamine metabolic process | 44.68 | 2.48E-09 | 6.68E-07 |
| disaccharide metabolic process | 63.29 | 6.92E-09 | 1.77E-06 |
| IMP metabolic process | 48.69 | 3.47E-08 | 7.43E-06 |
| steroid metabolic process | 6.87 | 1.78E-07 | 3.24E-05 |
| maltose metabolic process | 126.58 | 4.81E-07 | 8.06E-05 |
| kynurenine metabolic process | 50.63 | 7.51E-07 | 1.18E-04 |
| IMP biosynthetic process | 50.63 | 7.51E-07 | 1.19E-04 |
| tryptophan catabolic process | 50.63 | 7.51E-07 | 1.20E-04 |
| 'de novo' IMP biosynthetic process | 63.29 | 9.46E-06 | 1.13E-03 |
| ketone catabolic process |  | 3.90E-05 | 3.92E-03 |
| dopamine metabolic process | 20.25 | 4.12E-05 | 4.06E-03 |
| neurotransmitter catabolic process | 37.97 | 5.54E-05 | 5.25E-03 |
| keratan sulfate catabolic process | 31.65 | 1.00E-04 | 8.37E-03 |
| vitamin metabolic process | 6.38 | 3.60E-04 | 2.35E-02 |

### Proton nuclear magnetic resonance spectroscopy analysis reveals differential expression of phosphocreatine, glutamate and GABA in the EC of anesthetized aged *APOE4* mice

^1^H NMR spectroscopy provides *in vivo* measurements of brain metabolites and has been used in many studies of neurodegenerative disorders in both humans and rodents. This method allows for the simultaneous acquisition of several highly abundant metabolites in the brain, including neurotransmitters such as glutamate and gamma-aminobutyric acid (GABA), as well as other biological metabolites such as creatine, glutathione, and N-acetylaspartate (NAA). It also allows for the accurate quantification of individual metabolite concentrations. Since metabolomics and ^1^H NMR spectroscopy have been described as complimentary techniques, with advantages and disadvantages to both [see review by (60)], we chose to perform ^1^H NMR spectroscopy on the EC of 18-19 month-old *APOE* mice (7 *APOE3/3* and 7 *APOE4/4* males) in order to gain additional information about the metabolic changes occurring in the EC of aged *APOE* mice. While the EC is located in a very NMR unfriendly brain area because of its close proximity to sinus spaces, we were able to achieve full width half maximums of 8 – 15 Hz using a customized second and third order shim coil. Fig. 4A shows the position of the volume of interest (VOI) that was used to measure metabolite levels in the EC of these aged *APOE* mice. A representative spectrum from this region is shown in Fig. 4B, along with the spectral position of some of the major metabolites. To our knowledge, this is the first in vivo ^1^H NMR study of the EC in mice.

**Fig. 4.**
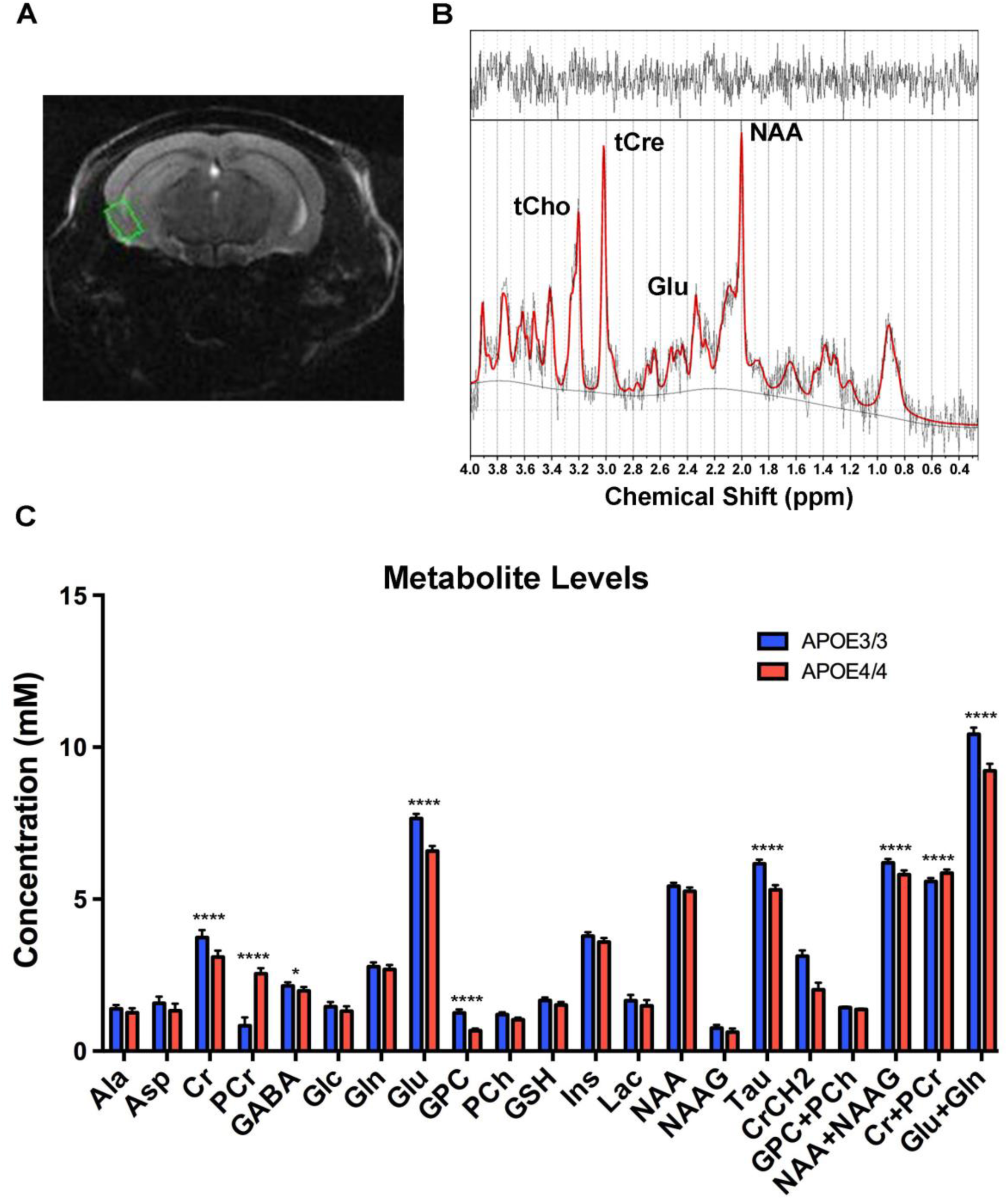
Proton nuclear magnetic resonance spectroscopy reveals differential expression of phosphocreatine, glutamate and GABA in the EC of aged *APOE4* mice. ^1^H NMR spectroscopy was performed in order to analyze the effects of differential *APOE* isoform expression on the levels of high-abundance metabolites in the EC of anesthetized *APOE* mice (7 *APOE4/4* males vs. 7 *APOE3/3* males). (A) A representative coronal anatomical image used in the analysis, with the volume of interest, highlighted in green, shown over the EC. (B) A typical spectrum from the EC showing the spectral position of some major metabolites: N-Acetylaspartic acid (NAA), glutamate (Glu), total creatine (tCr) and total choline (tCho). The raw spectrum is shown in black (with no smoothing), and the software fit of the model spectra shown in red. The top of the figure shows the difference between the raw spectrum and the fit, with a “good fit” showing random oscillations with little noise. The solid black line at bottom of the figure is the baseline. (C) The metabolite concentrations in the EC of aged *APOE3/3* and *APOE4/4* mice. The metabolites reported are: alanine (Ala), aspartate (Asp), creatine (Cr), phosphocreatine (PCr), gamma-aminobutyric acid (GABA), glucose (Glc), glutamine (Gln), glutamate (Glu), glycerophosphocholine (GPC), phosphocholine (PCh), glutathione (GSH), myo-inositol (Ins), lactate (Lac), N-Acetylaspartate (NAA), N-Acetylaspartateglutamate (NAAG), taurine (Tau), total choline (GPC and PCh), total NAA (NAA and NAAG), total creatine (Cr and PCr) and total glutamate and glutamine (Glu and Gln). (* denotes p < 0.05; ** denotes p < 0.01; *** denotes *p* < 0.001; **** denotes p < 0.0001)

As shown in Fig. 4C and Table 4, this ^1^H NMR study revealed significant differences in the levels of multiple metabolites in the EC of aged *APOE4/4* vs. *APOE3/3* mice, including decreased levels of creatine (Cre), glutamate (Glu), GABA, and taurine (Tau) in the EC of aged *APOE4/4* mice, as compared to that of aged *APOE3/3* mice. Perhaps the most interesting observation, however, was the large increase in the levels of phosphocreatine (PCr) and the ratio of PCr:Cr observed in the *APOE4* EC. Creatine is mainly phosphorylated in mitochondria, where creatine kinase uses the newly synthesized ATP to produce PCr and ADP (61). This PCr is then shuttled to the cytosol where it acts as a rapidly mobilized reserve of phosphate groups that can be used to replenish ATP levels, whereas the increased levels of ADP in the mitochondria will stimulate oxygen uptake and oxidative metabolism, especially during enhanced neuronal activity (62). Thus, the observation that PCr levels are increased in aged *APOE4/4* mice supports and expands upon the bioenergetics observations from our transcriptomics, metabolomics and Seahorse analyses. In addition to these changes in PCr levels, the observation that glutamate and GABA are both decreased in the EC of *APOE4/4* mice may also be related to these bioenergetic differences between *APOE* genotypes and brain regions. Although glutamate and GABA are mostly known for their roles as neurotransmitters in excitatory and inhibitory neurons, respectively, they can also be utilized as energy metabolites (63-66).

**Table 4.** Metabolite concentrations in the EC of *APOE3/3* and *APOE4/4* mice, as observed by

| **Metabolite** | ***APOE3/3***<br>**concentration (mM)** | ***APOE4/4***<br>**concentration (mM)** | **p-value** |
| --- | --- | --- | --- |
| Creatine (Cr) | 3.73 +/- 0.25 | 3.09 +/- 0.21 | 3.04E-04 |
| Phosphocreatine (PCr) | 0.84 +/- 0.27 | 2.55 +/- 0.18 | 1.00E-05 |
| Gamma-aminobutyrate (GABA) | 2.14 +/- 0.12 | 1.98 +/- 0.13 | 0.036 |
| Glutamate (Glu) | 7.65 +/- 0.15 | 6.58 +/- 0.17 | 5.30E-05 |
| Glycerophosphocholine (GPC) | 1.25 +/- 0.12 | 0.67 +/- 0.07 | 2.90E-05 |
| Taurine (Tau) | 6.16 +/- 0.14 | 5.31 +/- 0.15 | 1.59E-05 |
| Cr+PCr | 5.58 +/- 0.11 | 5.85 +/- 0.12 | 8.90E-04 |
| Glu+Gln | 10.43 +/- 0.21 | 9.22 +/- 0.23 | 2.30E-05 |
| NAA+NAAG | 6.19 +/-0.14 | 5.81 +/- 0.12 | 8.90E-04 |

**Table 5.** Differentially expressed targeted lipids from the EC of APOE4/4 vs. APOE3/3 mice

| Metabolite | Regulation<br>in E4/4 | Fold<br>Change | p-value |
| --- | --- | --- | --- |
| **Cholesteryl esters** |  |  |  |
| CE 18:0 | down | 1.44 | 0.039 |
| **Diacylglycerols** |  |  |  |
| DAG 28:0/14:0 | down | 1.33 | 0.003 |
| DAG 30:0/14:0 | down | 1.50 | 0.002 |
| DAG 32:0/16:0 | down | 1.30 | 0.006 |
| DAG 32:1/16:0 | down | 1.45 | 0.001 |
| DAG 32:2/16:1 | down | 1.38 | 0.031 |
| DAG 34:1/16:0 | down | 1.28 | 0.011 |
| DAG 34:2/16:0 | down | 1.38 | 0.016 |
| DAG 38:3/18:0 | down | 1.24 | 0.032 |
| DAG 38:4/18:1 | down | 1.32 | 0.022 |
| DAG 40:6/18:0 | down | 1.19 | 0.013 |
| **Ceramides** |  |  |  |
| Cer 18:1 | up | 1.60 | 2.00E-04 |
| **Sphingomyelins** |  |  |  |
| SM 18:0 | down | 1.09 | 0.045 |
| **Hexosylceramides**<br>**(glucosyl- and galactosyl-)** |  |  |  |
| HexCer 16:0 | up | 2.45 | 1.19E-05 |
| HexCer 16:1 | up | 1.50 | 0.001 |
| HexCer 18:0 | up | 1.43 | 0.020 |
| HexCer 20:0 | up | 1.47 | 0.027 |
| HexCer 26:0 | up | 2.08 | 0.049 |
| **Sulfatides** |  |  |  |
| Sulf(2OH) 18:0 | up | 1.31 | 0.028 |
| **Lactosylceramides** |  |  |  |
| LacCer 16:0 | up | 1.81 | 0.005 |
| **Phosphatidic acids** |  |  |  |
| PA 40:6 | up | 1.18 | 0.044 |
| **Phosphatidylcholines** |  |  |  |
| PC 30:0 | down | 1.14 | 0.045 |
| PC 36:0 | up | 1.11 | 0.006 |
| PC 42:7 | up | 1.28 | 0.020 |
| **Phosphatidylinositols** |  |  |  |
| PI 38:3 | down | 1.39 | 0.003 |
| PI 38:4 | down | 1.32 | 0.033 |
| **Phosphatidylserines** |  |  |  |
| PS 42:4 | up | 1.33 | 0.022 |
| PS 42:5 | up | 1.29 | 0.031 |
| **Lyso ether**<br>**phosphatidylcholines** |  |  |  |
| LPCe 18:0 | up | 1.14 | 0.010 |
| **Lyso phosphatidylinositols** |  |  |  |
| LPI 16:0 | down | 1.15 | 0.002 |
| **Bis(monoacylglycero)**<br>**phosphates, LBPA** |  |  |  |
| BMP 32:0 | up | 1.41 | 0.003 |
| BMP 34:0 | up | 1.39 | 0.014 |
| BMP 34:1 | up | 1.37 | 0.013 |
| BMP 36:0 | up | 1.75 | 0.011 |
| BMP 36:1 | up | 1.44 | 0.014 |
| BMP 38:4 | up | 1.95 | 0.011 |

**Table 6.** Differentially expressed targeted lipids from the PVC of APOE4/4 vs. APOE3/3 mice

| Metabolite | Regulation<br>in E4/4 | Fold<br>Change | p-value |
| --- | --- | --- | --- |
| **Diacylglycerols** |  |  |  |
| DAG 38:3/18:1 | down | 1.47 | 0.023 |
| DAG 40:5/18:0 | down | 1.41 | 0.014 |
| **Ceramides** |  |  |  |
| Cer 18:1 | down | 1.30 | 0.017 |
| **Sphingomyelins** |  |  |  |
| SM 16:0 | down | 1.30 | 0.015 |
| SM 20:0 | down | 1.27 | 0.011 |
| **Dihydrosphingomyelins** |  |  |  |
| dhSM 16:0 | down | 1.64 | 0.020 |
| **Hexosylceramides**<br>**(glucosyl- and galactosyl-)** |  |  |  |
| HexCer 16:1 | down | 1.56 | 3.16E-04 |
| **Ether**<br>**phosphatidylcholines** |  |  |  |
| PCe 36:0 | down | 1.26 | 0.043 |
| PCe 40:6 | down | 1.29 | 0.005 |
| **Lyso ether**<br>**phosphatidylcholines** |  |  |  |
| LPCe 16:0 | down | 1.53 | 0.007 |
| **Bis(monoacylglycero)**<br>**phosphates, LBPA** |  |  |  |
| BMP 32:0 | up | 1.66 | 0.041 |
| BMP 32:1 | up | 1.54 | 0.031 |
| BMP 34:0 | up | 1.62 | 0.035 |
| BMP 34:2 | up | 1.36 | 0.007 |
| **N-acyl**<br>**phosphatidylserines** |  |  |  |
| NAPS 38:4 | up | 2.09 | 0.024 |

Taken together, our data suggest that, contrary to other brain region such as the cortex and hippocampus, the EC of aged *APOE4/4* vs. *APOE3/3* mice exhibit an increased rate of oxidative metabolism and ATP turnover, which may play a causative role in the pathogenesis of AD.

## DISCUSSION

Possession of the *APOE4* allele greatly increases an individual’s risk of developing AD. Although numerous theories have been proposed, the precise cause of this association remains unknown. In order to further investigate the effects of *APOE4* gene expression in AD-relevant brain regions, we initially chose to perform a multi-omic analysis on RNA and small-molecules extracted from an AD-vulnerable brain region (the EC) vs. an AD-resistant brain region (the PVC) of aged *APOE* targeted replacement mice. In addition to our recently published reports describing *APOE4*’s impact on endosomal-lysosomal dysregulation (41) and neuronal hyperactivity (42), these studies also revealed *APOE4*-associated alterations in the levels of numerous genes and small-molecules related to energy metabolism (Fig. 1 and 2). Follow-up experiments utilizing the Seahorse platform revealed *APOE4*-associated deficits in mitochondrial respiration in several brain regions, including the PVC, the Hip, and the Ctx as a whole, which is consistent with previous reports showing that *APOE4* expression has numerous deleterious effects on the brain’s bioenergetic capabilities (24-29). Intriguingly, however, this *APOE4*-associated decrease in mitochondrial respiration was not observed in the EC, with Complex-1 mediated respiration actually increasing in the EC of 20-month-old *APOE4* mice. (Fig. 3). Additional studies, including proton nuclear magnetic resonance (^1^H NMR) spectroscopy and a bioinformatics analysis of our untargeted metabolomics data provided further support and elucidation of this region-specific bioenergetic regulation by *APOE4*, including the observations of increased PCr levels and decreased glutamate and GABA levels (Fig. 4), as well as an upregulation of the ketone pathway (Table 3), in the EC of aged *APOE4* mice.

Bioenergetic regulation in the brain is a complex process involving multiple interconnected pathways and cell-types, which makes it difficult to decipher the mechanism or impact of these region-specific bioenergetic effects of *APOE4* expression. It is important to note that the decline in mitochondrial functionality that we observed in the cortex and hippocampus, while significant, is likely below the pathological threshold (67). Mitochondria are part of a multifaceted network of metabolic pathways that are modulated by the cell to compensate for any reductions in respiratory activity and ATP production (68, 69). Moreover, mitochondria have a “reserve respiratory capacity” that allows them to increase their oxidative capacity and ATP production depending on the cell’s demand (70). Furthermore, mitochondrial adaptability is tissue and cell-type specific, depending on the different compensatory mechanisms and the availability of substrates (67, 71)

The most intriguing finding from our study, however, is that, within the brain, mitochondria appear to have *regional* differences in their response to metabolic stressors, at least in this case of *APOE4* expression in aged *APOE* mice. In both the EC and the PVC, for example, it appears that there are attempts to use alternative fuel sources to compensate for *APOE4*’s effects on mitochondrial function. This is best represented by the decreased fatty acid levels and increased carnitine levels in the metabolomics data, perhaps due to increased efforts to supply the TCA cycle with additional acetyl-CoA molecules via β-oxidation of fatty acids. However, the data from our ^1^H NMR spectroscopy study, as well as further mining of our transcriptomics and metabolomics data, suggest that the mitochondria in the EC of aged *APOE4* mice have additional mechanisms that allow them to counteract the *APOE4*-associated reductions in mitochondrial respiration. Specifically, we hypothesize that mitochondria in the EC of aged *APOE4* mice are distinctively able to increase their respiratory capacity by increasing the coupling efficiency between oxygen consumption and ATP production.

This hypothesis is supported by our Seahorse data, where mitochondria in the EC of aged *APOE4* mice show increased ADP-stimulated oxygen consumption and increased maximal respiration rate, without changes in proton leak (Fig. 3 and Supplemental Fig. S1) or mitochondrial mass (Supplementary Fig. S2). In addition, we also observed that the RCR is increased in the EC of aged *APOE4/4* vs. *APOE3/3* mice (Fig. 3C and Supplementary Fig. S1C), further confirming that ADP-driven mitochondrial respiration in the aged *APOE4* EC is better coupled to ATP production than in the Hip and the Ctx. Interestingly, the reductions in mitochondrial respiration observed in the Hip and Ctx of aged *APOE4/4* vs. *APOE3/3* mice by Seahorse analysis appears to be more robust in complex I-mediated respiration compared to complex II-mediated respiration, whereas in the EC, complex I-mediated respiration in aged *APOE4* begins to outpace complex I-mediated respiration in *APOE3* mice with increasing age (Fig. 3 and Supplemental Fig. S1). Since complex I-mediated mitochondrial respiration is driven by pyruvate, this suggests that the *APOE4*-associated bioenergetic deficits observed in the hippocampus and cortex of aged *APOE4* mice are primarily related to decreased aerobic respiration that occurs downstream of glycolysis, whereas in the EC we observe an amplification of this process.

While the reason for the increased efficiency in OxPhos in the EC of aged *APOE4* mice requires further investigation, we note that the expression of some mitochondrial transporter genes is specifically upregulated in mitochondria from this brain region, as detected in the transcriptomics data (Supplementary tables S1 and S2). In order to ensure a sufficient rate of solute flux into mitochondria to fuel metabolic pathways, numerous mitochondrial transporters link biochemical pathways in the cytosol and mitochondria (72). Regulation of these transporters is an essential part of the overall regulation of cellular metabolism, and their expression is highly variable depending on the tissues and the metabolic conditions. Intriguingly, the expression of the adenine nucleotide translocase (ANT) genes, ANT1 (*Slc24a4*) and ANT2 (*Slc24a5*), were significantly increased in the EC of aged *APOE3/4* vs. *APOE3/3* mice. ANT participates in ATP-for-ADP exchange through the inner mitochondrial membrane, which supplies the cytoplasm with newly synthesized ATP from OxPhos. Interestingly, ANT can also interact with creatine kinase (CK), gaining preferential access to ATP, which is needed to synthesize PCr in cells with high energetic needs (73). Similarly, the expression of the genes for the voltage dependent-anion channel (VDAC1; Vdac1), one of the main gatekeepers for the regulation of the crosstalk between mitochondria and cytosol (72), is also increased in the EC of aged *APOE3/4* mice, as is the gene for the mitochondrial phosphate transport protein (PTP; *Slc25a3*), which transports inorganic phosphate (P_i_) to the mitochondrial matrix in order to supply the P_i_ required for ADP phosphorylation into ATP (72).

Another important mitochondrial transporter that ensures the balanced cooperation between OxPhos and glycolysis (and to a lesser extent the pentose phosphate pathway) in order to maintain ATP levels is the malate-aspartate shuttle. Briefly, in order for the two NADH molecules generated during glycolysis to be translocated into mitochondria, malate must be produced in the cytosol via conversion of oxaloacetate (OAA) by malate dehydrogenase (MDH1), for which NADH is a necessary cofactor, and then malate is converted back to OAA by MDH1 inside mitochondria (converting NAD+ back to NADH in the process). Since this newly formed OAA cannot be transported back to the cytosol, OAA must first be converted to aspartate by glutamate oxaloacetate transaminase 1 (GOT1; *Slc25a22*). But this requires the donation of an amino radical from glutamate. Thus, glutamate is an essential component for maximizing the number of ATPs produced in glycolysis per molecule of glucose metabolized. Interestingly, our transcriptomic data reveals that the expression of both Slc25a22 (GOT1) and *Mdh1* are upregulated in the EC of aged *APOE3/4* vs. *APOE3/3*, suggesting that activity of the malate-aspartate shuttle is increased in the EC of these mice.

A summary of all these bioenergetics-related changes that we observe in the EC of aged *APOE4* mice is depicted in Fig. 5. Taken together, the observed differences on mitochondrial functionality among the brain regions studied suggests that the EC possesses a different subset of compensatory mechanisms in order to improve OxPhos efficiency under metabolic stress. While this apparent compensation of EC mitochondria may be regarded as a positive attribute, it is plausible that it may have negative consequences in the long run. For example, it has been shown that, over time, elevated mitochondrial respiratory activity and increased coupling efficiency can result in an excess of free radical production and resulting oxidative damage within cells (74).

**Fig. 5.**
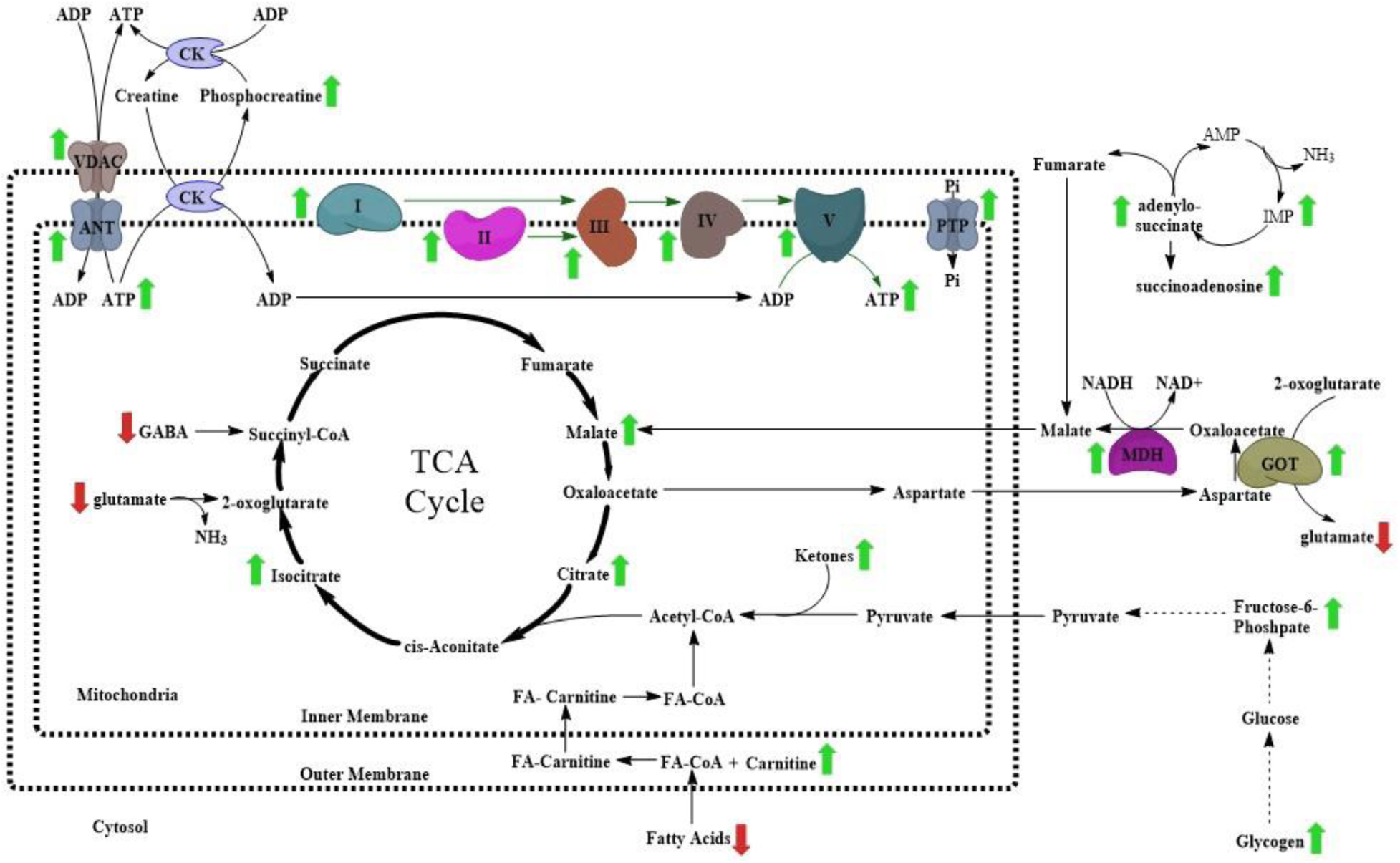
Identification of differentially expressed bioenergetic genes and metabolites in the EC of aged *APOE4* mice. A schematic of the bioenergetics genes and metabolites that have been predicted to be altered by differential *APOE* isoform expression in this study, as depicted in a simplified manner with respect to their relationships to each other and their location inside or outside of mitochondria. Genes or metabolites that were putatively identified as increased in aged *APOE4* mice, as compared to aged *APOE3* mice, are marked with a green arrow, and those that are decreased are marked with a red arrow.

More work will be necessary in order to further elucidate this phenomenon, including an understanding of the specific contributions of different cell-types such as neurons, astrocytes and microglia. It will also be important to investigate whether the other phenotypes that we have observed in the EC of these aged *APOE4* mice, such as endosomal-lysosomal dysregulation and neuronal hyperactivity, are related to the bioenergetic observations described here. For example, Zhao et al. recently reported that *APOE4* expression causes insulin receptors to be inefficiently recycled via the endosomal-lysosomal system, potentially causing the insulin resistance that the authors observed in the brains of aged *APOE4* mice (25). As the endosomal-lysosomal dysregulation that we observed in the brains of aged *APOE4* mice was not restricted to the EC (41), this may play a role in the *APOE4*-associated bioenergetic deficits that we observed throughout the brains of these mice. On the other hand, the neuronal hyperactivity that we observed did appear to be primarily localized to the EC within the hippocampal formation (42). Intriguingly, neuronal activity and glucose utilization have previously been shown to exist in a near stoichiometric relationship (75), suggesting that the bioenergetic counterbalancing that we are observing in the EC of aged *APOE4* mice may be directly correlated to the observed neuronal hyperactivity in these mice (42).

Finally, it will be important to determine whether these bioenergetic differences in the EC are involved in the increased risk of AD among *APOE4* carriers. The EC is a unique brain region in many ways. It is vital for learning and memory (especially spatial memory), acting as a bridge between the hippocampus and the neocortex (76, 77). As such, it is known to have higher energy demands and to be more metabolically active that many other brain regions (78). In AD, the EC is one of the first regions to develop tauopathy, which eventually spreads into the hippocampus and other cortical regions as the disease progresses (33). It is possible, therefore, that due to its unique bioenergetic needs, the EC may possess additional mechanisms for increasing its bioenergetic capacity under conditions of metabolic stress such as *APOE4* expression. However, perhaps in the process of inducing these mechanisms, other negative consequences may result, such as increased ROS production or increased tau hyperphosphorylation or increased neuronal hyperactivity, such that it results in an earlier onset of AD.

## METHODS

### Mice

Human APOE targeted replacement mice were first developed by Sullivan et al. (79, 80) and were acquired from Taconic Biosciences. All mice used in this study were treated in accordance with the National Institutes of Health Guide for the Care and Use of Laboratory Animals and approved by the Columbia University Medical Center and SUNY Medical Center Institutional Animal Care and Use Committee (IACUC).

### RNA-Sequencing

Male mice expressing human *APOE3/3* (10 mice) or *APOE3/4* (19 mice) were aged to 14-15 months, at which point they were sacrificed by cervical dislocation, and brain tissues containing the EC and PVC were dissected and snap frozen on dry-ice. Brain tissues were stored in RNase-free eppendorf tubes at −80°C prior to extraction. Total RNA was extracted from frozen tissues by homogenizing each tissue sample using a battery-operated pestle mixer (Argos Technologies, Vernon Hills, IL) in 1 ml of TRIzol reagent according to the manufacturer’s protocol (Life Technologies, Carlsbad, CA). RNA concentration was measured using a nanodrop 1000 (Thermo Fisher Scientific, Waltham, MA), and RNA integrity (RIN) was assessed using an Agilent 2100 Bioanalyzer (Agilent Technologies, Santa Clara, CA). All RNA samples possessed a RIN of 8 or higher. RNA was stored at −80°C prior to use. Starting with 2 μg of total RNA per sample, Poly(A)+ mRNA was purified, fragmented and then converted into cDNA using the TruSeq RNA Sample Prep Kit v2 (Illumina cat# RS-122-2001) as per the manufacturer’s protocol (Illumina, San Diego, CA). For RNA-Sequencing of the cDNA, we hybridized 5 pM of each library to a flow cell, with a single lane for each sample, and we used an Illumina cluster station for cluster generation. We then generated 149bp single end sequences using an Illumina HiSeq 2000 sequencer. For analysis, we used the standard Illumina pipeline with default options to analyze the images and extract base calls in order to generate fastq files. We then aligned the fastq files to the mm9 mouse reference genome using Tophat (v2.0.6) and Bowtie (v2.0.2.0). In order to annotate and quantify the reads to specific genes, we used the Python module HT-SEQ with a modified NCBIM37.61 (containing only protein coding genes) gtf to provide reference boundaries. We used the R/Bioconductor package DESeq2 (v1.10.1) for comparison of aligned reads across the samples. We conducted a variance stabilizing transformation on the aligned and aggregated counts, and then the Poisson distributions of normalized counts for each transcript were compared across *APOE3/4* vs. *APOE3/3* groups using a negative binomial test. We corrected for multiple testing using the Benjamini-Hochberg procedure and selected all genes that possessed a corrected p-value of less than 0.05. Finally, a heat map based on sample distance and a volcano plot based on fold change and adjusted p-values were generated using the R/Bioconductor package ggplot2 (v2.0.0). For pathway analysis, enriched KEGG pathways were identified using the ClueGo application (version 2.1.7) in Cytoscape (version 3.2.1). Briefly, all differentially expressed EC genes from the RNA-Seq analysis were entered into the application and searched for significantly enriched KEGG pathways possessing a Benjamini-Hochberg adjusted p-value of less than 0.05.

### Metabolomic Analysis

Male mice expressing human *APOE3/3* (8 mice), *APOE3/4* (9 mice), or *APOE4/4* (7 mice) were aged to 14-15 months, at which point they were sacrificed by cervical dislocation to maintain the brain environment, and individual brain regions were immediately removed and snap-frozen on dry ice. Tissues were stored at −80°C for prior to extraction. Metabolite extraction was performed using a methyl tert-butyl ether (MTBE)/methanol extraction protocol modified from previous reports (39, 40). Briefly, individual EC or PVC tissues were homogenized in 400 μl of ice-cold methanol using a bead mill homogenizer (TissueLyser II, Qiagen) at 25 beats/sec, 2x for 45 sec each. Following homogenization, samples were incubated in 1200 μl of MTBE for 1 hr at room temperature to separate organic-soluble lipids from aqueous-soluble lipids and other small-molecules. Finally, 360 μl of ultrapure water was added (for a final ratio of 3:1:0.9 MTBE:methanol:water) to resolve the two liquid phases, and each samples were centrifuged at 10,000 × g for 10 min. The lower aqueous phase and the upper organic phase was collected from each sample and stored in separate tubes, and the remaining protein pellets were resuspended in 25 mM ammonium bicarbonate, pH 8, with 2.5% SDS. A BCA protein assay was performed on each protein fraction, and both the aqueous phase and organic phase were normalized to their protein concentration equivalent with 50% and 100% methanol respectively. All samples were then stored at −80°C prior to analysis. Metabolite profiling was performed using an Agilent Model 1200 liquid chromatography (LC) system coupled to an Agilent Model 6230 time-of-flight (TOF) mass analyzer as described previously (81). Metabolite separation was accomplished using aqueous neutral phase (ANP) gradient chromatography on a Diamond Hydride column (Microsolv), with mobile phase buffer A consisting of 50% isopropanol and 0.025% acetic acid, and mobile phase buffer B consisting of 90% acetonitrile and 5 mM ammonium acetate. Each aqueous phase sample was analyzed in both positive and negative ion detection mode for comprehensive coverage of metabolites with a mass of 50-1000 Da. Prior to analysis, it was observed that one APOE3/4 PVC sample possessed a chromatographic error and was removed from all future analysis.

Following MS analysis, the raw data was analyzed using Agilent Profinder (version B.06.00) and Mass Profiler Professional (version 12.0) software. Briefly, Profinder performs batch recursive feature extraction, which consists of a first-pass identification of features (ions) that possess a common elution profile (e.g. identical mass-to-charge [m/z] ratios and retention times), followed by a recursive analysis to re-mine each sample for the presence of features that were identified in at least one sample. Feature extraction was limited to the top 2000 metabolites per ion detection mode, with each metabolite possessing a minimum abundance of 600 ion counts. Each feature was also manually validated following extraction (with investigators blinded to genotype assignments during validation), and re-integration was performed when necessary. In addition to this untargeted feature extraction, we also used ProFinder to perform a targeted feature extraction, matching features against a proprietary database of 626 biologically-relevant metabolites whose mass and retention times were previously determined using pure chemical standards. For this analysis, we allowed for a maximum mass deviation of 10 mDa and a retention time deviation of 1 min, and identified features were manually validated following extraction. For both untargeted and targeted feature extraction, Profinder analysis was followed up by multivariate and differential expression analysis using Mass Profiler Professional. Following differential expression analysis, metabolites identified as differentially expressed were further validated for chromatographic and biological accuracy, and non-validated metabolites were removed.

### Seahorse Analysis

In order to investigate the effect of differential apoE isoform expression on electron transport chain activity, mitochondria were isolated from tissues collected from 12-17 month old apoE3/3 or E4/4 mice (10 mice per group, separated into three experiments), and the oxygen consumption rates (OCR) were measured in a complex I or complex II-mediated fashion, as previously described (49). Briefly, each mouse was sacrificed by cervical dislocation, and the different brain areas from each brain hemisphere were immediately removed and placed into ice-cold PBS containing protease inhibitors. For each experiment, tissues were pooled from 3-4 mice per apoE group. Following tissue collection, samples were homogenized in ∼10 volumes of isolation buffer (70 mM sucrose, 210 mM mannitol, 5 mM HEPES, 1 mM EGTA and 0.5% fatty acid-free BSA, pH 7.2) and rinsed 3 times. Tissues were then homogenized using a Teflon glass homogenizer with 2-3 strokes. Homogenized samples were then centrifuged at 800 × g for 10 min at 4°C. Following centrifugation, fat/lipid was aspirated off, and the remaining supernatant was decanted through 2 layers of cheesecloth into a separate tube and centrifuged at 8000 × g for 10 min at 4°C. After removal of the supernatant, the pellet was resuspended in mitochondrial isolation buffer, and the centrifugation was repeated. The resulting pellet was resuspended in mitochondrial isolation buffer, and total protein concentrations were determined using Bradford Assay reagent (Bio-Rad).

For complex I experiments, 8 μg of protein was added to each well and for complex II analysis 6 μg per well.

To prepare samples for OCR measurements, samples were prepared in mitochondrial assay buffer (70 mM sucrose, 220 mM mannitol, 10 mM KH2PO4, 5 mM MgCl2, 2 mM HEPES, 1 mM EGTA and 0.2% fatty acid-free BSA, pH 7.2) plus substrate (pyruvate and malate for complex I-mediated ETC activity assay, or succinate and rotenone for complex II-mediated ETC activity assay).

Respirometry was performed using the XF24e Extracellular Flux Analyzer (Seahorse Bioscience). Oxygen consumption was measured in basal conditions (Seahorse media with 25 mM glucose and 2 mM pyruvate) and after the sequential addition of 1 μM oligomycin (Complex V inhibitor), 0.75 μM FCCP (uncoupler) and 1 μM rotenone/1 μM antimycin A (Complex I and Complex III inhibitors respectively). All results are averages of five or more biological replicates. Every biological replicate consisted of three technical replicates.

OCR data was generated by the Seahorse XF24 1.5.0.69 software package and displayed in point-to-point mode. Final calculations of total OCR were performed by normalizing apoE4/4 values to apoE3/3 for each of the three experiments and then combining the data in Prism.

### PIUMet Analysis

We used the previously published PIUMet algorithm (51) to analyze the untargeted metabolomics data from APOE4/4and APOE3/3 in the EC of aged APOE mice. PIUMet discovers dysregulated molecular pathways and components associated with changes in untargeted metabolomic data by analyzing a network built from curated protein-protein and protein-metabolite interactions (PPMI). PIUMet represents each metabolite peak as a node in this network and connects it to metabolites with masses that correspond to the m/z values of the peak. Using the prize-collecting Steiner tree algorithm, it searches the PPMI interactome to find an optimum subnetwork that connects the input metabolite peaks via their putative identities and other metabolites and proteins that were not detected in the experiments. The Gene Ontology (GO) analysis of the resulting network further reveals associated molecular pathways with the *APOE4* phenotype. We selected peaks 304 untargeted metabolite peaks or features that were significantly altered between APOE4/4and APOE3/3 models (corrected p-value <0.05). We then assigned a prize to each input data point to show the significance of their alterations, as –log (P values) of the significance of changes between two phenotypic conditions. PIUMet accepts several parameters that regulate the size of the resulting networks, including w and beta. While w tunes the number of trees in the resulting network, beta tunes the number of input nodes that are included in the output. Another parameter, mu, controls the bias toward high degree nodes. Higher values of mu results in a lower number of high degree nodes in the resulting networks, and thus less bias to highly studied molecules or ubiquitous interactions such as those with ions. We determined the optimum parameters (w=6.0, beta=0.5, and mu=0.0005) based on their effects on the size of the resulting networks. We further generated 100 networks by adding random noise to the underlying database, and calculated robustness scores for each node. Nodes with robustness scores less than 60% were removed from the results. Additionally, for each node we calculated a specificity score, based on the number of their presence in the 100 random networks obtained from a set of mock metabolite peaks randomly selected from the input feature list that mimic real input. A score of 100 indicates that the node did not show in any of the randomly generated networks, while a zero score shows the node appears in all the randomly generated networks. 79% of the resulting nodes have specificity score of over 60%.

### ^**1**^H NMR Spectroscopy

All animal procedures for in vivo proton MR spectroscopy were performed following the National Institutes of Health guidelines with approval from the Institutional Animal Care and Use Committee at the Nathan S. Kline Institute for Psychiatric Research. All animals were anesthetized using an isoflurane vaporizer set at the following percentages: 3% for induction, 2% during pilot scanning and 1.5% during data acquisition. An animal monitoring unit (model 1025, SA Instruments, Stony Brook, NY, USA) was used to record respiration and rectal temperature. Respiration was measured with a pressure transducer placed under the abdomen just below the ribcage. Body temperature was maintained using forced warm air, controlled by a feedback circuit between the heater and thermistor. After induction, the animals were placed on a holder and restrained using a bite bar and ear bars placed half way into the auditory canal. Oxygen was used as the carrier gas delivered to a cone positioned before the bite bar, where gases mixed with air and passed over the rodent’s nose. All animals were maintained at 37.0 ± 0.2 °C. All data were obtained on a 7.0 T Agilent (Santa Clara, CA, USA) 40 cm bore system. The gradient coil insert had an internal diameter of 12 cm with a maximum gradient strength of 600 mT/m and minimum rise time of 200 μs, with customized second and third order shim coils. A Rapid (Rimpar, Germany) volume transmit coil (72mm ID) and a two-channel receive-only surface coil was used for RF transmission and reception, respectively.

The shim settings for the selected volume of interest (VOI) were automatically adjusted using FASTMAP^1^ (Fast, Automatic Shimming Technique by Mapping Along Projections), a high order shim method, which samples the magnetic field along a group of radial columns which focus on the center of a localized voxel. It is a method for optimizing the field homogeneity in a cubical local region.

The water signal was suppressed using variable power RF pulses with optimized relaxation delays (VAPOR)^2^. The spectral acquisition consisted of a short echo time Point Resolved Spectroscopy (PRESS)^3^ sequence with the following parameters: repetition time = 4s, echo time = 7.5 ms, number of averages = 512 acquired in blocks of 128, number of points = 2048 and bandwidth of acquisition = 5 kHz, total acquisition time = 34 minutes. Outer volume suppression was also used to minimize signal contamination by extra cranial muscle and lipids. The VOI size was 3.3 µl (2.0 × 1.3 × 1.3 mm^3^) placed in the entorhinal cortex (EC). The target VOI is depicted in Fig. 4 overlaid on a schematic of the coronal brain slice at approximately – 2.92 mm relative to Bregma. An anatomical T_2_ weighted pilot scan was used to position the VOI (coronal). These scans were acquired with a fast spin echo sequence with the following parameters: field of view = 20 mm with 256 × 256 matrix size, slice thickness = 0.5 mm, number of slices = 11, repetition time = 4s, echo train length = 8, echo spacing = 15 ms, effective echo time = 60 ms, number of averages = 8, total acquisition time = 8 minutes 40 s. All data were processed using the LCModel software developed by Provencher ^4^. This software calculates the best fit to the acquired data of a linear combination of model spectra acquired from *in vitro* solutions of all the brain metabolites of interest. This basis set of the model spectra has the same echo time, sequence acquisition and field strength as the acquired data of the study. The LCModel software outputs the estimated concentration along with estimated standard deviations (Cramer-Rao lower bounds) expressed in percent of the estimated concentration, which can be used as a quantitative measure of reliability. To improve statistical significance, Miller et al. recently demonstrated the use of weighted averaging in NMR spectroscopic studies (82). In this current study, we also used weighted averaging to calculate the standard unequal variance t-test (Welch’s t-test) as outlined by Miller ^(82)^ and in the reference manual of the LCModel software (83). All data used in the final analysis had Cramer-Rao lower bounds of 20% or less. The metabolites measured were: alanine(Ala), aspartate (Asp), creatine (Cr), phosphocreatine (PCr), γ-Aminobutyric Acid (GABA), glucose (Glc), glutamine (Gln), glutamate (Glu), glycerophosphocholine (GPC), phosphocholine (PCh), glutathione (GSH), myo-inositol (Ins), lactate (Lac), N-Acetylaspartate (NAA), N-Acetylaspartateglutamate (NAAG) and taurine (Tau). It is often quite difficult to resolve Glu from Gln, NAA from NAAG and PCh from GPC, particularly if the spectral quality is poor due to an inadequate shim. So, in addition to the principal metabolites the total of Glu and Gln, NAA and NAAG and PCh and GPC are also reported. These total concentrations are thought to be a more reliable metric. An unsuppressed water signal was also used for absolute concentration calculation and eddy current correction. This internal reference method assumes known values of water concentrations of gray and white matter^5-7^.

There are several limitations to ^1^H NMR spectroscopy which can make reliable measures particularly challenging. One such limitation is that is that ^1^H NMR metabolite measures are inherently low in signal to noise. The metabolite signals measured are approximately 10,000-fold less than the proton signal used in imaging. The EC is also a small structure and thus requires a small VOI which reduces the measured signal strength. However, at 7T with shim values in the range 8 – 15 Hz and a short echo time of 8 ms, the spectral peaks were reasonably separated, allowing for direct and reliable measures of metabolite concentrations. For metabolites which have significant spectral overlap, total values (e.g. Glu and Gln) are also reported. Another error source is chemical shift displacement. The bandwidth of the 180° RF pulse is 5 kHz, which would give a displacement error of 0.3 mm in the vertical and horizontal directions over the range of 0.2 to 4.2 ppm. However, this displacement was relatively small and still allowed for sufficient specificity. Despite the challenges of in vivo mouse brain ^1^H NMR, we were able to achieve high quality spectra in this difficult region with high quantification accuracy.

### Data Availability

The transcriptomics datasets are available in the NCBI Gene Expression Omnibus (GEO) repository: https://www.ncbi.nlm.nih.gov/geo/query/acc.cgi?acc=GSE102334. The metabolomics datasets are available in the MetaboLights repository: http://www.ebi.ac.uk/metabolights/MTBLS530

## Supporting information

## ACKNOWLEDGEMENTS

We acknowledge Drs. Wai-Haung Yu and Natura Myeku for advice on this manuscript, as well as Laidon Piroli for assistance with genotyping. We also acknowledge Dr. Paul Mathews for providing the APOE mice that were used in the ^1^H NMR experiment. This work was supported by grants from NIA and NINDS to K.E.D. (AG048408 and NS071836), grants from NIA to E.A.G. (AG045335 and AG056387), grants from NICHD and NHLBI to S.S.G. (HD67244 and HL87062), grants from NIA and NINDS to E.F. (AG057331 and NS089076), a grant from NCCR (1S10RR023534-01) for the 7T rodent scanner at Nathan Kline Institute, grants from the U.S. Dept. of Defense to D.L., (W911NF-12-1-9159 and W911F-15-1-0169), funding from the Cure Alzheimer’s Fund to KD and TN, and fellowships from NIA (AG047797) and the Brightfocus Foundation to T.N. In addition, this work was supported in part by the Intramural Research Program of the NIH, National Institute on Aging.

## AUTHOR CONTRIBUTIONS

This study was designed and managed by T.N., E.A.G., and K.E.D. Animal care and breeding was performed by H.F. Transcriptomic analysis was performed by A.A.D., in the laboratory of M.R.C. Metabolomic analysis was performed by Q.C. and T.N., in the laboratory of S.S.G. Seahorse analysis was performed by D.L. and M.P., in the laboratory of E.A.G. PIUMet analysis was performed by L.P., in the laboratory of E.F. ^1^H NMR spectroscopy was performed by K.S. and D.G. Additional molecular biology experiments and bioinformatics were performed by T.N., A.A. and H.A. Manuscript preparation was performed by T.N. and E.A.G., with input from K.E.D.

## Supplementary Material

**Fig. 6. Targeted lipidomics reveals differential expression of diacyglycerols and bis(monacylglycero)phosphates in the EC of aged *APOE4* mice.**

A targeted lipidomic analysis was performed in order to analyze the effects of differential *APOE* isoform expression on lipid levels in the EC and PVC of aged *APOE* mice (8 *APOE3/3*, 9 *APOE3/4* and 7 *APOE4/4* males). (A-C) Shown here are graphs of the three primary groups of lipids that were shown to be differentially expressed in the EC of the *APOE4/4* vs. *APOE3/3* mice: (A) Diacylglycerols, (B) Bis(monoacylglycero)phosphates, and (C) Hexosylceramides, which include both glucosylceramides and galactosylceramides. (* denotes p < 0.05; ** denotes p < 0.01; *** denotes *p* < 0.001; **** denotes p < 0.0001)

**Fig. S1.**
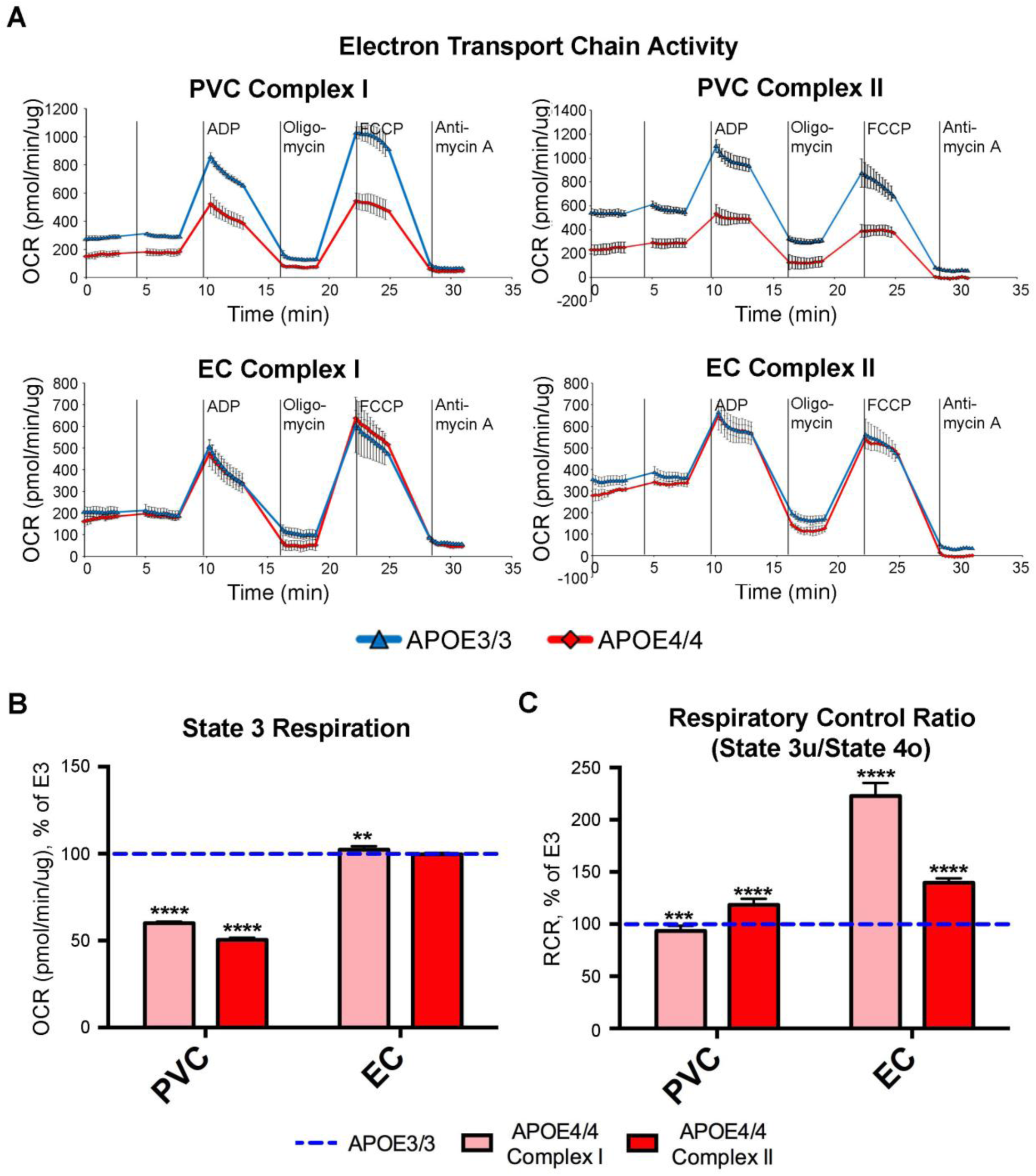
Seahorse analysis reveals decreased mitochondrial respiration in the PVC, but not in the EC, of aged *APOE4* mice. Seahorse analysis was performed in order to analyze the effects of differential *APOE* isoform expression on mitochondrial respiration in mitochondria that were purified from the PVC and EC of aged *APOE* mice (4 *APOE4/4* males, tissues pooled vs. 4 *APOE3/3* males, tissues pooled). (A) The complex I- and complex II-driven ETC activity from each region shows decreased mitochondrial respiration in the mitochondria purified from the PVC, but not the EC of the aged *APOE4/4* mice. (B-C) Bar graphs showing the average oxygen consumption rate (OCR) from (B) State 3 and for (C) the Respiration Control Ratio (RCR; state 3u/state 4o) in mitochondria purified from each region of the *APOE4/4* mice, as a percentage of the *APOE3/3* OCR from the equivalent tissues. The dotted blue line represents the normalized levels in the *APOE3/3* tissues. (* denotes p < 0.05; ** denotes p < 0.01; *** denotes *p* < 0.001; **** denotes p < 0.0001)

**Fig. S2.**
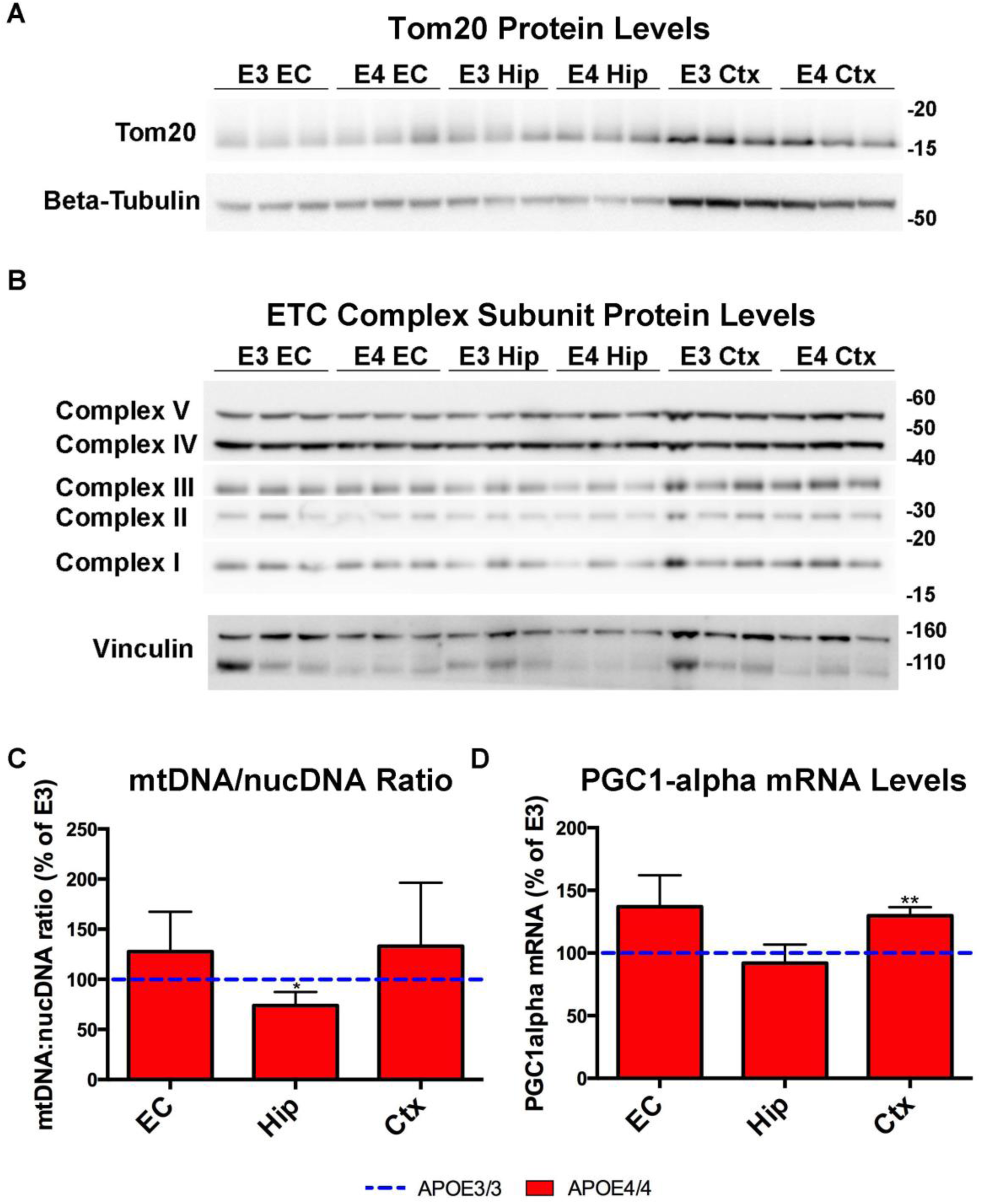
Western blot and qPCR analysis shows no significant effects of APOE4 on mitochondrial mass or ETC protein expression. Western blotting and qPCR analysis was performed in order to investigate the effects of differential APOE isoform expression on various markers of mitochondrial mass and ETC proteins in the EC, Hip and Ctx of 21-month-old *APOE4/4* vs. *APOE3/3* mice. (A-B) Western blot analysis did not reveal any unique differences in the levels of mitochondrial outer membrane protein Tom20 or in ETC complex subunits between genotypes in the EC. Likewise, (C) the ratio of Mitochondrial:Nuclear DNA, as well as (D) the levels of PGC1α RNA were unchanged between *APOE* genotypes in the EC of 21-month-old *APOE* mice. (* denotes p < 0.05; ** denotes p < 0.01)

**Fig. S3.**
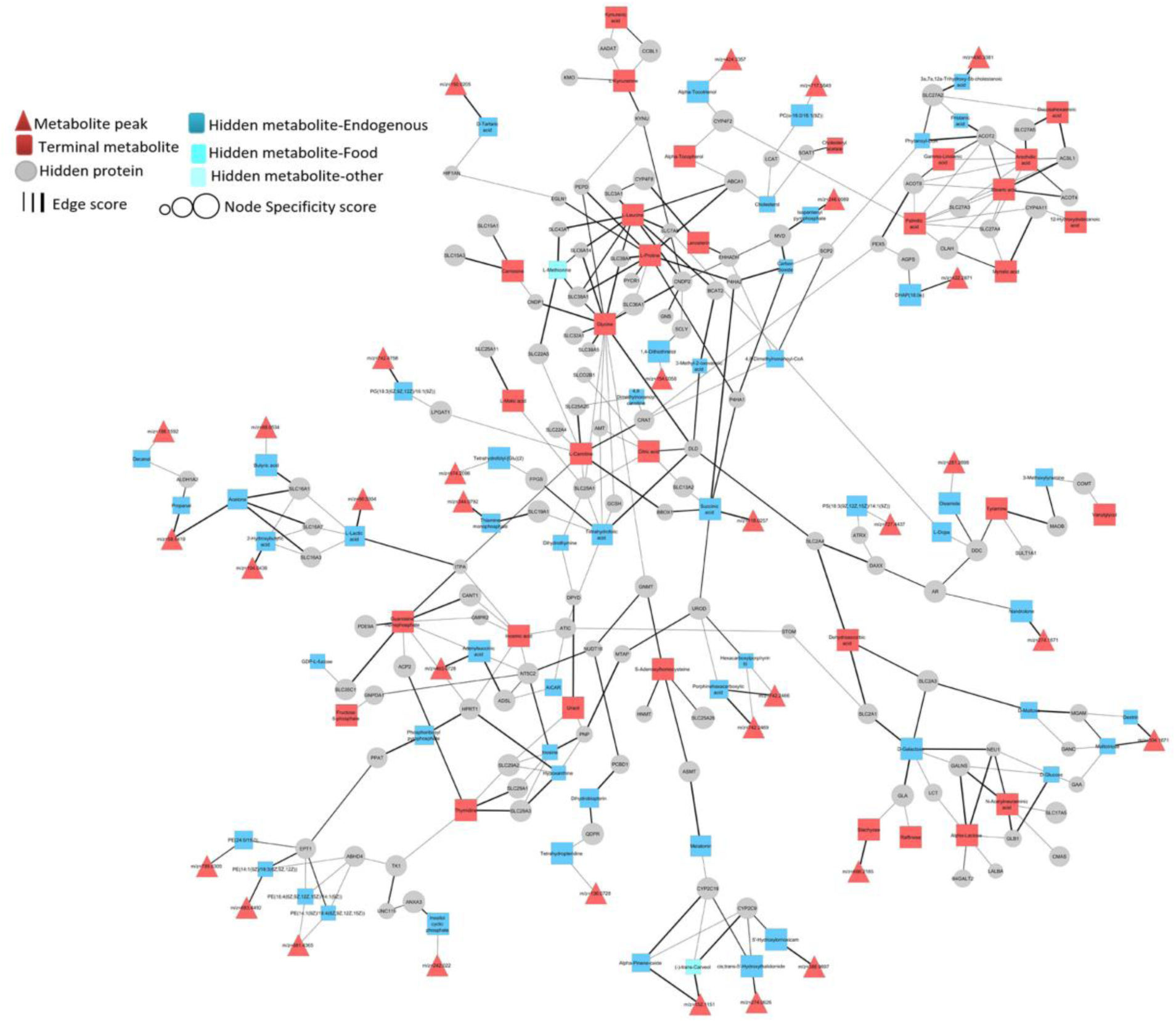
Results network from the PIUMet analysis of differentially expressed untargeted metabolites from the EC of *APOE4/4* vs. *APOE3/3* mice. Shown here is the results-network that was generated from the PIUMet (Prize-collecting Steiner forest algorithm for Integrative Analysis of Untargeted Metabolomics) analysis. PIUMet was able to predict identities for the differential expressed unidentified metabolite features or peaks detected in the EC of aged *APOE4/4 vs. APOE3/3* mice (corrected p-value <0.05, 4 *APOE4/4* males, tissues pooled vs. 4 *APOE3/3* males, tissues pooled). The resulting network shows 242 nodes connected by 328 edges. Input metabolite peaks or features are shown as red triangles, while grey circles represent proteins and square nodes display the metabolites in the network. The metabolites directly connected to the differential metabolite features show their putative identities. The size of the nodes reflects the specificity score and the thickness of the edges the confidence of the interaction. In addition, the robustness score of each node is associated with the thickness of each node line width.

**Table S1.**
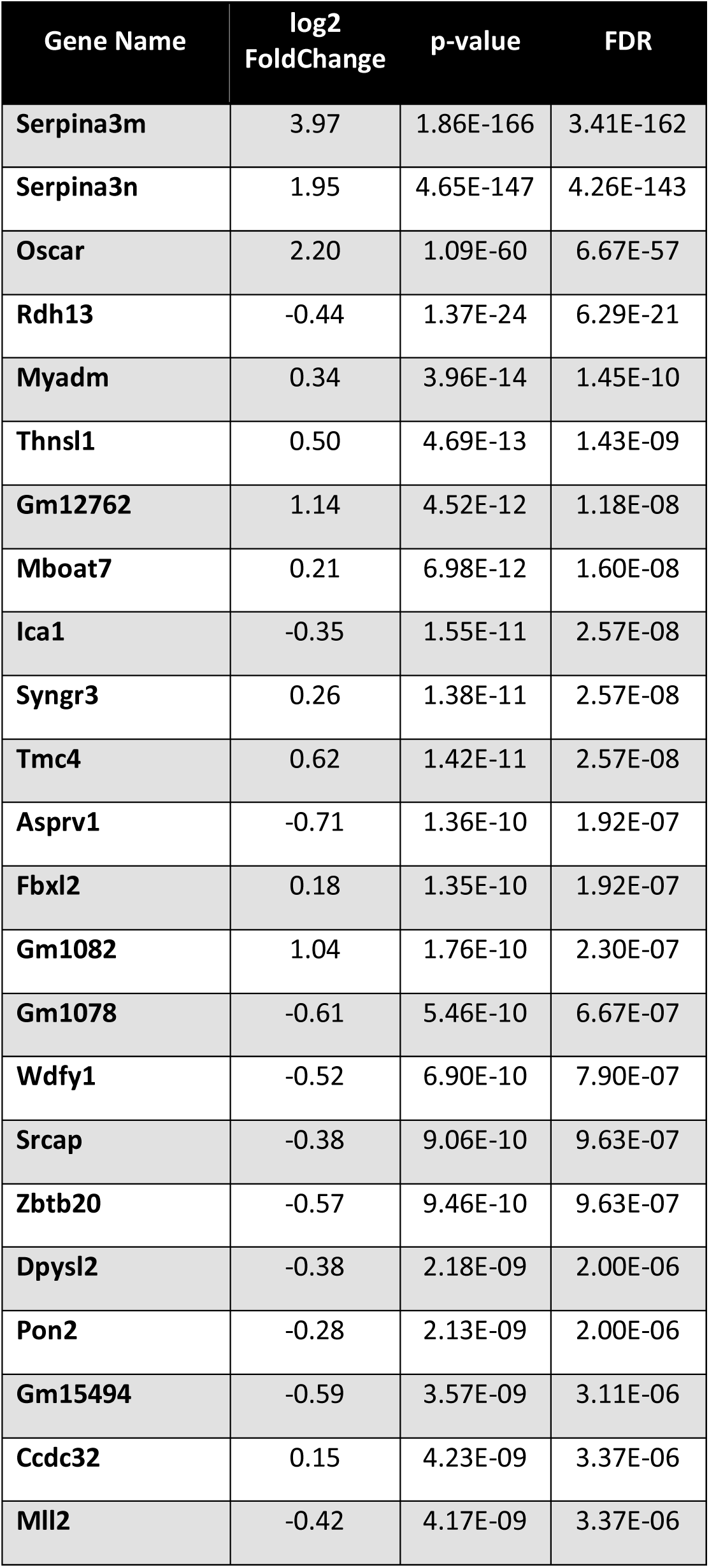
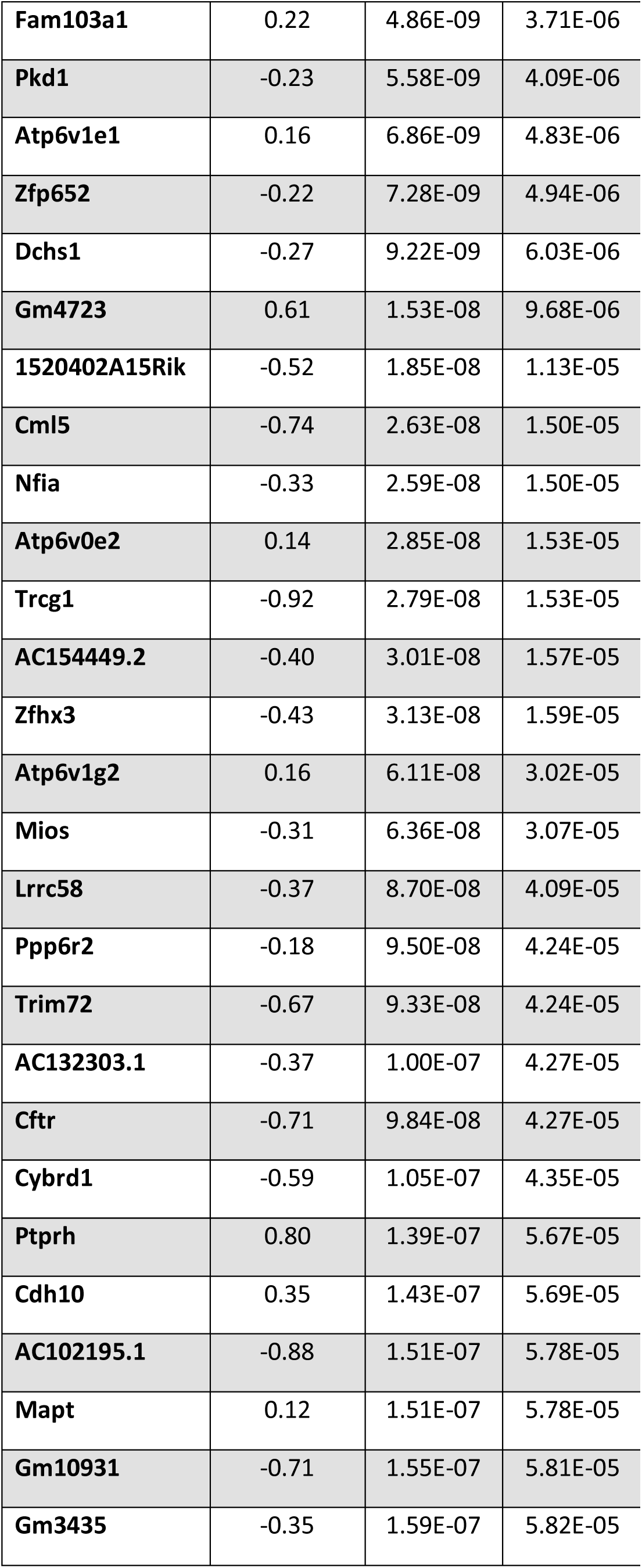
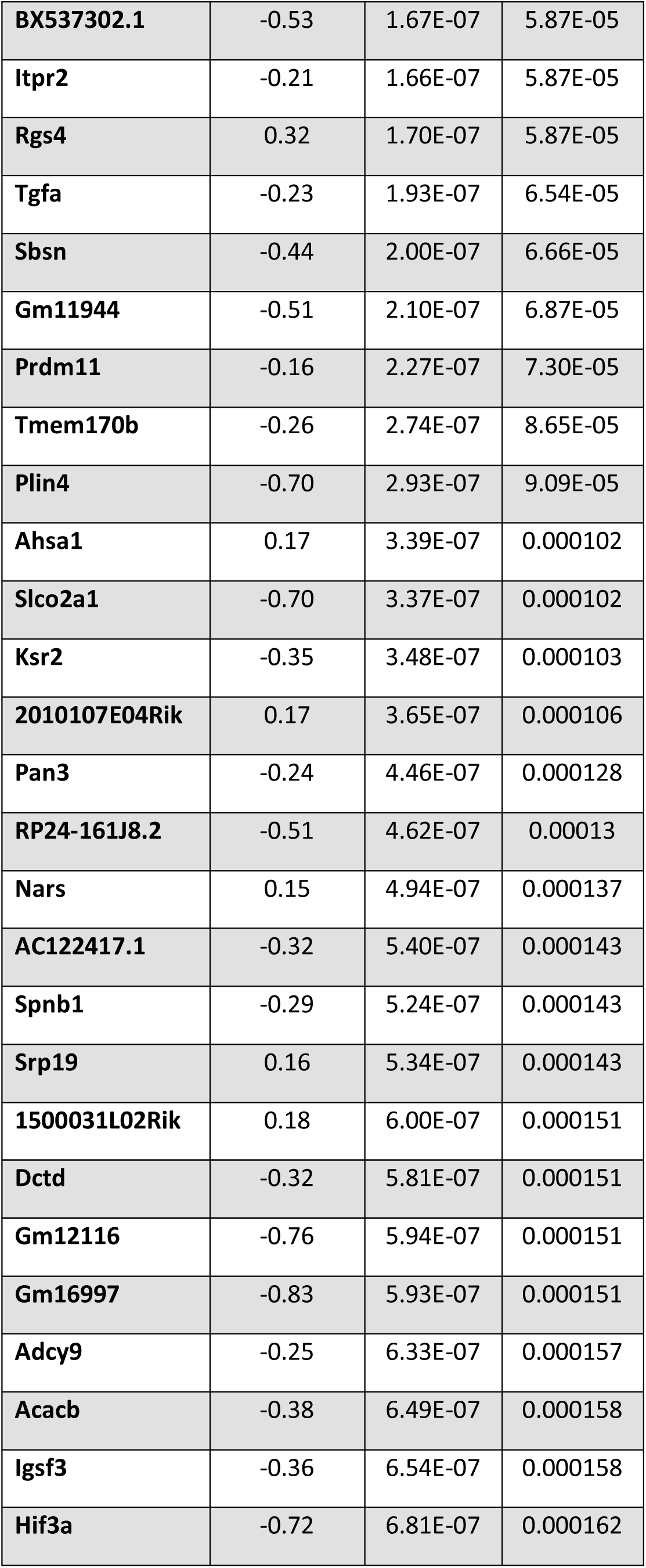
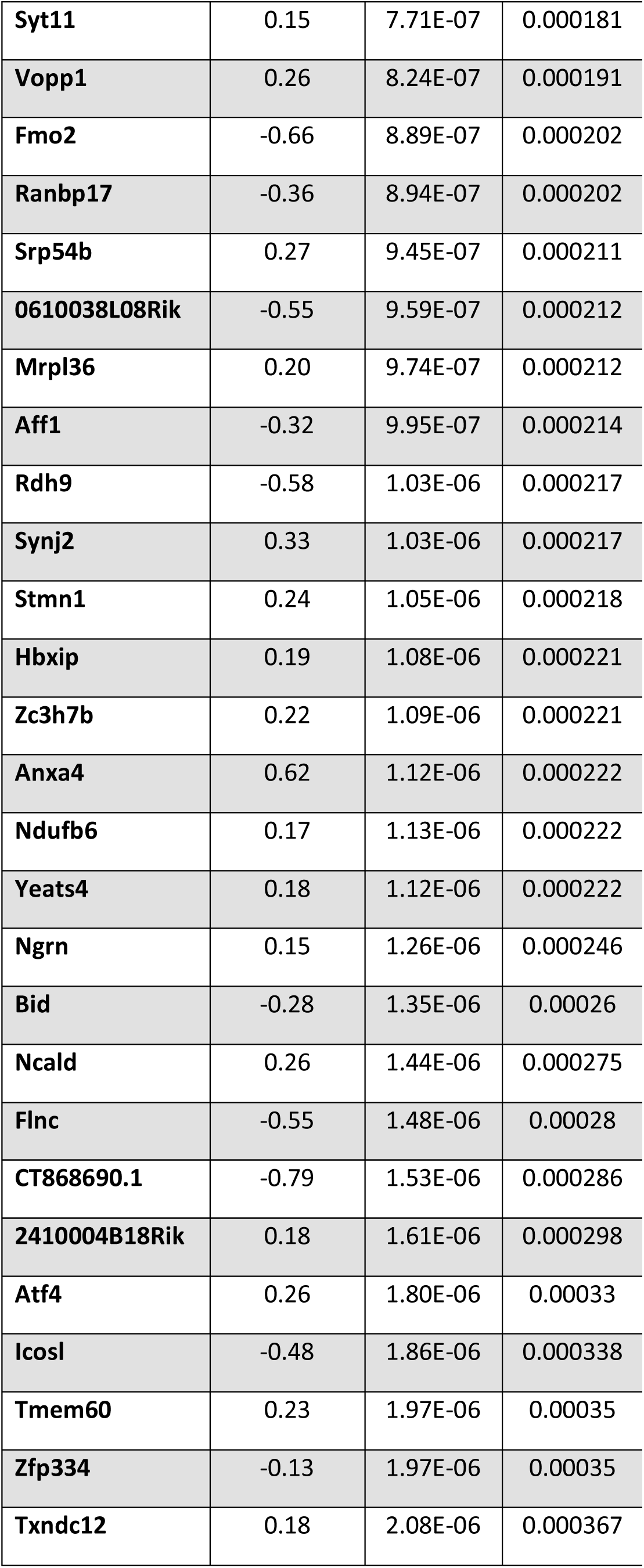
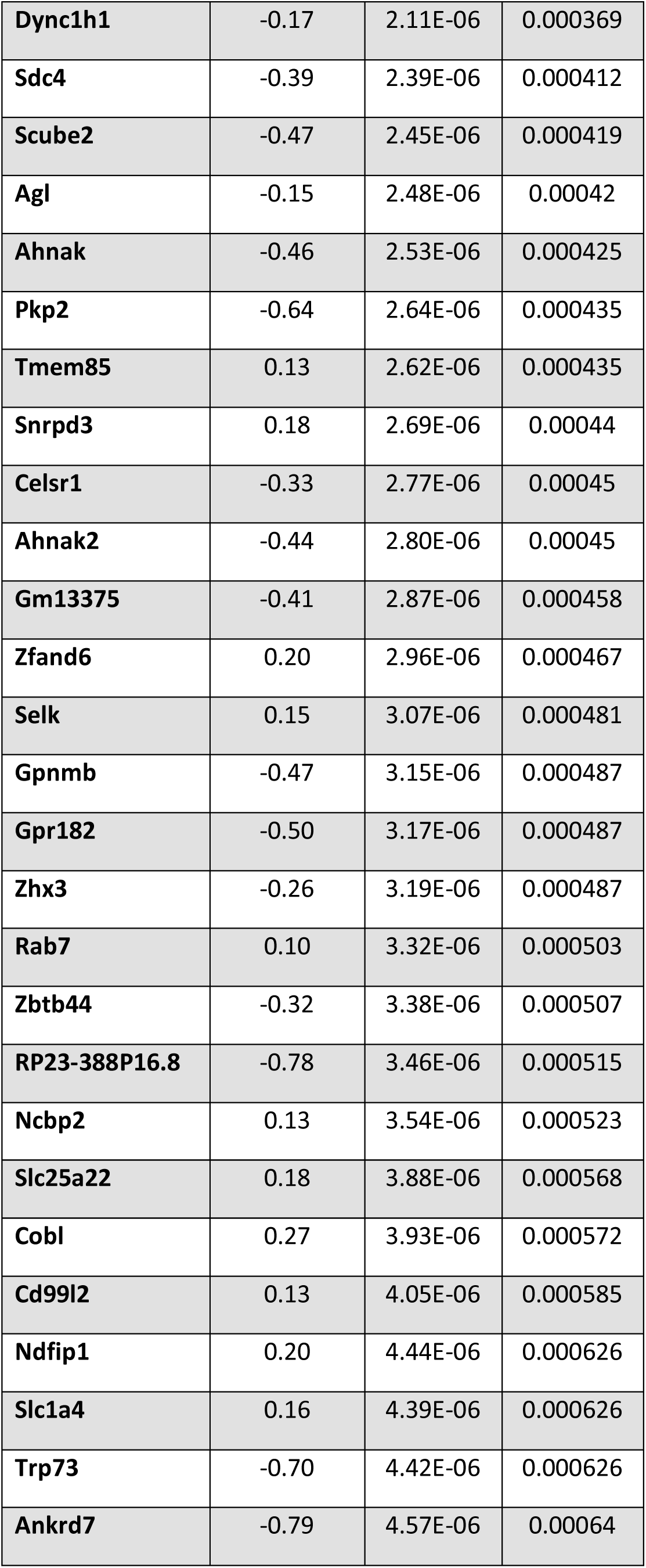
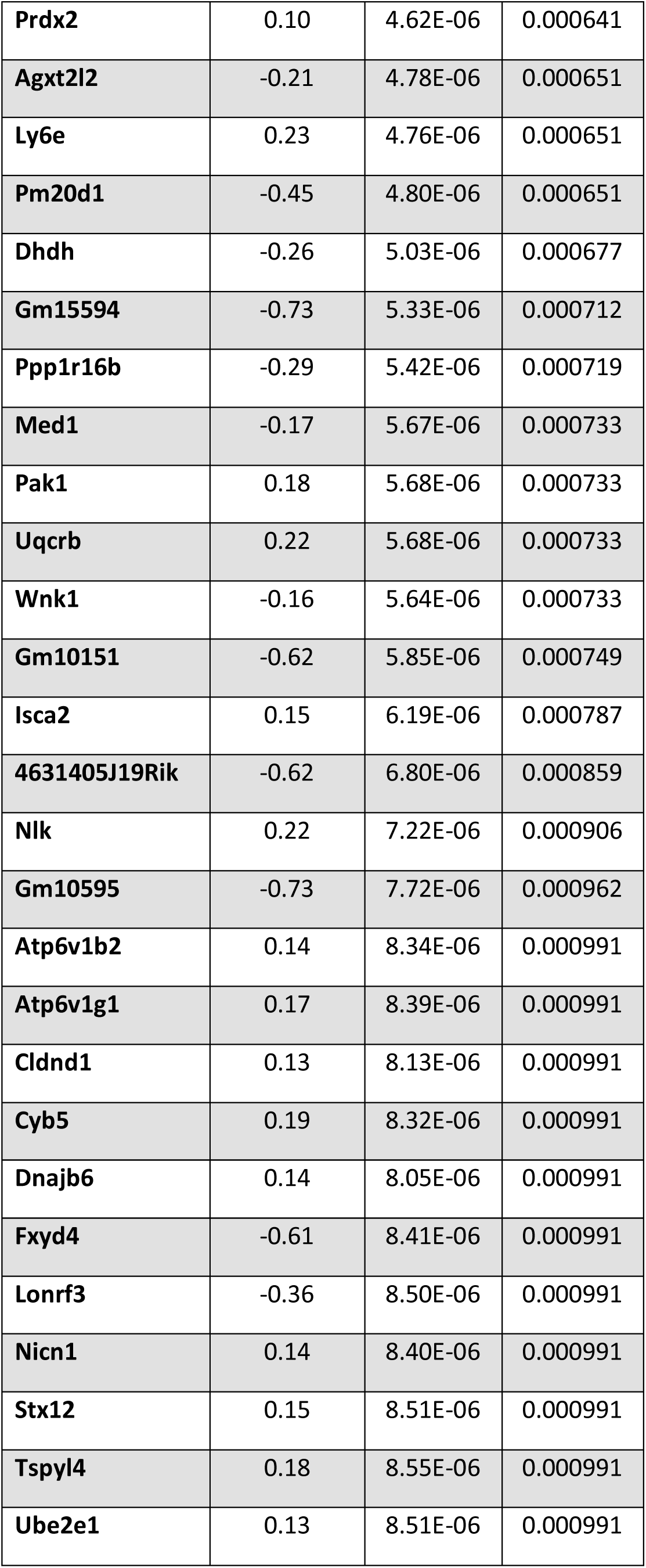
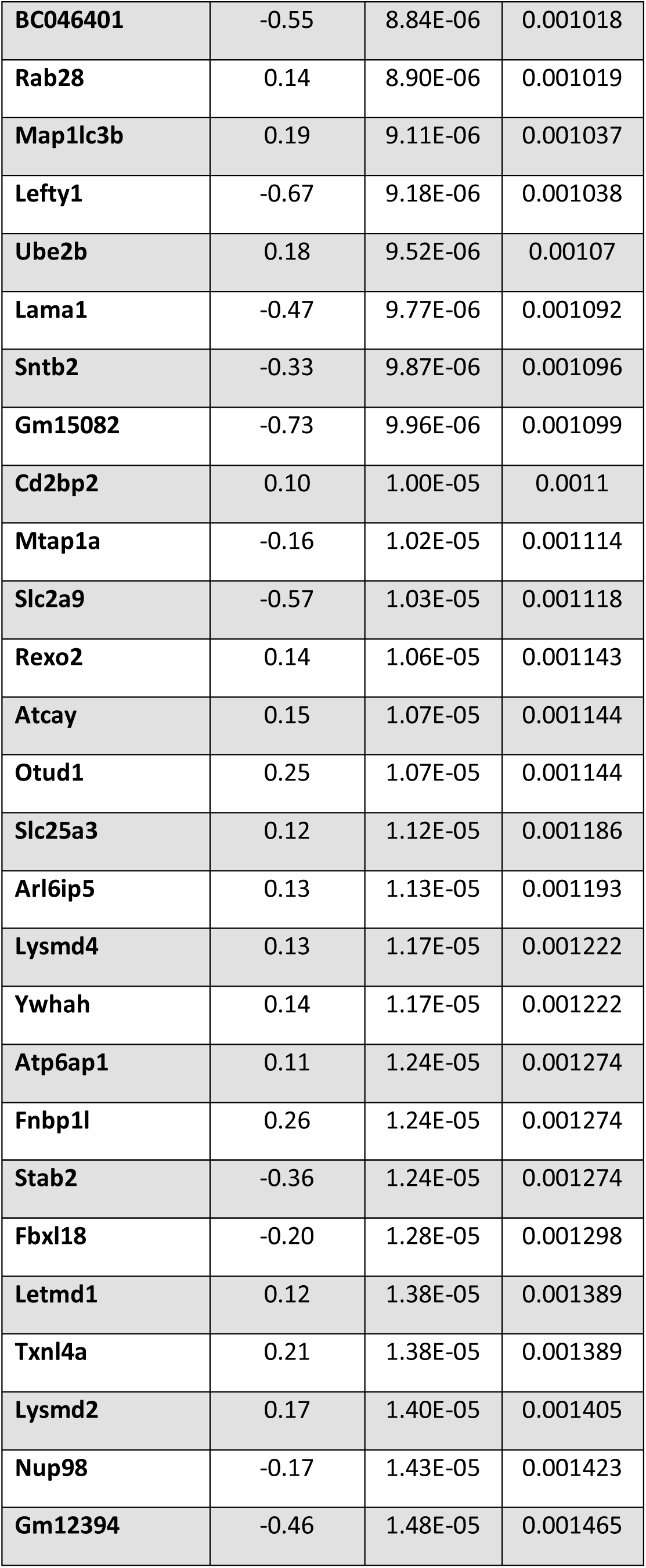
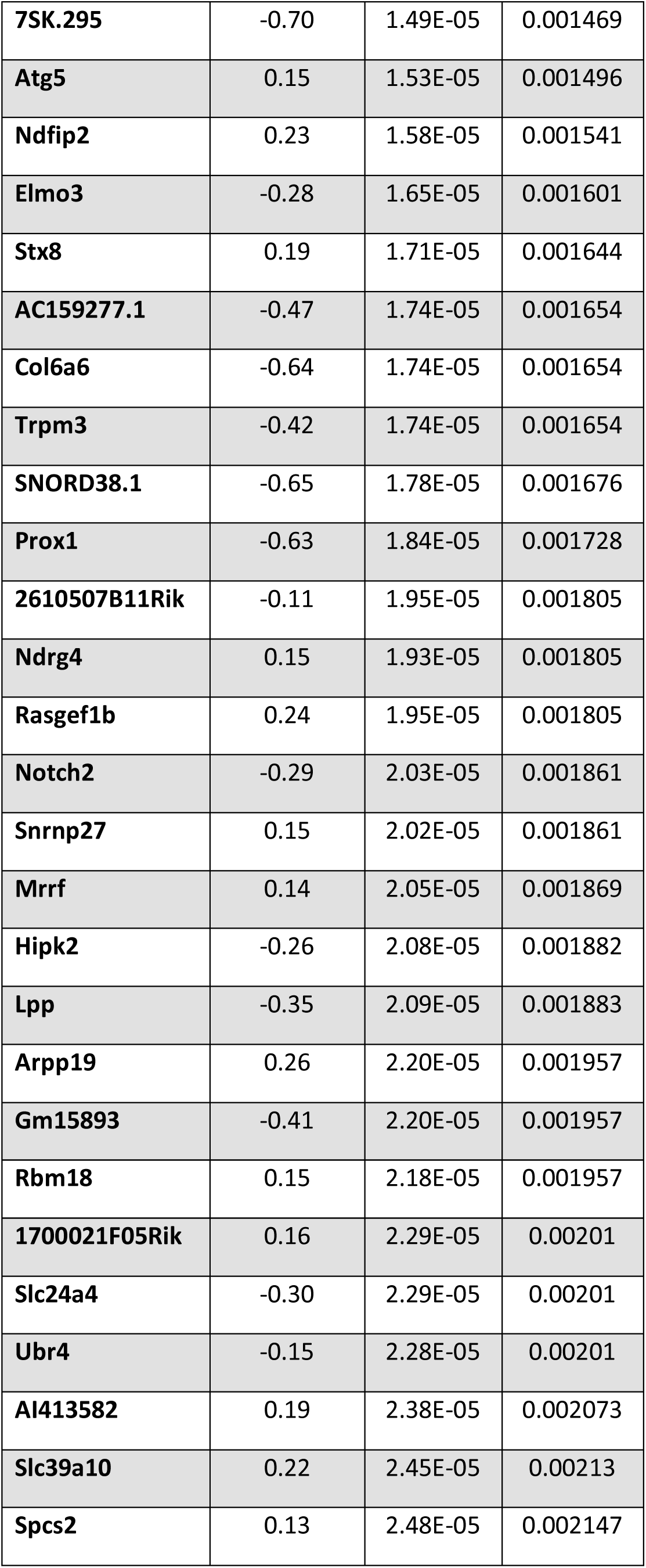
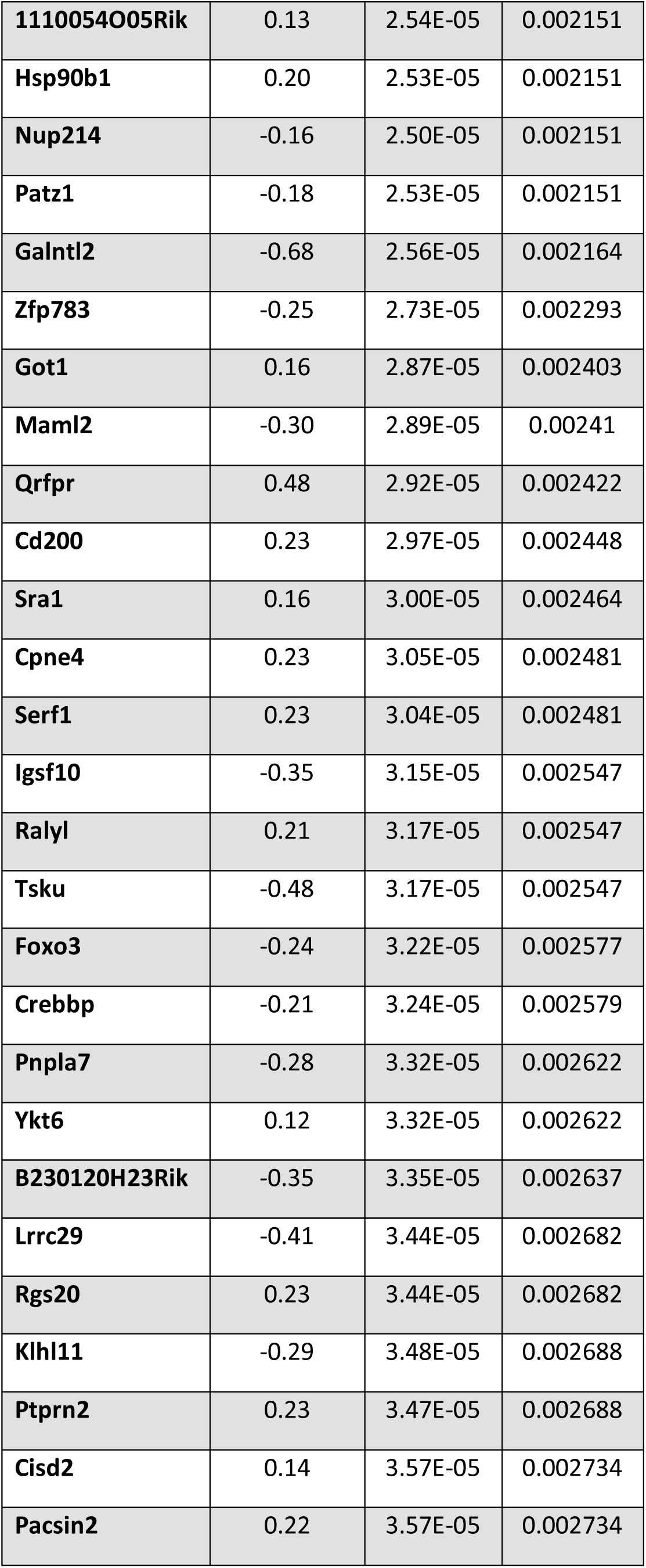
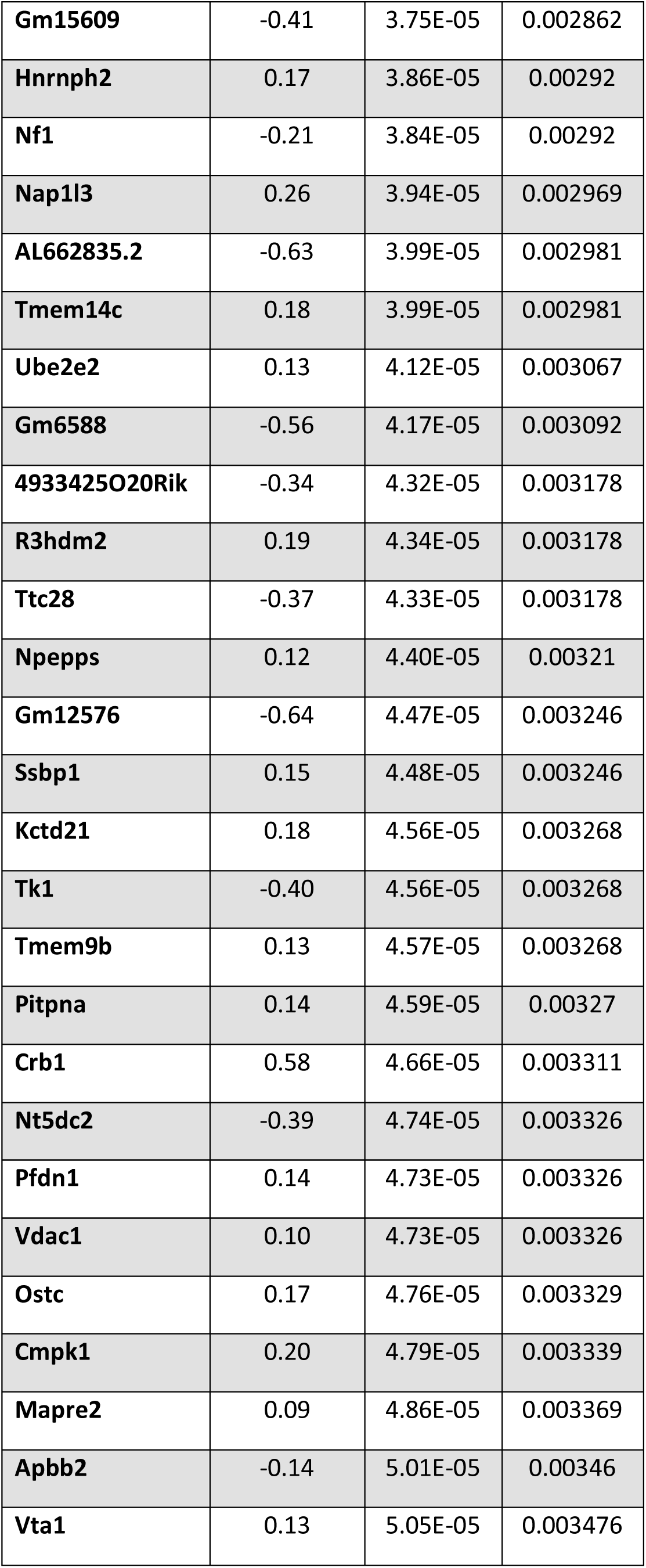
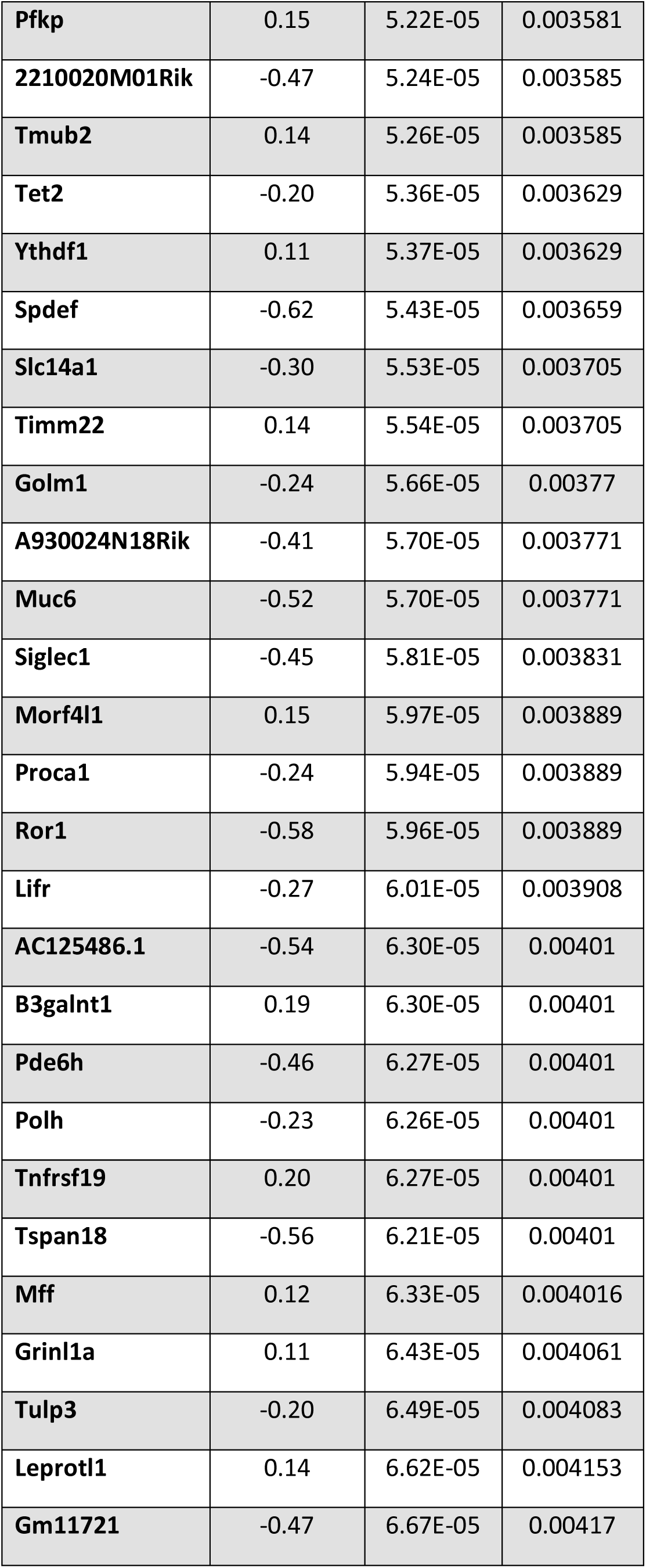
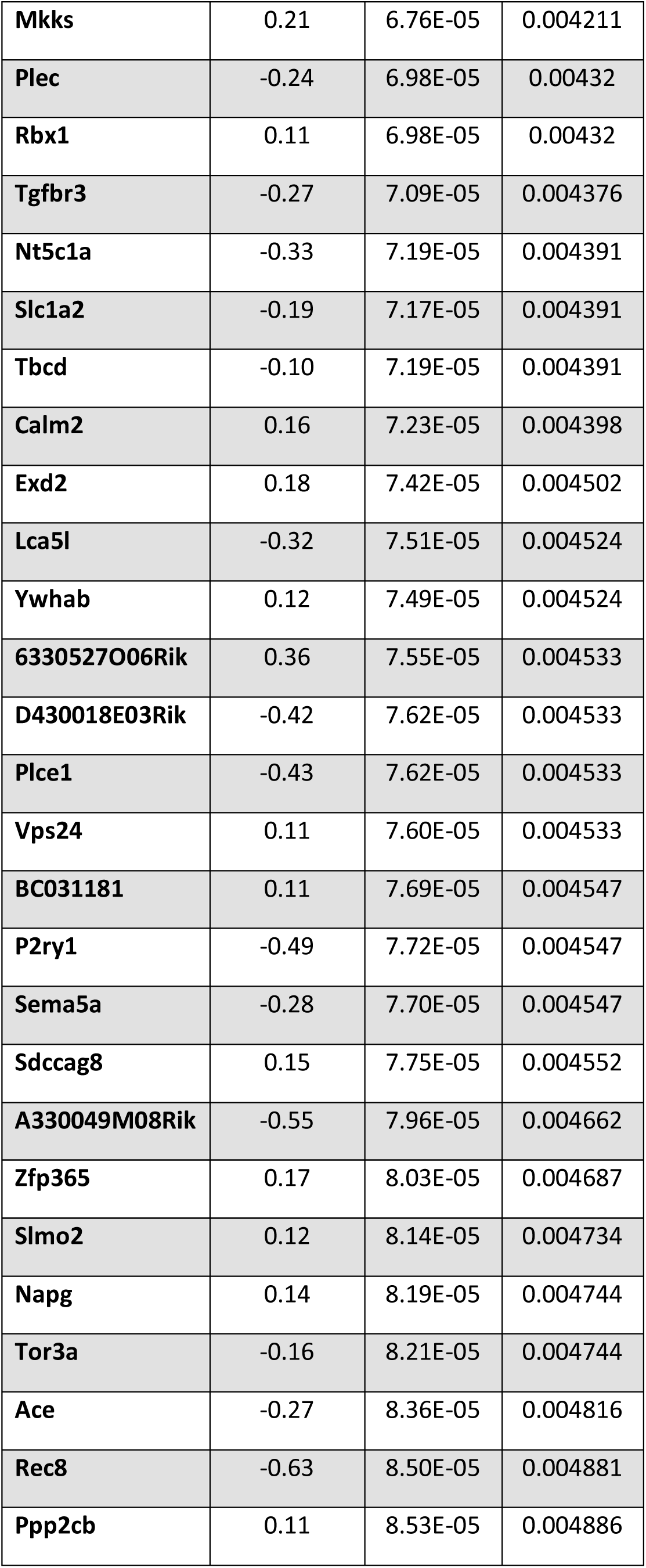
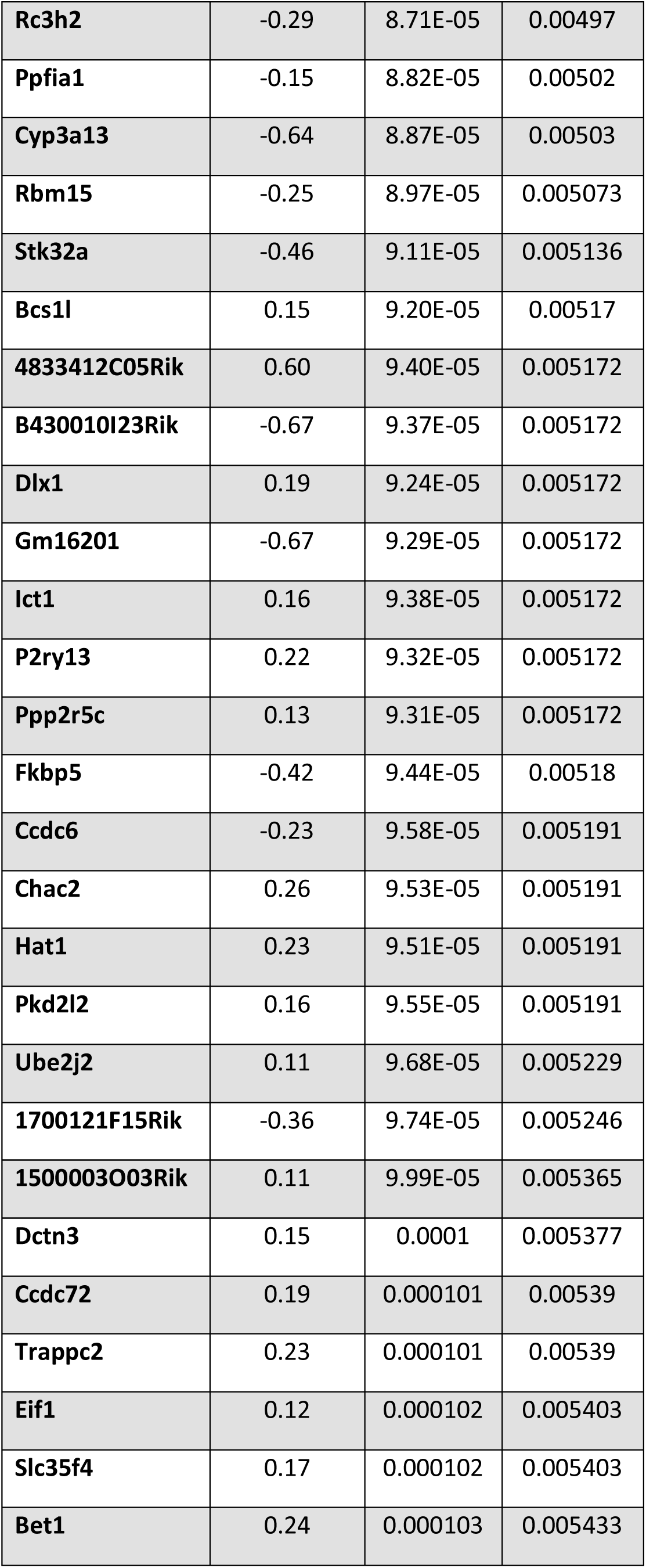
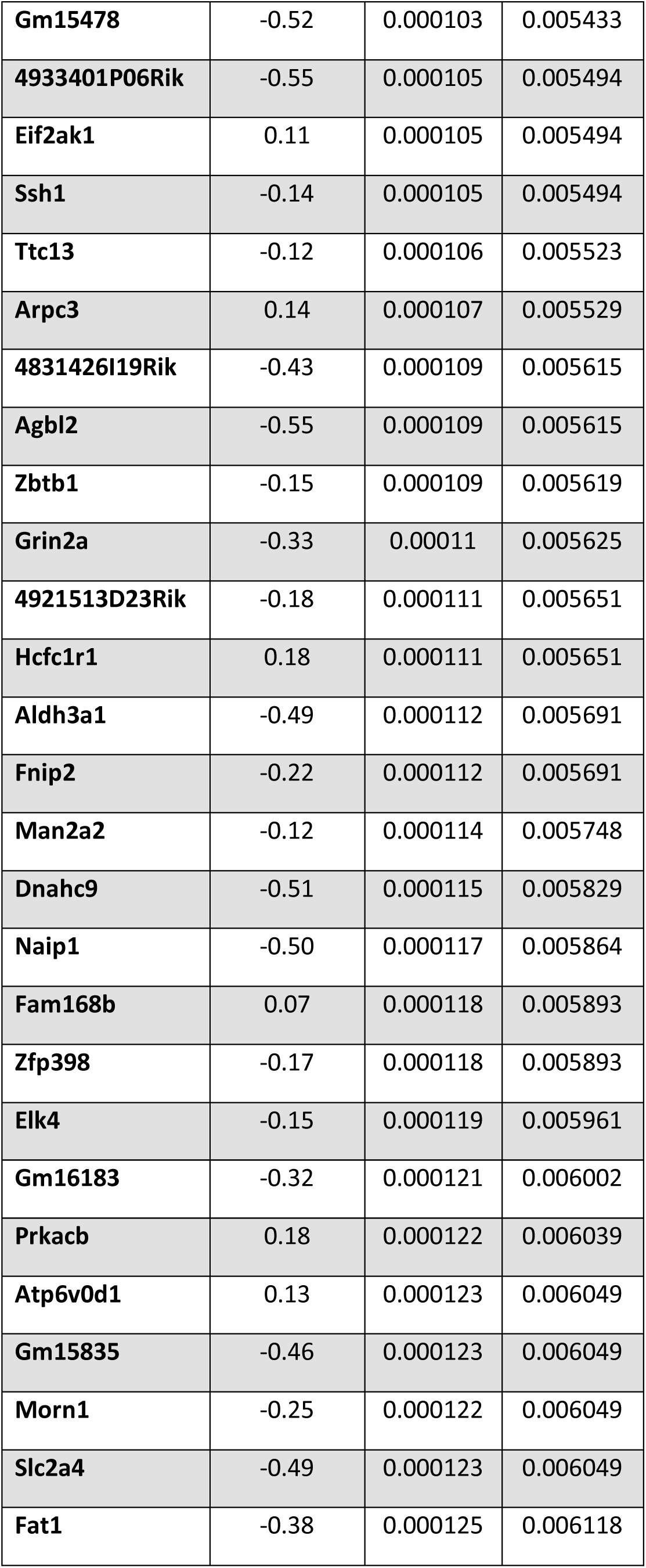
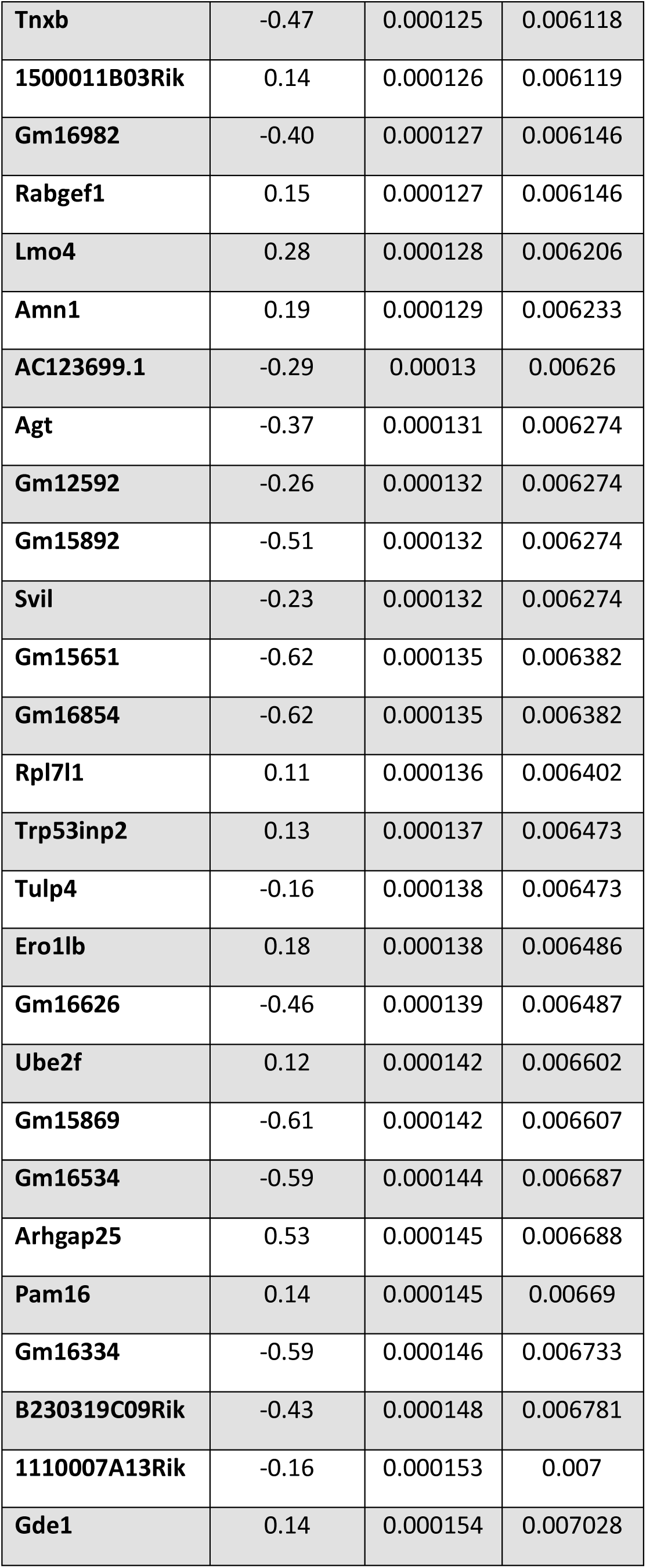
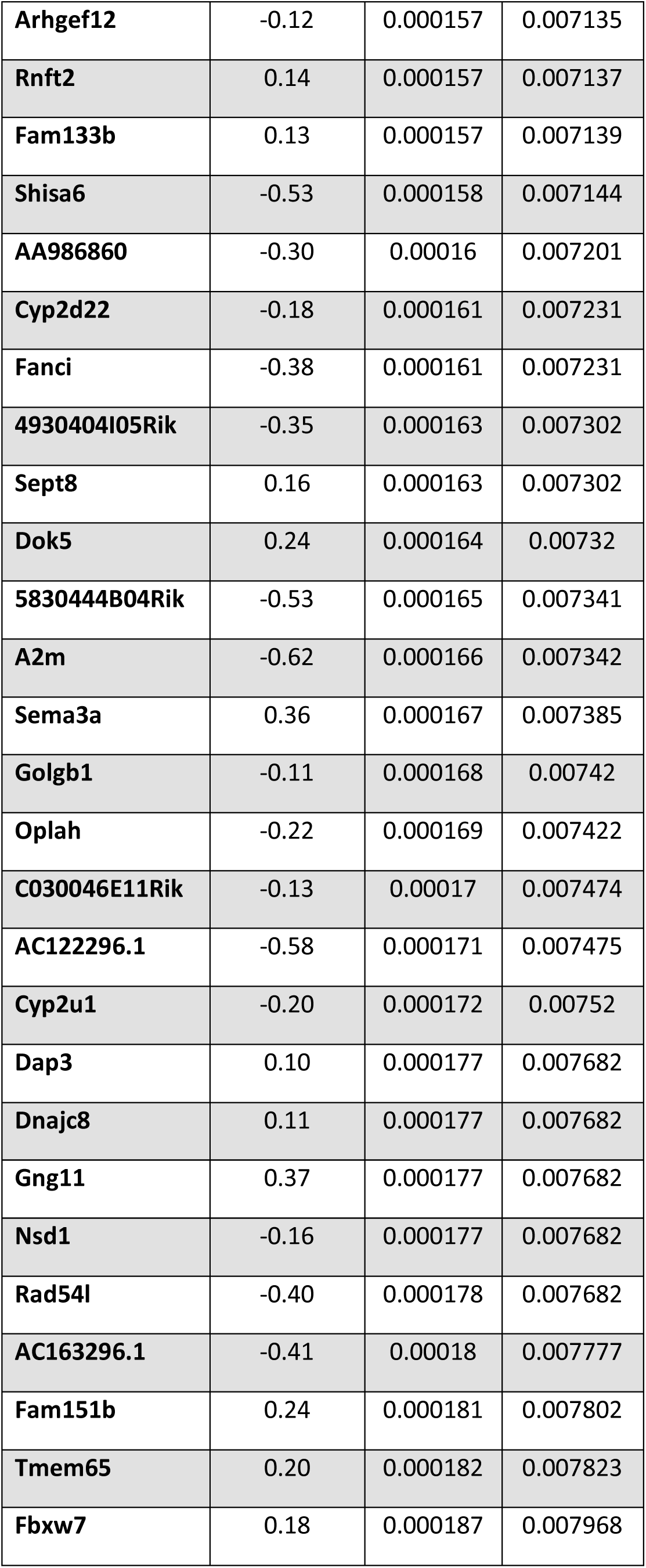
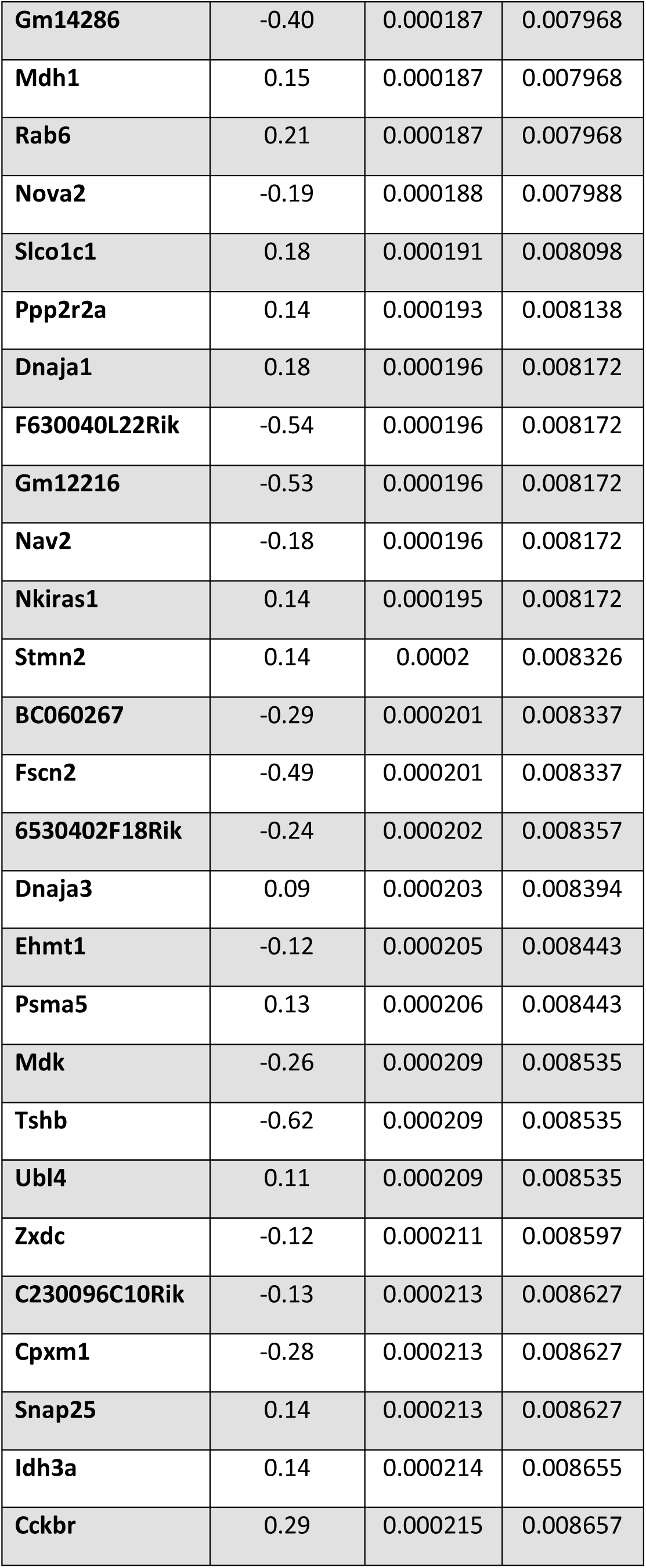
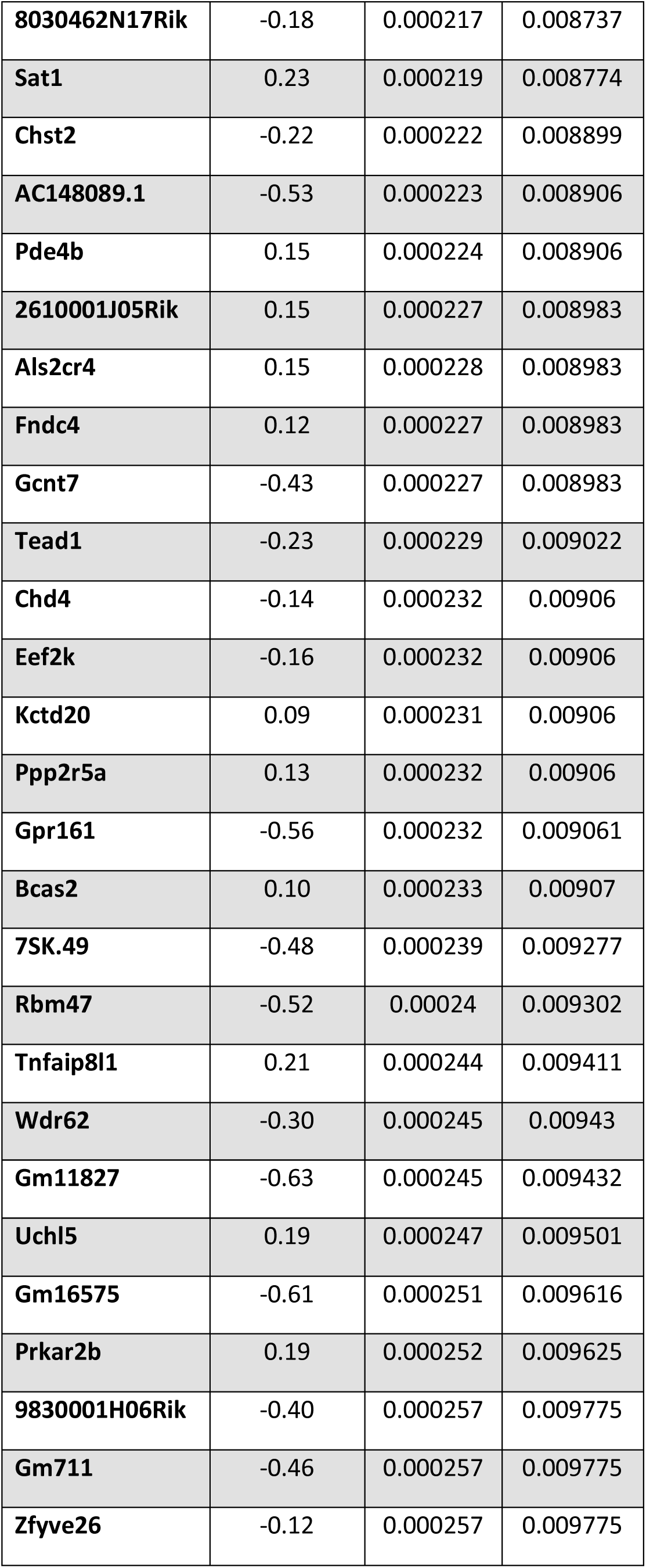
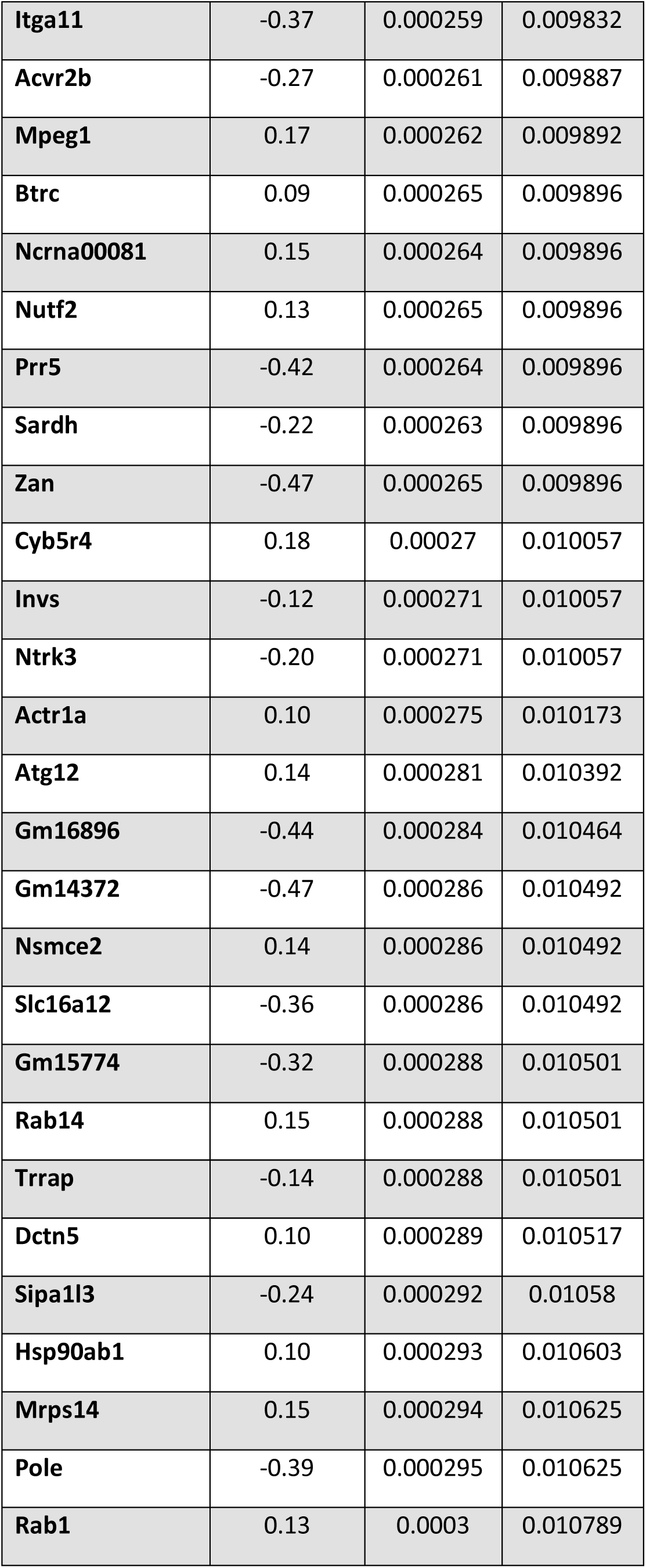
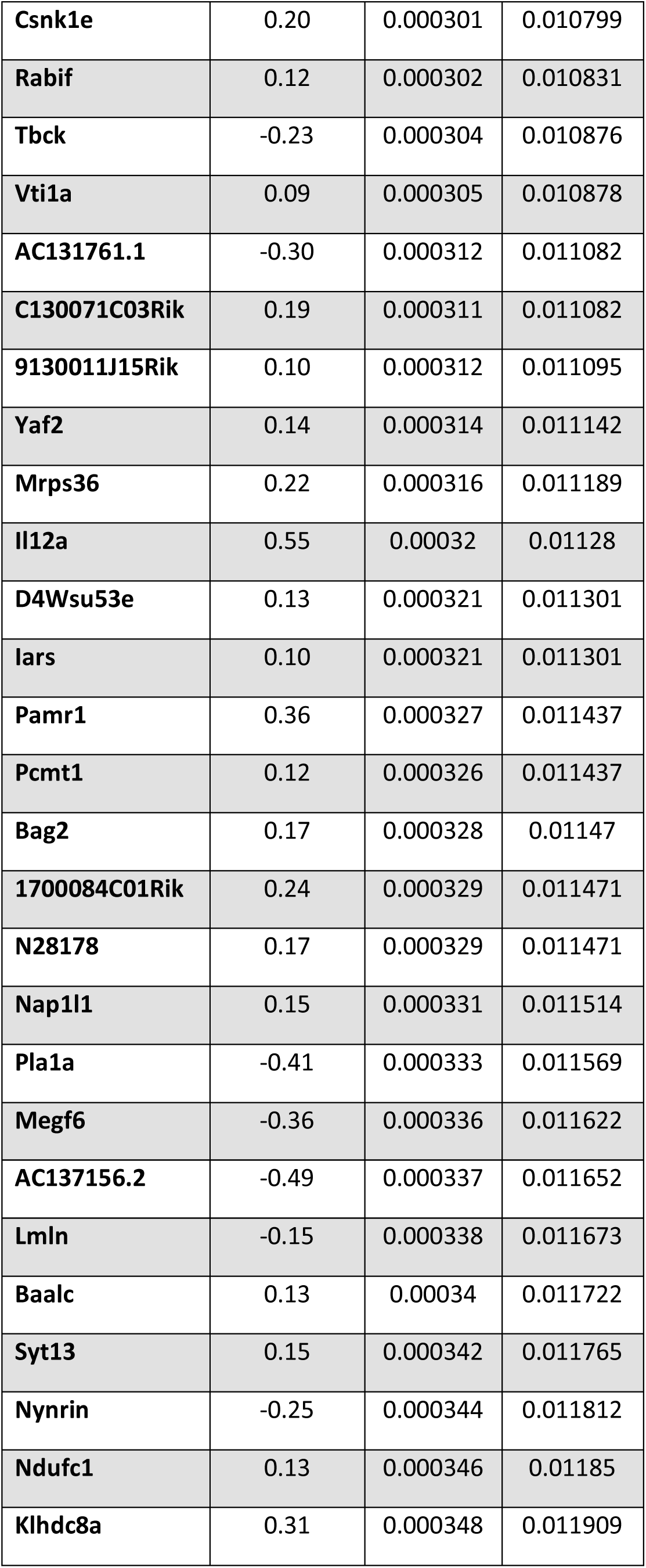
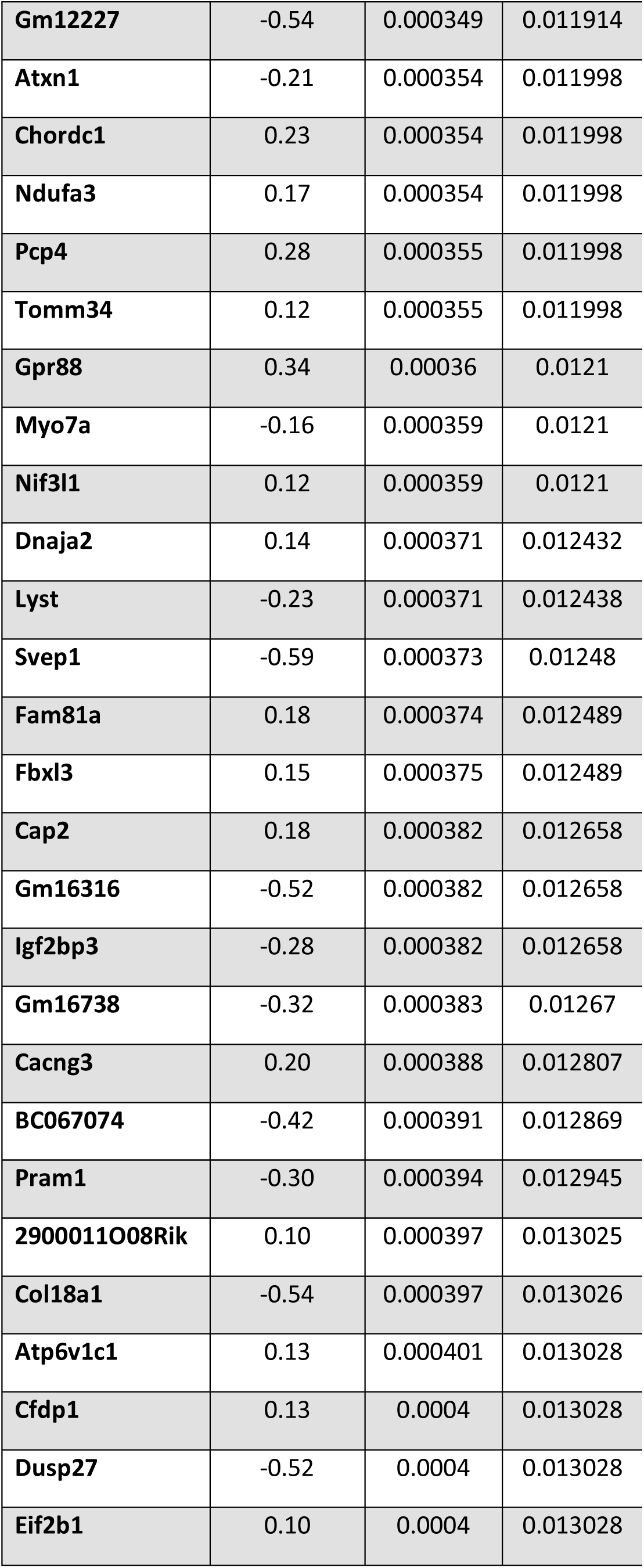
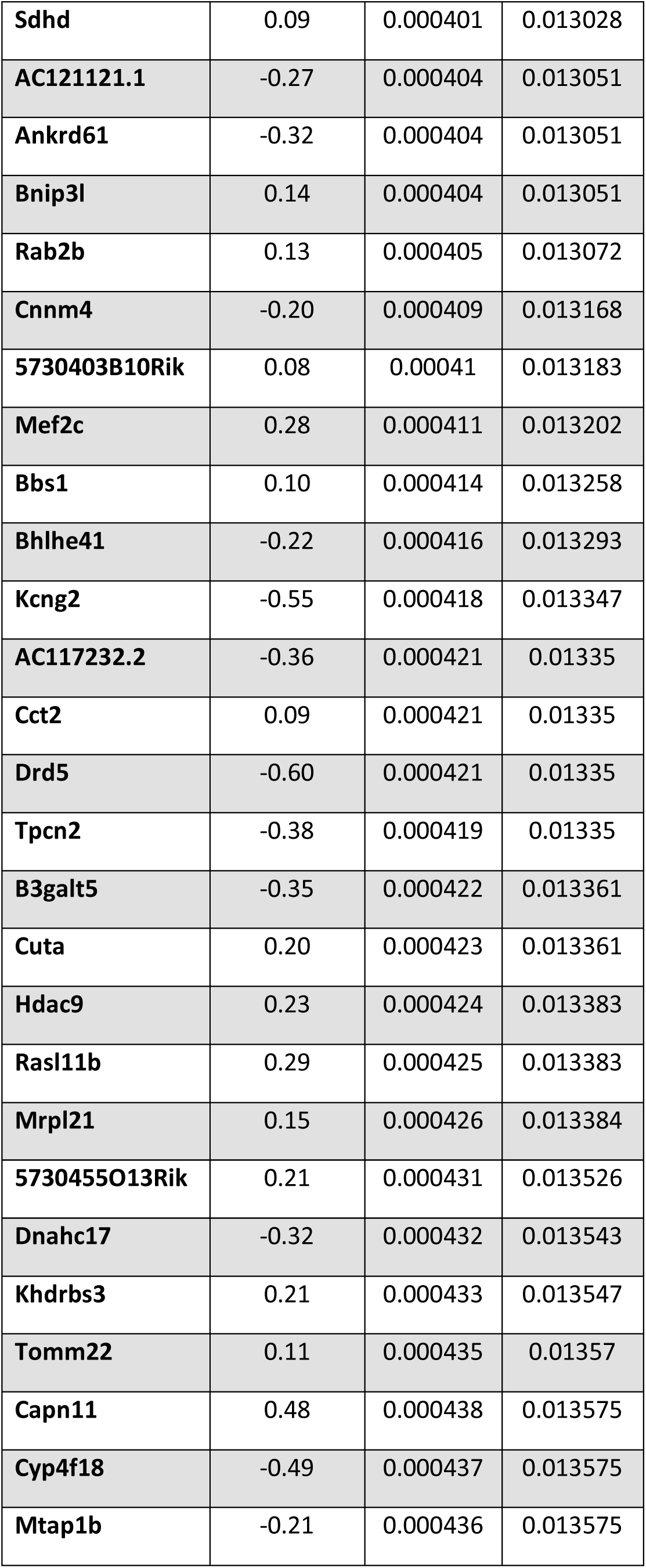
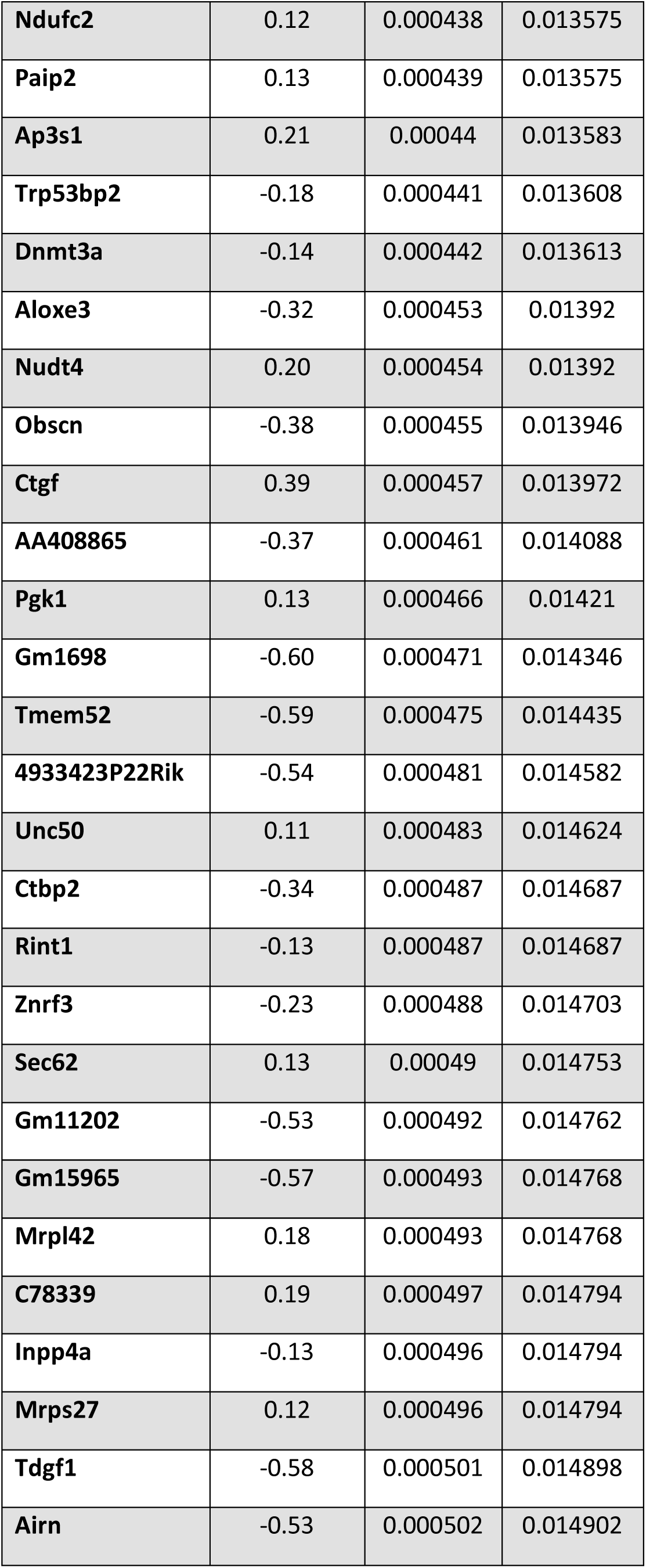
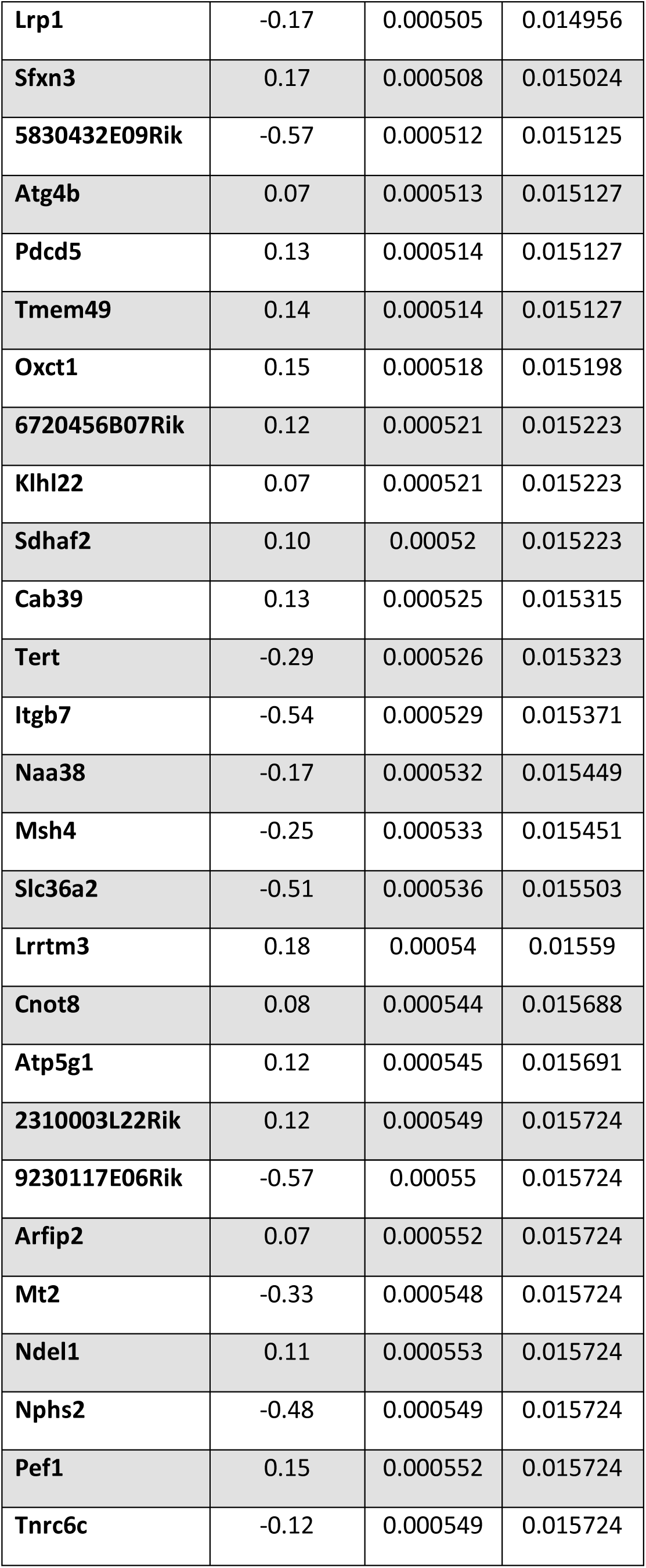
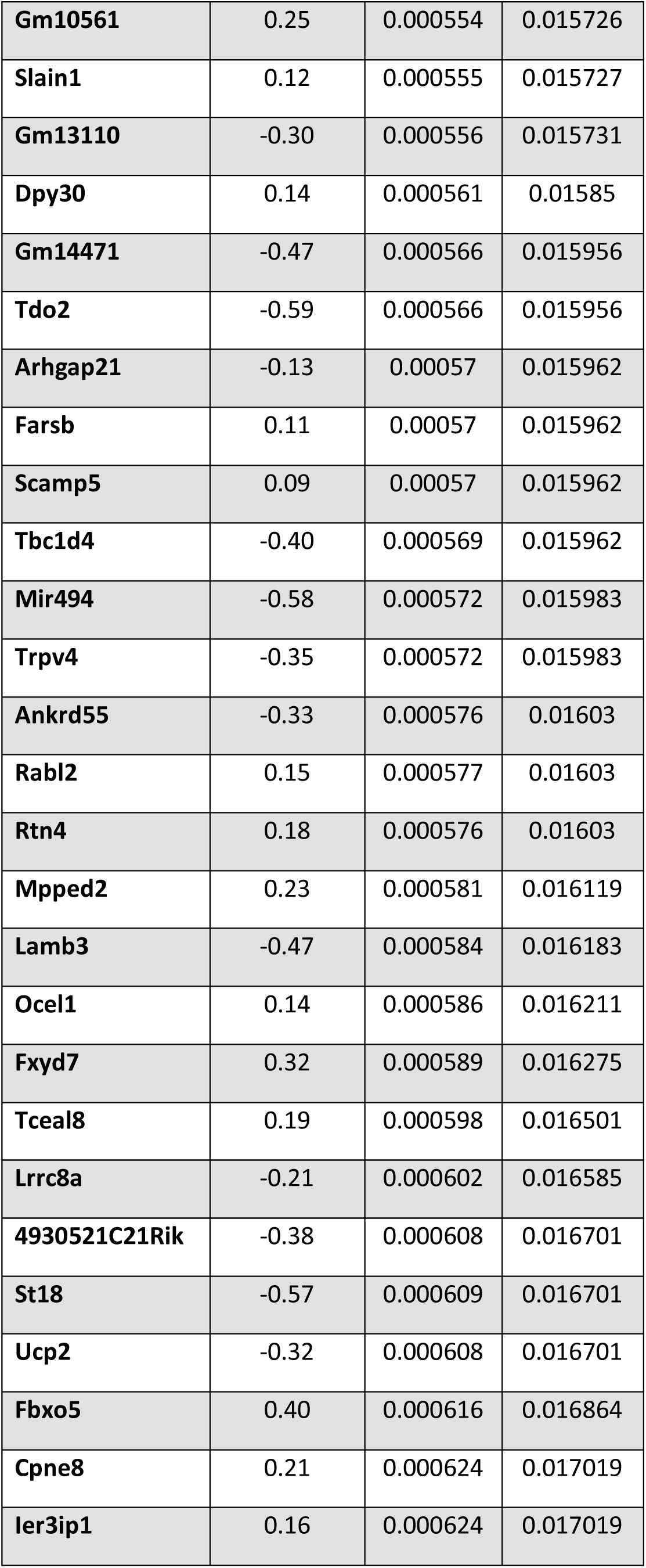
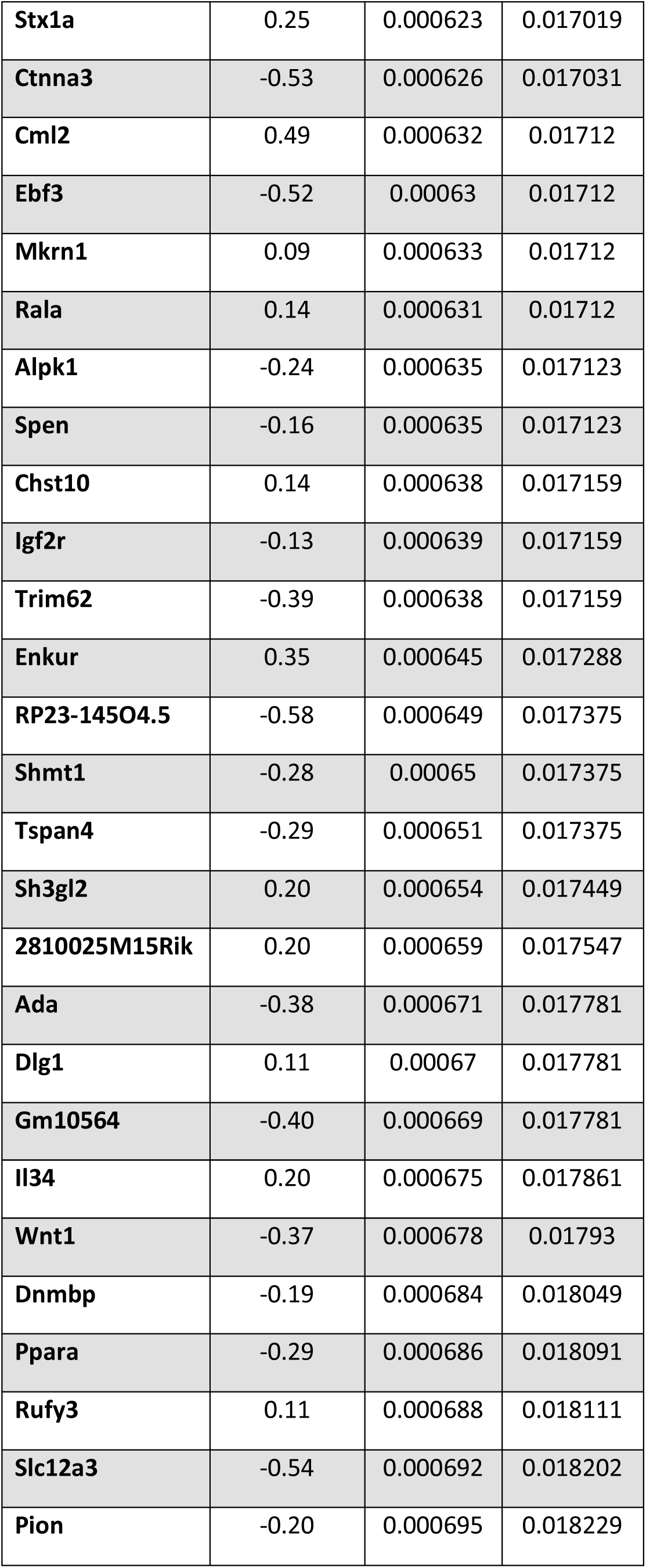
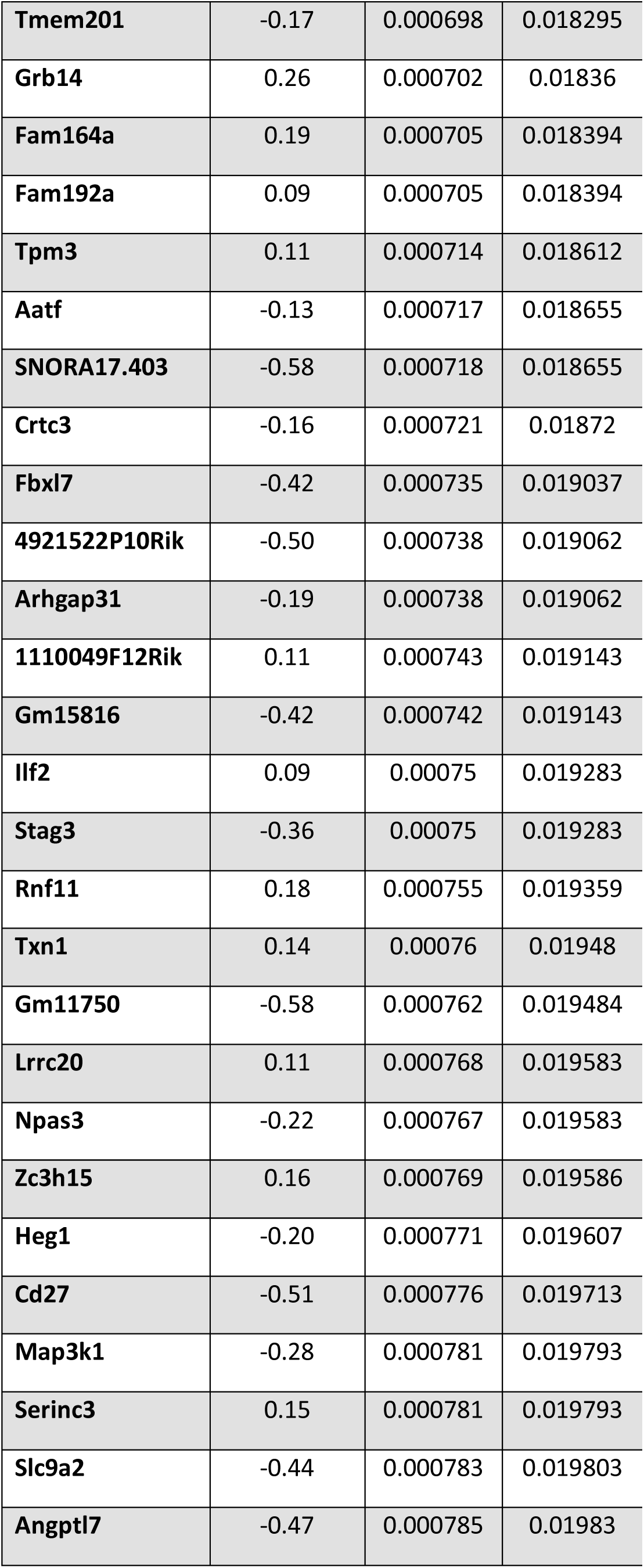
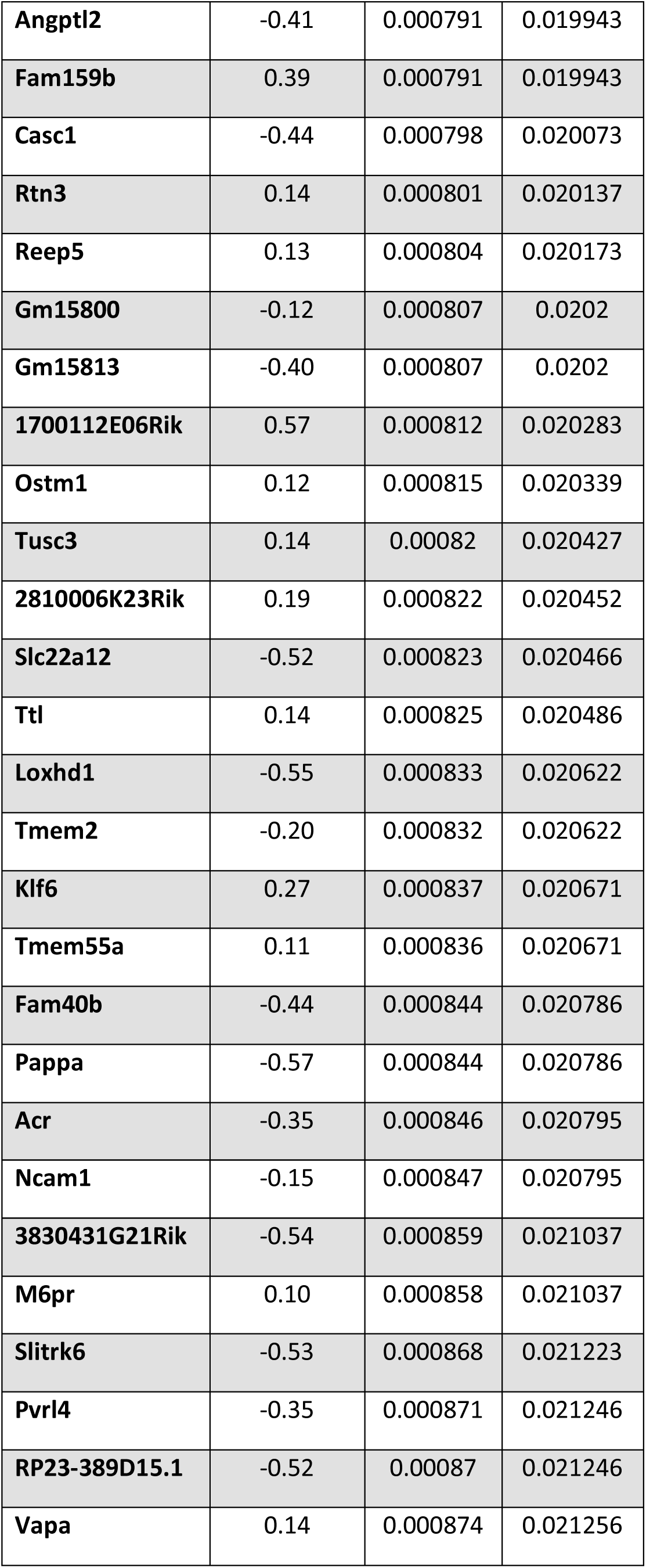
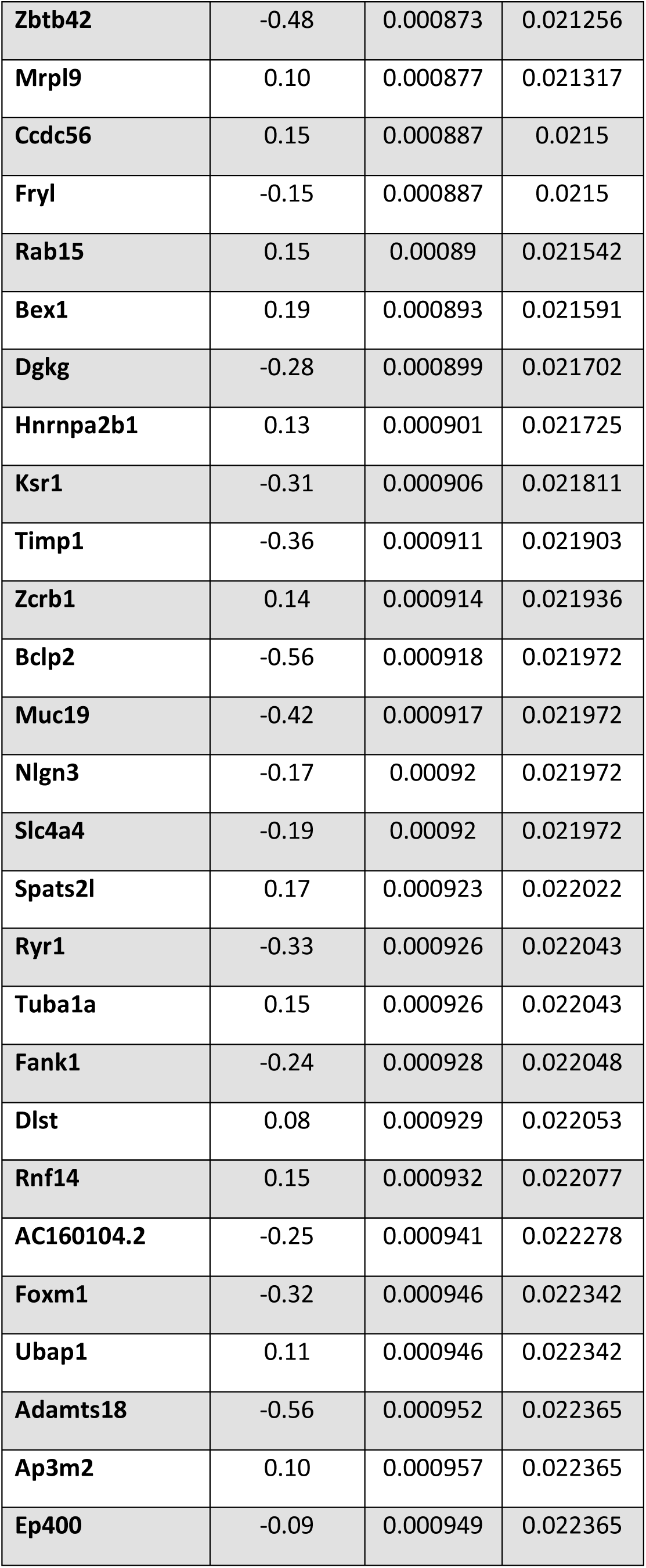
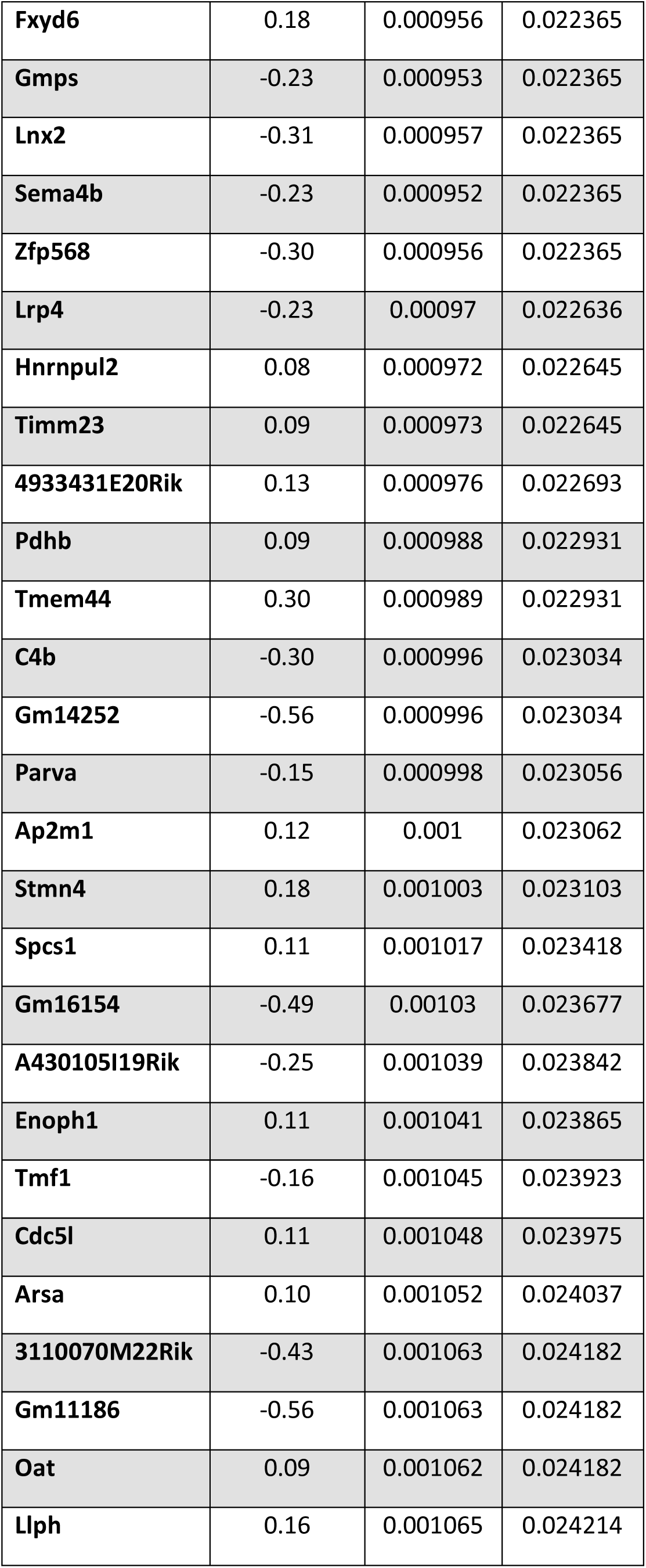
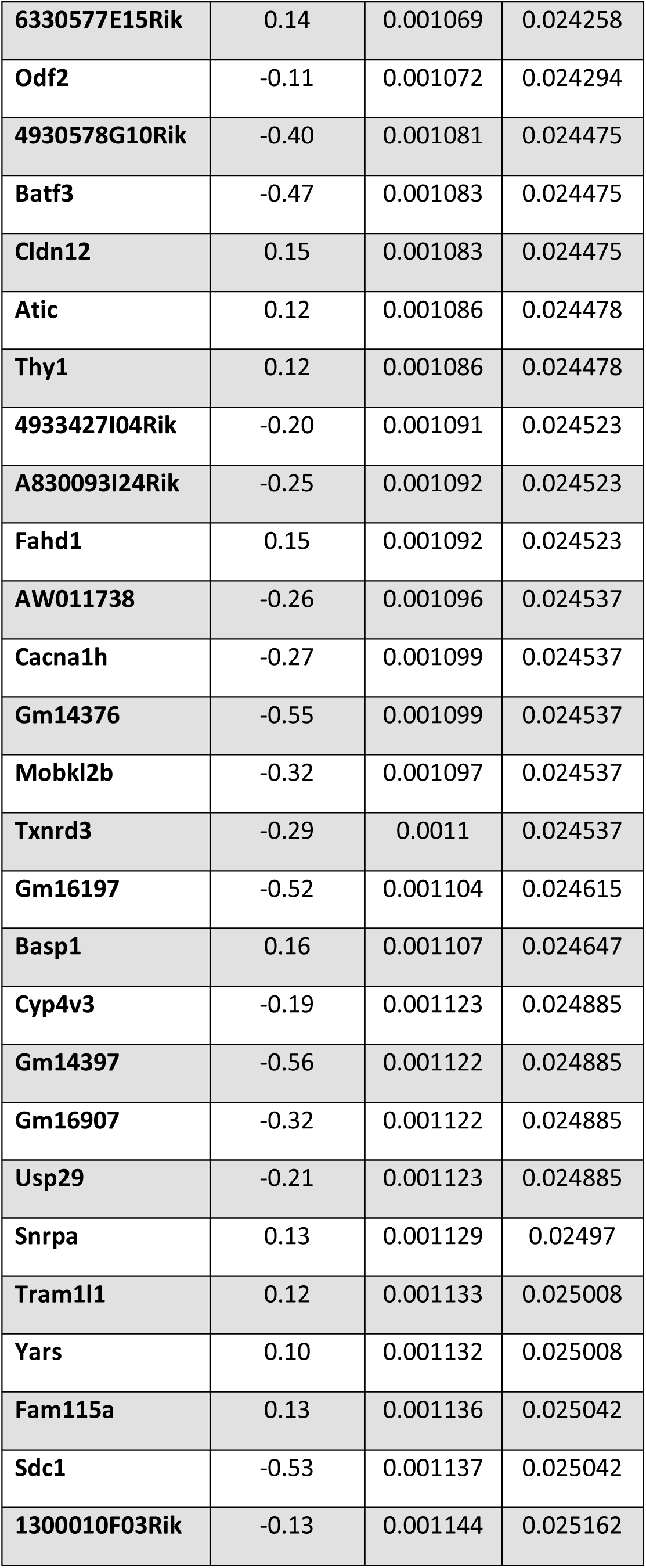
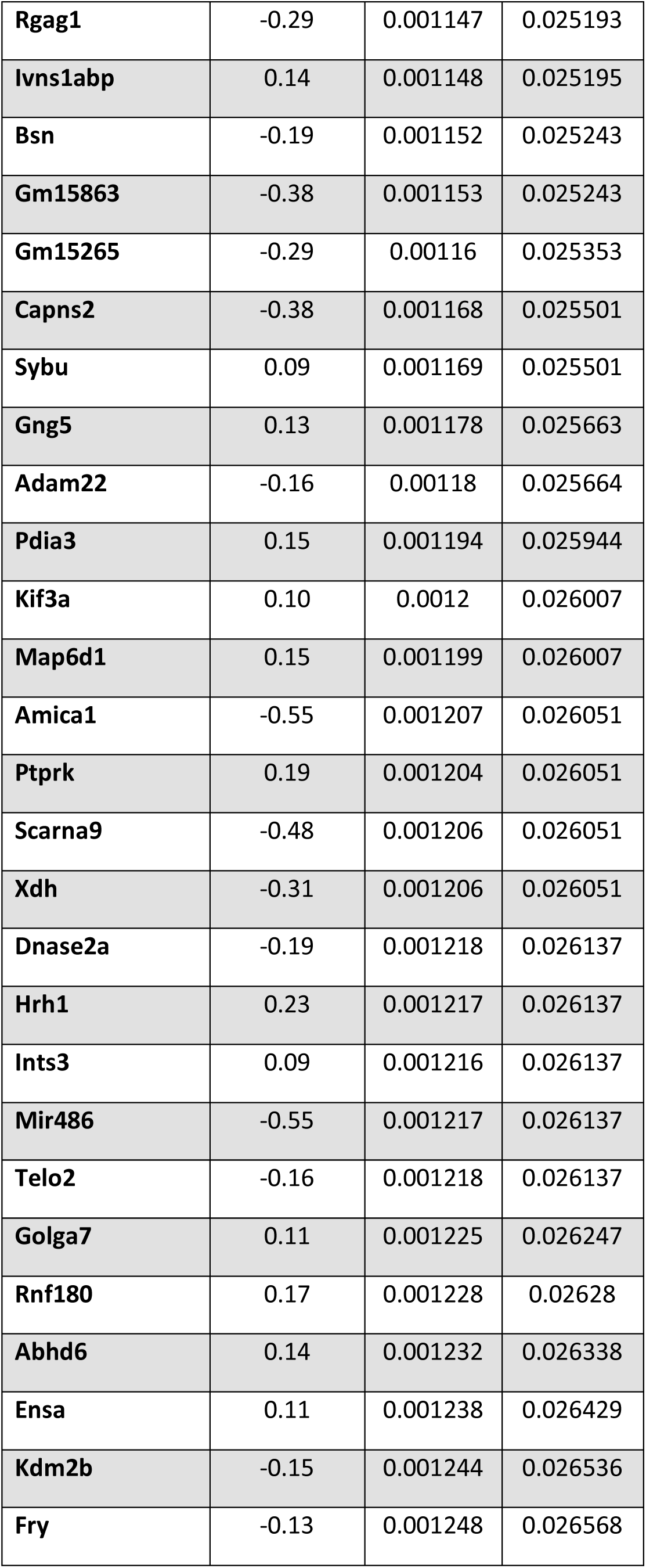
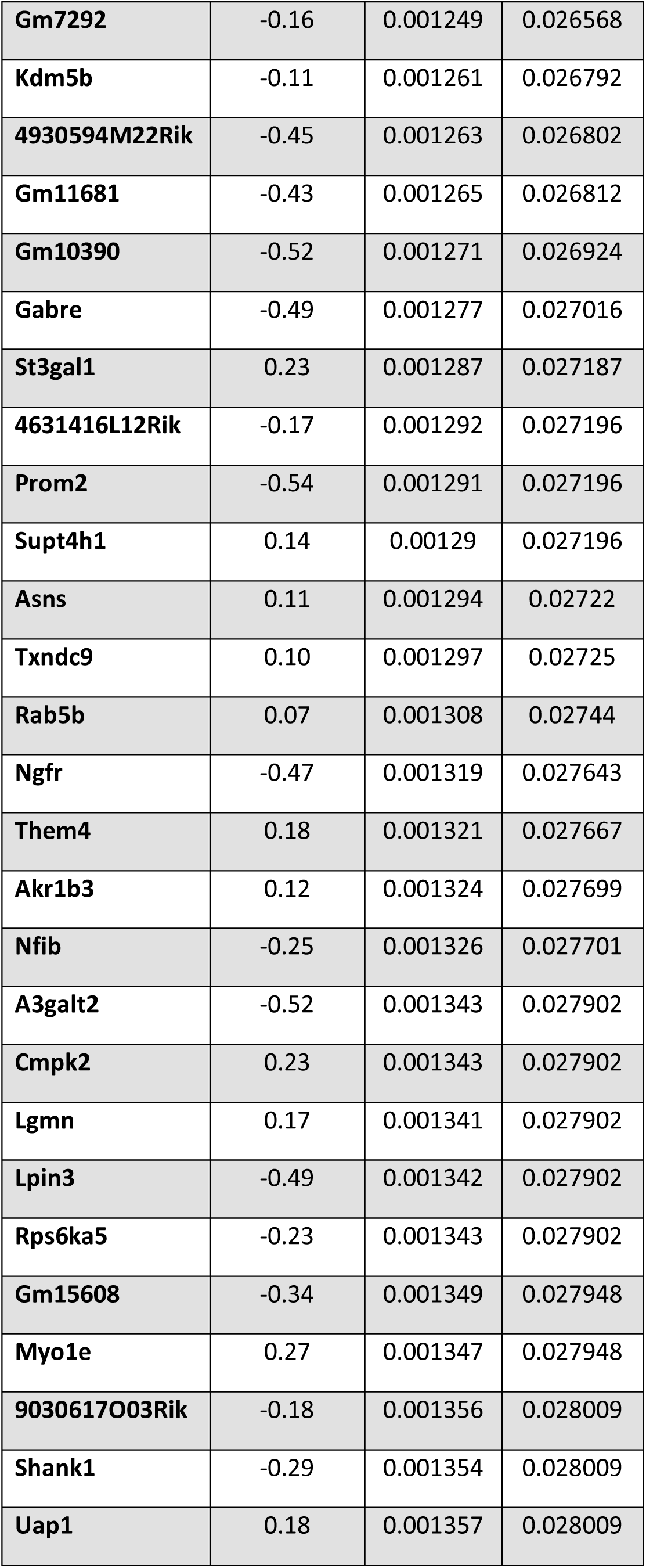
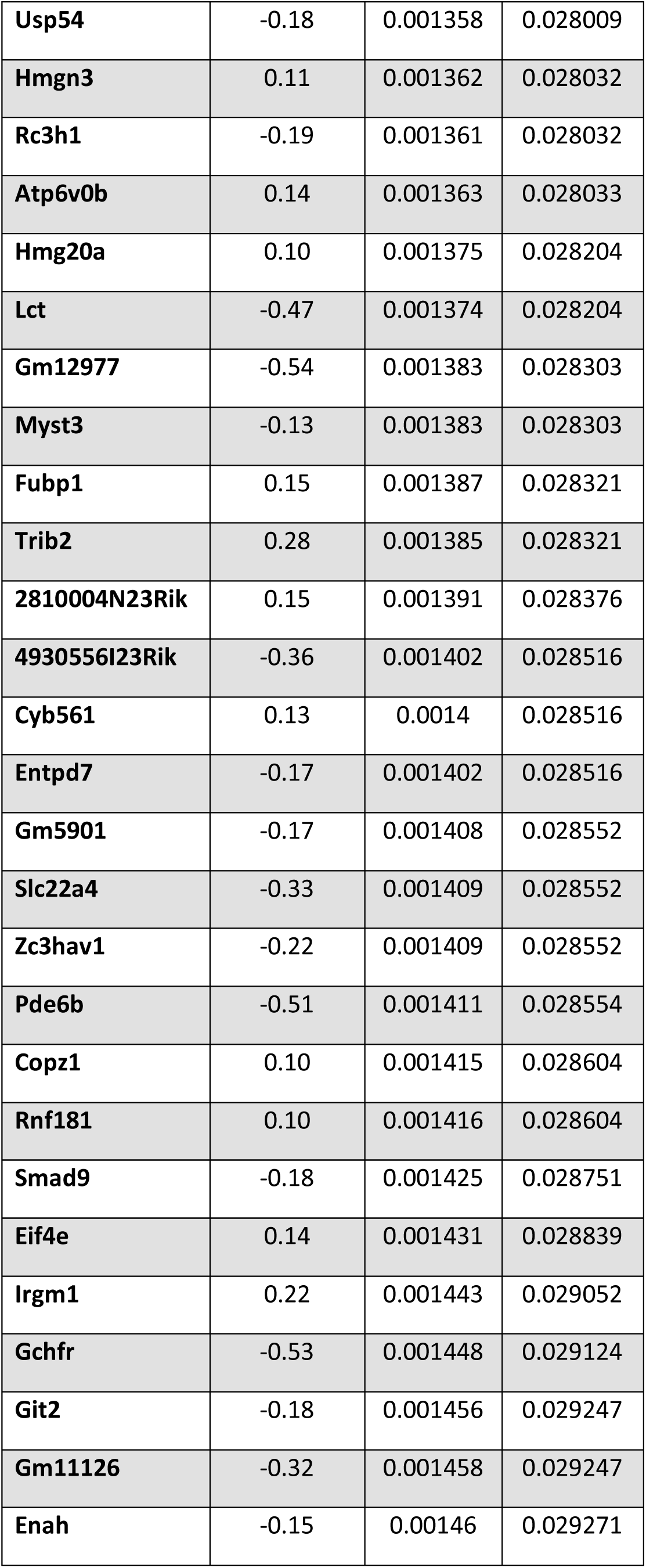
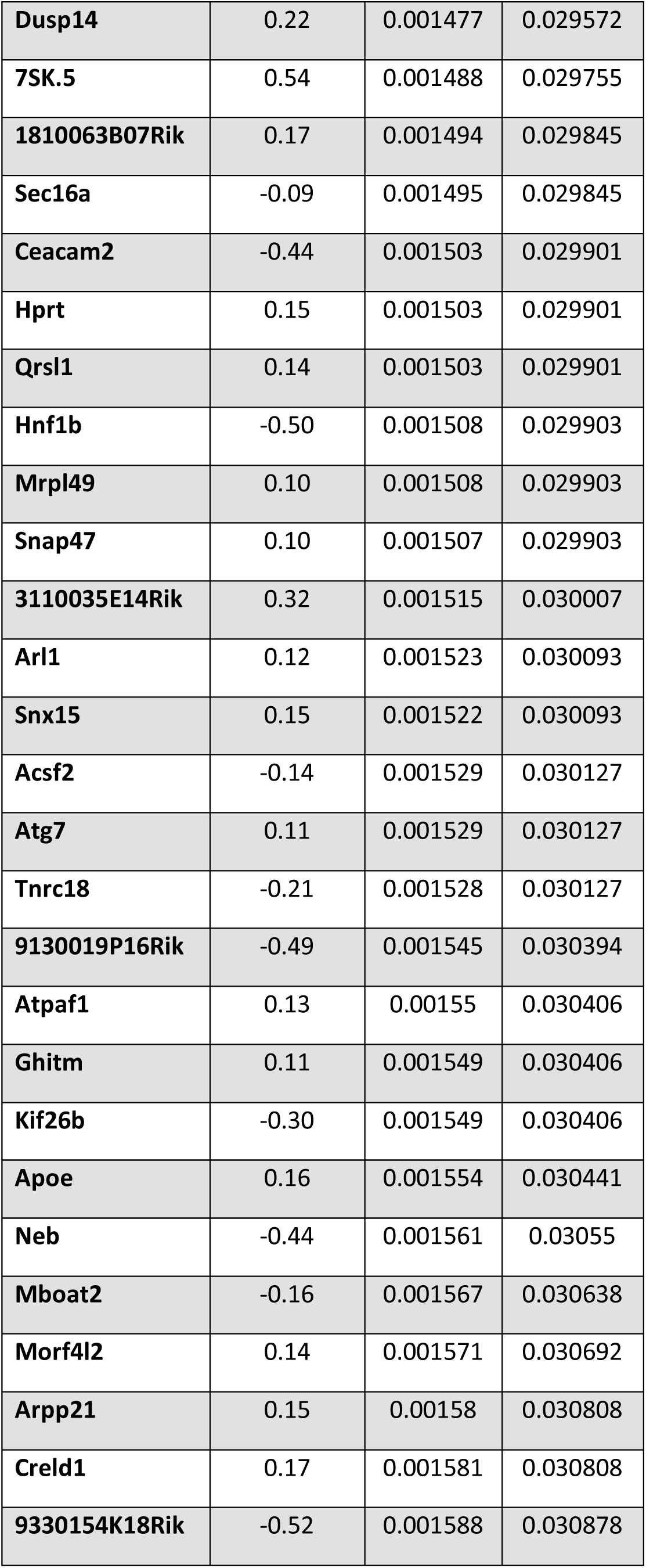
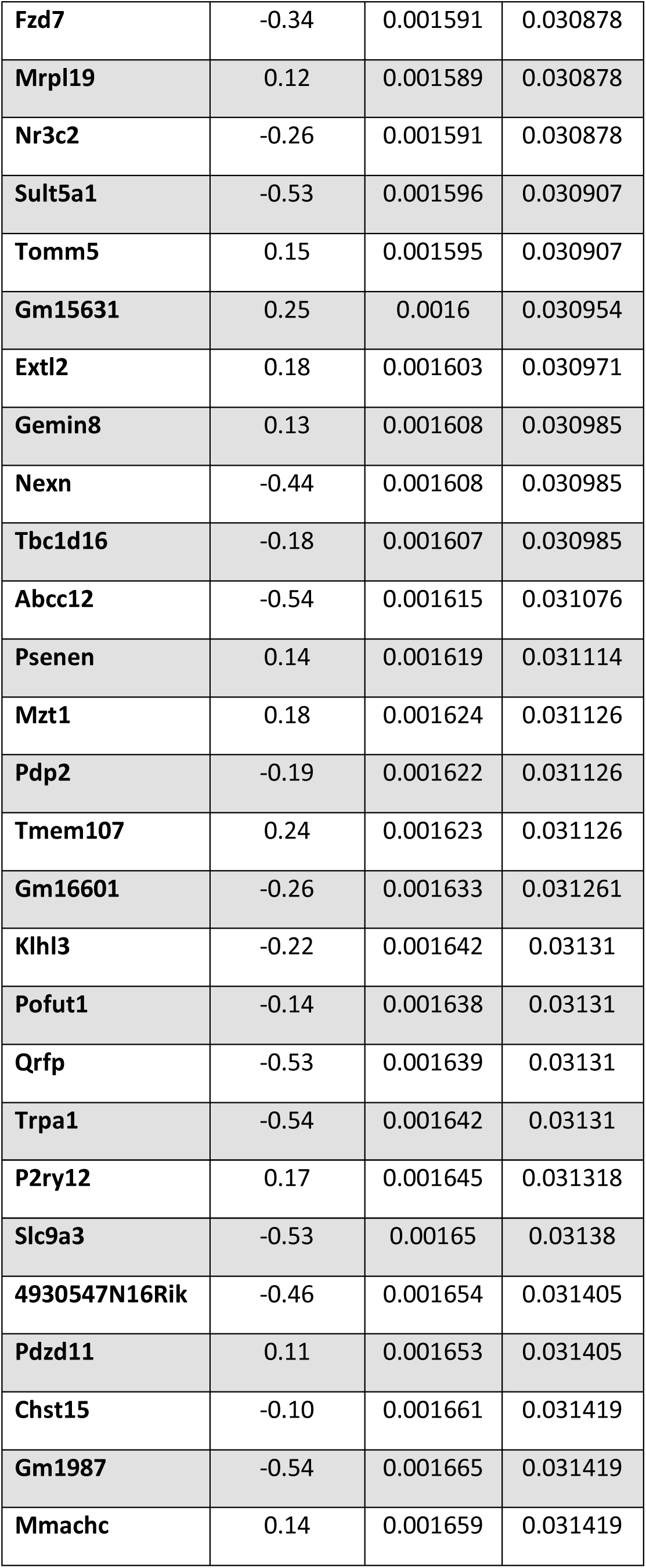
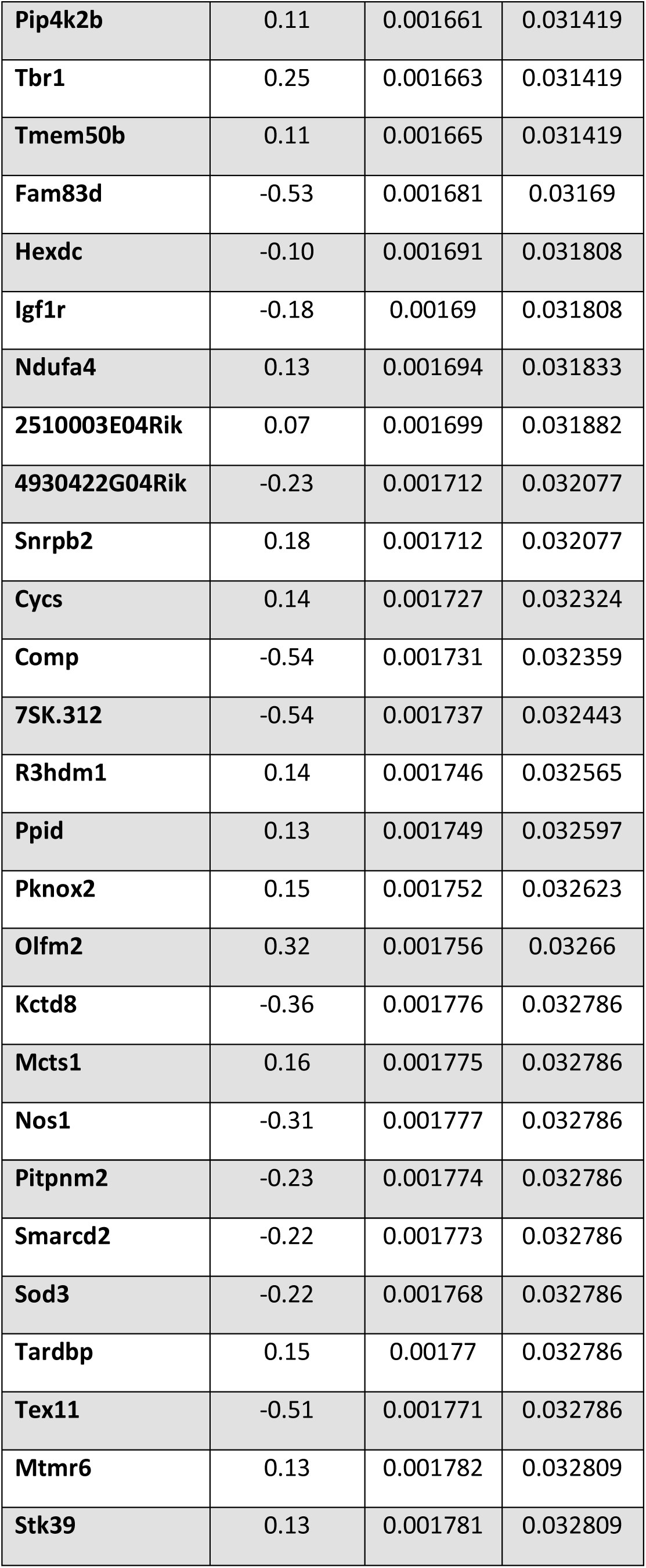
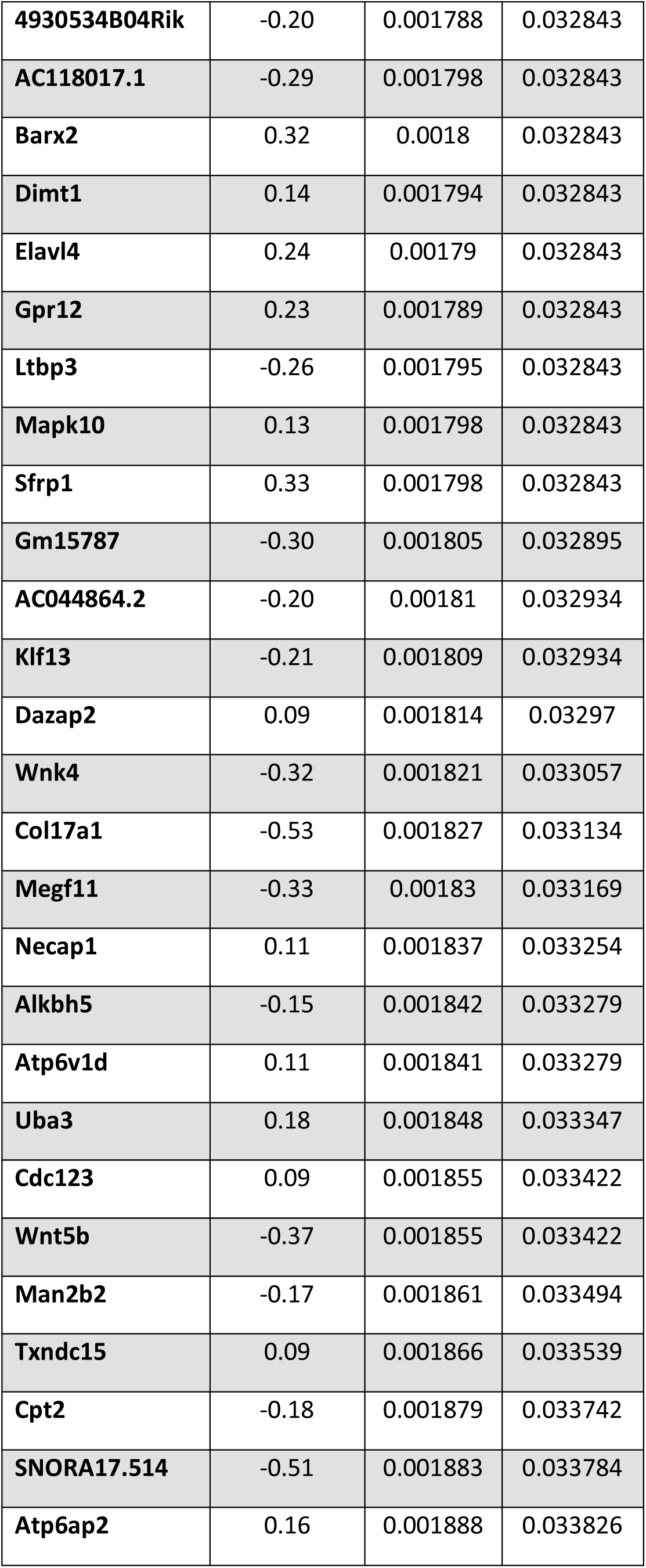
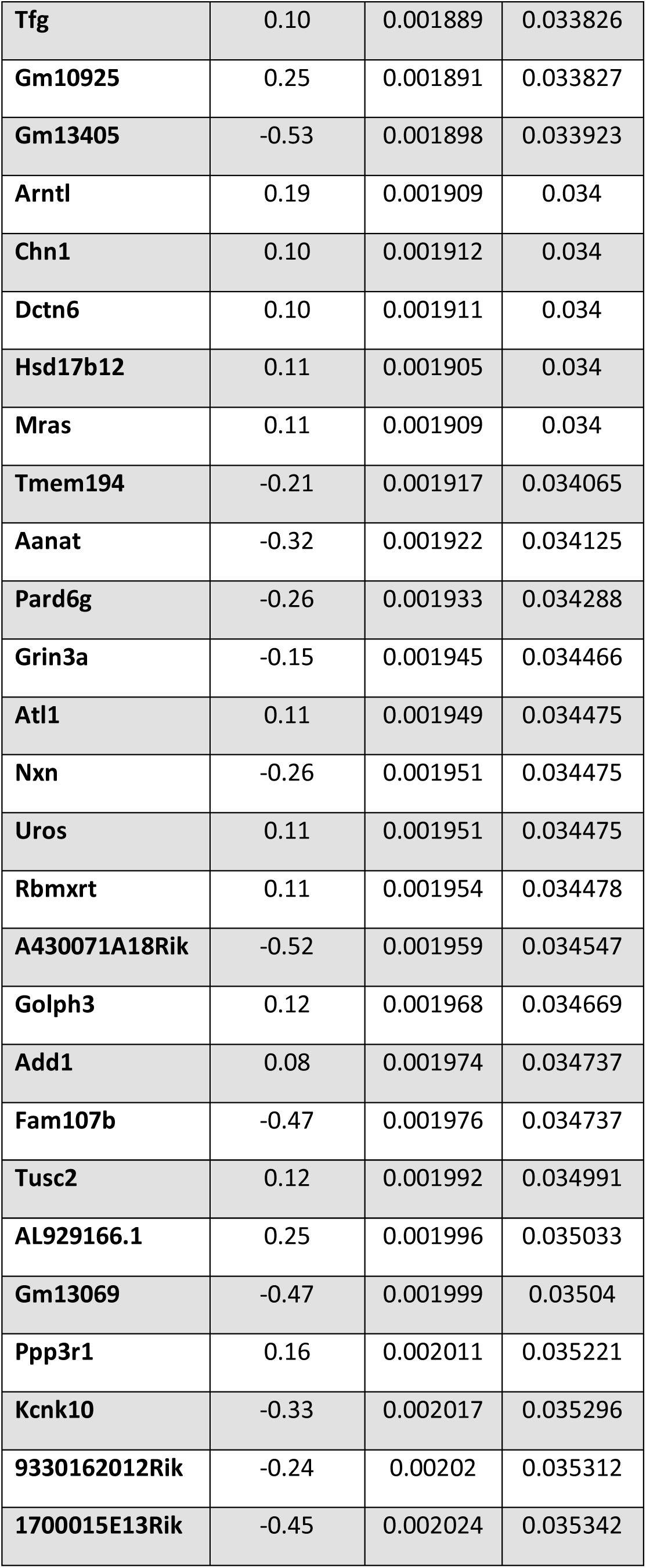
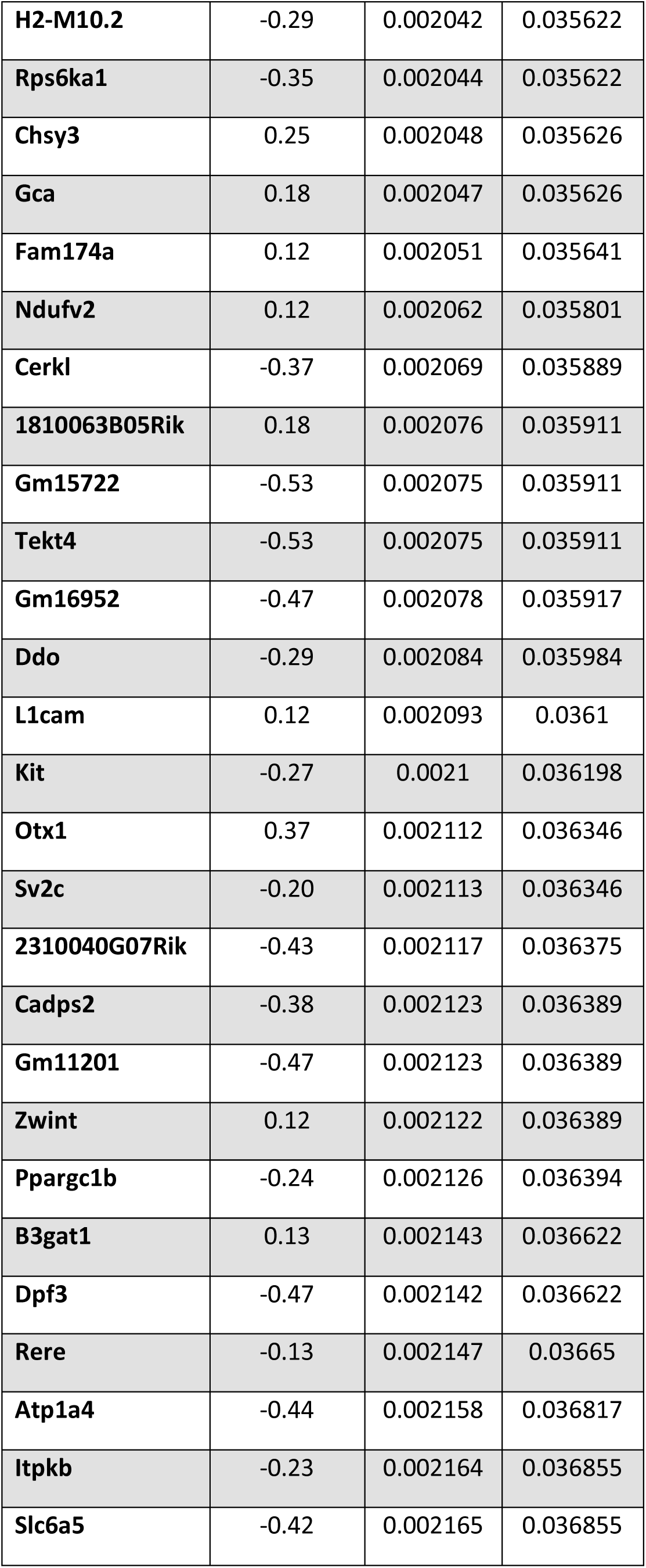
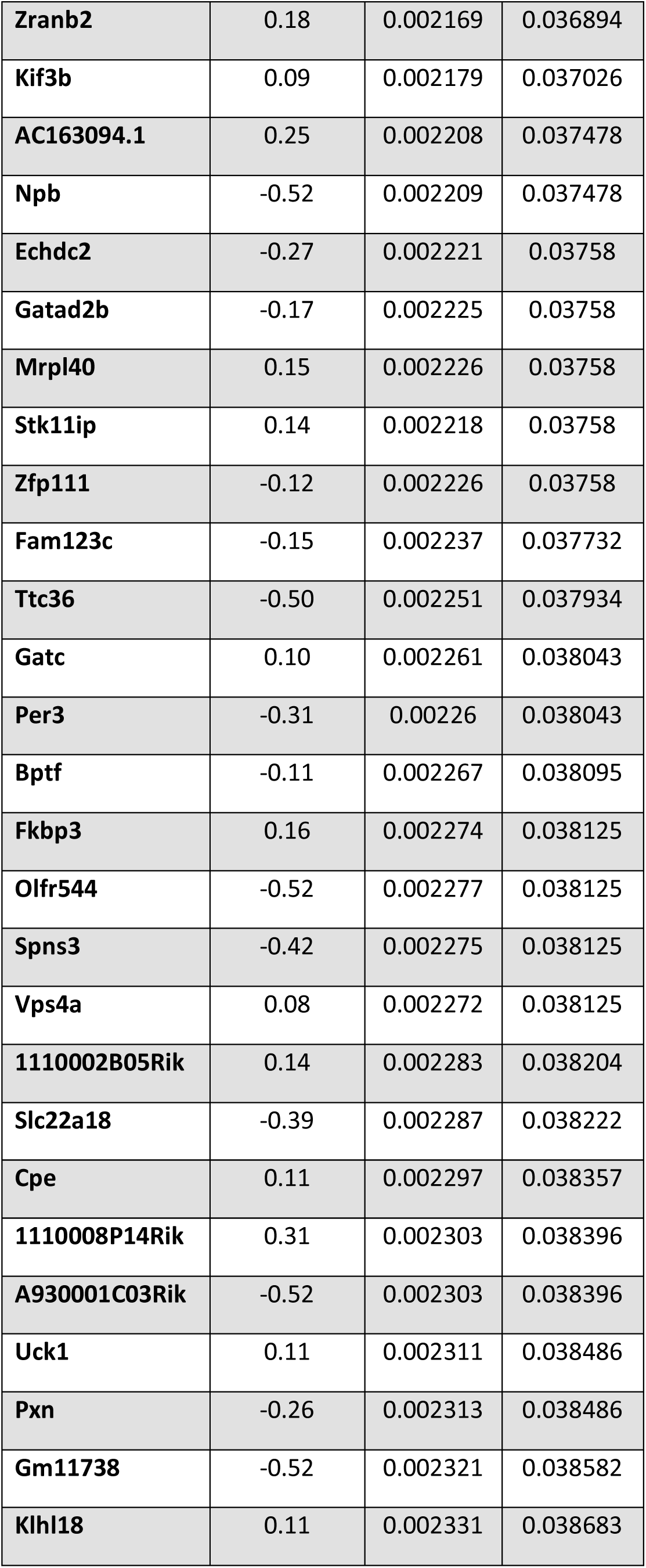
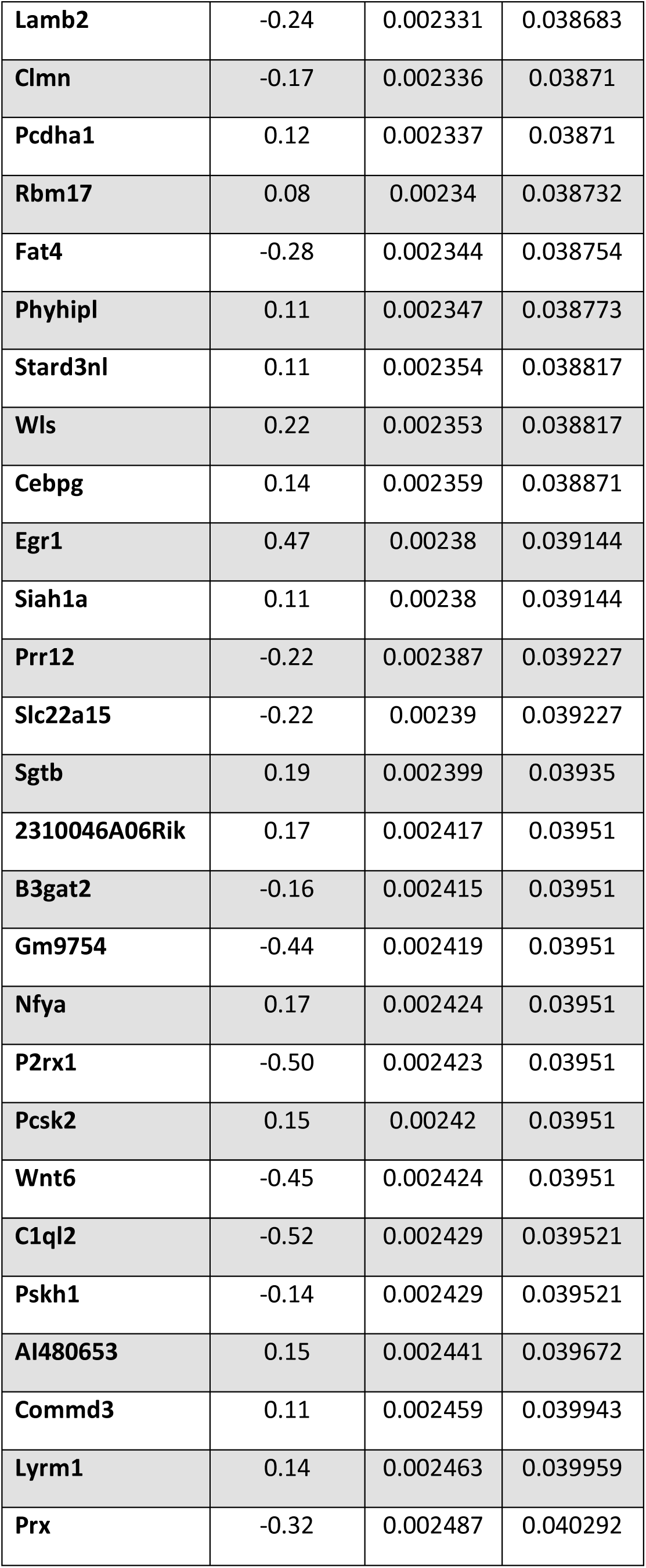
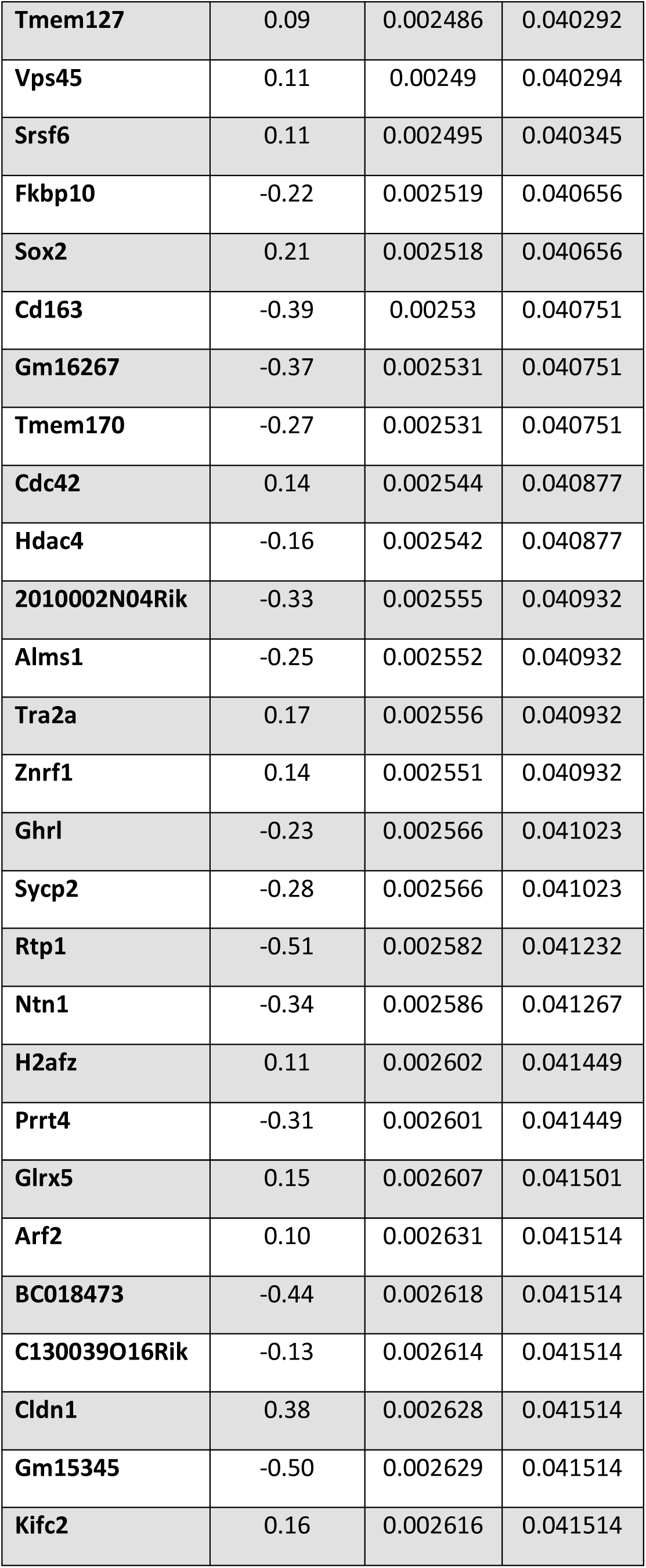
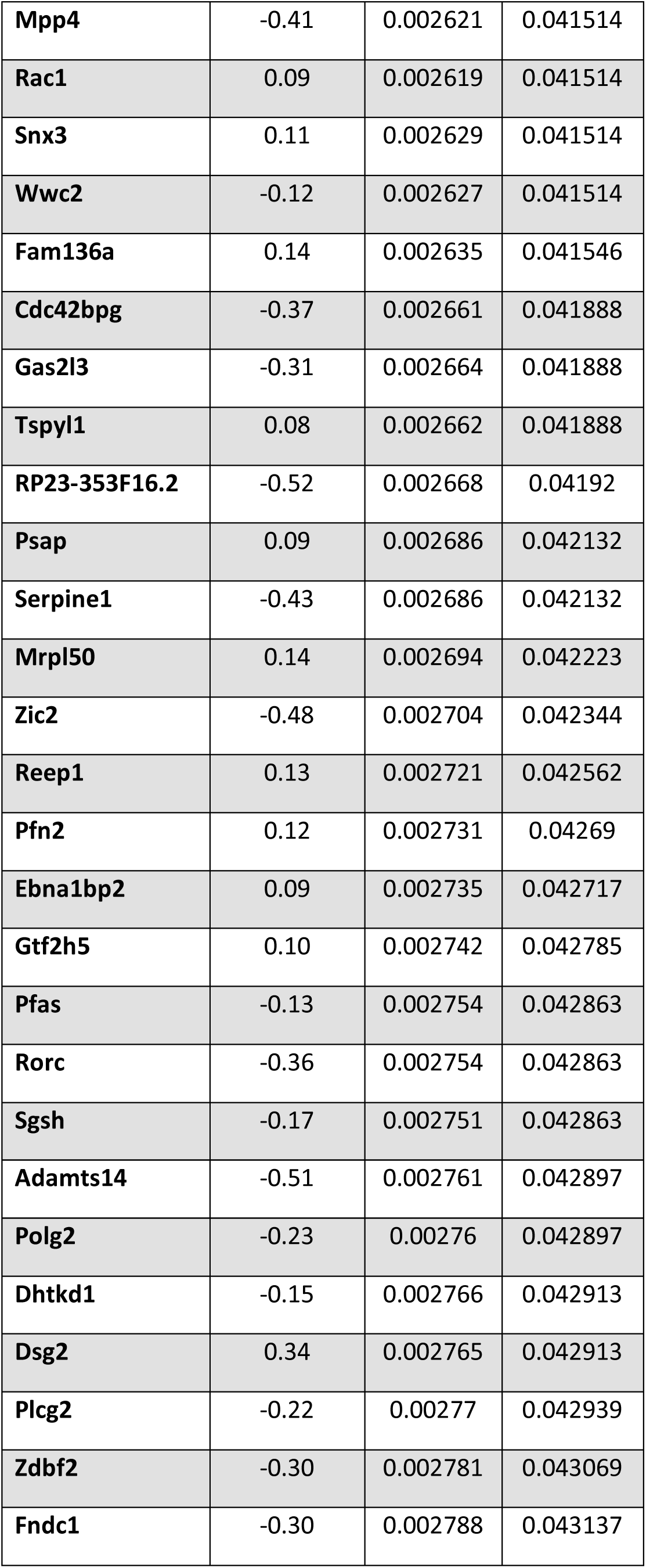
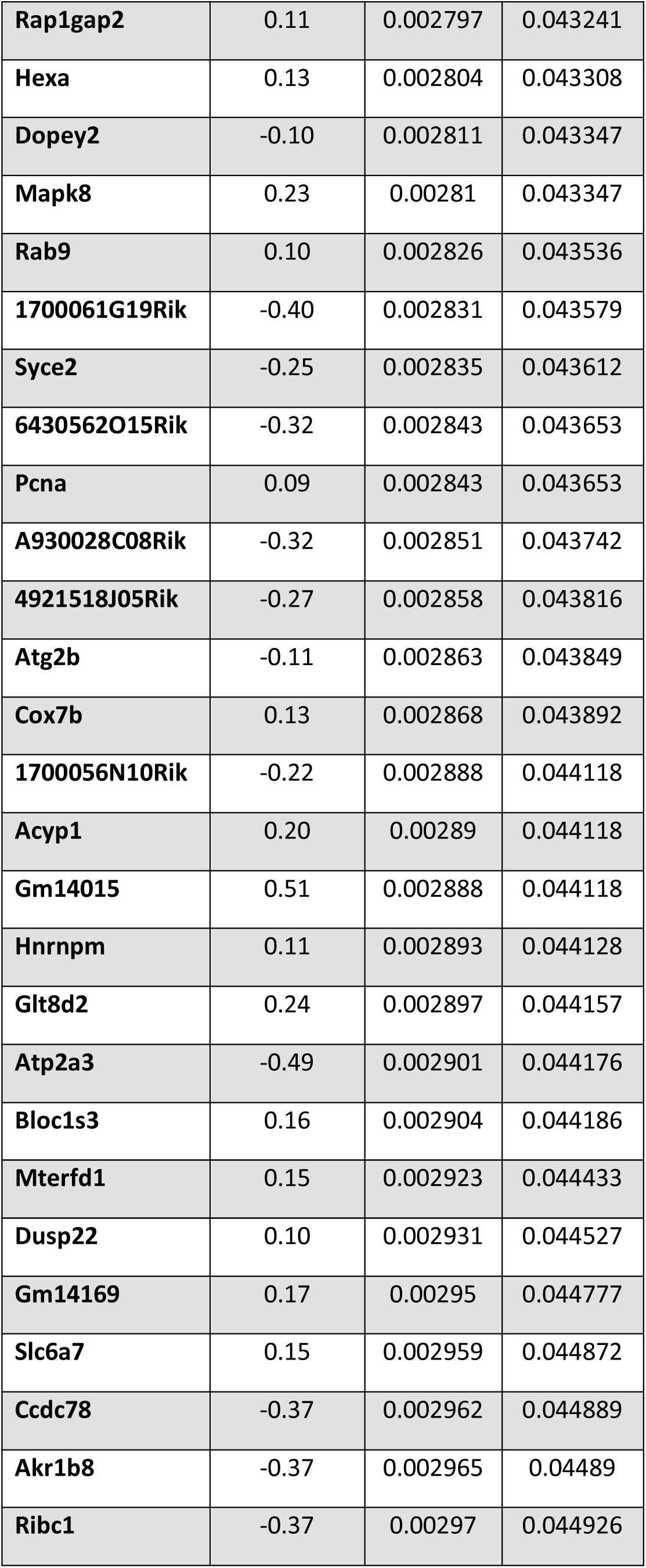
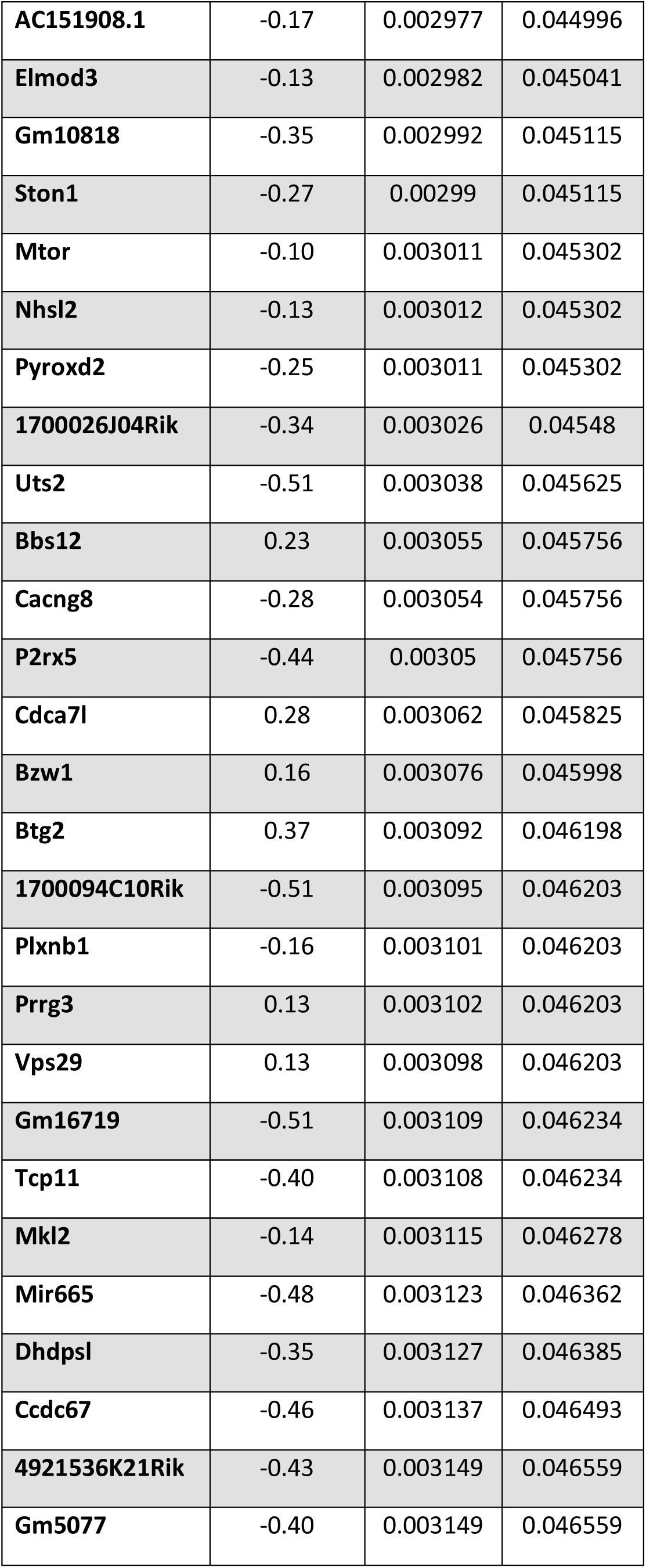
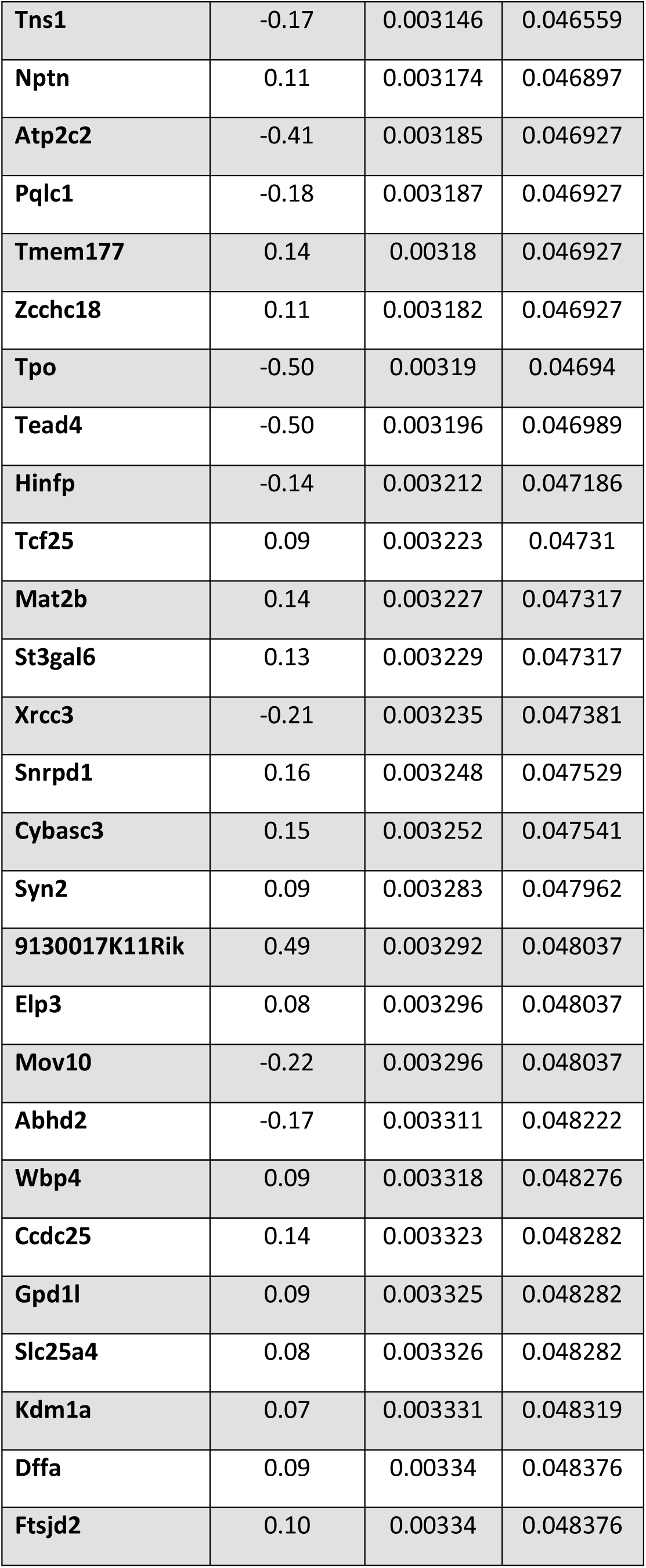
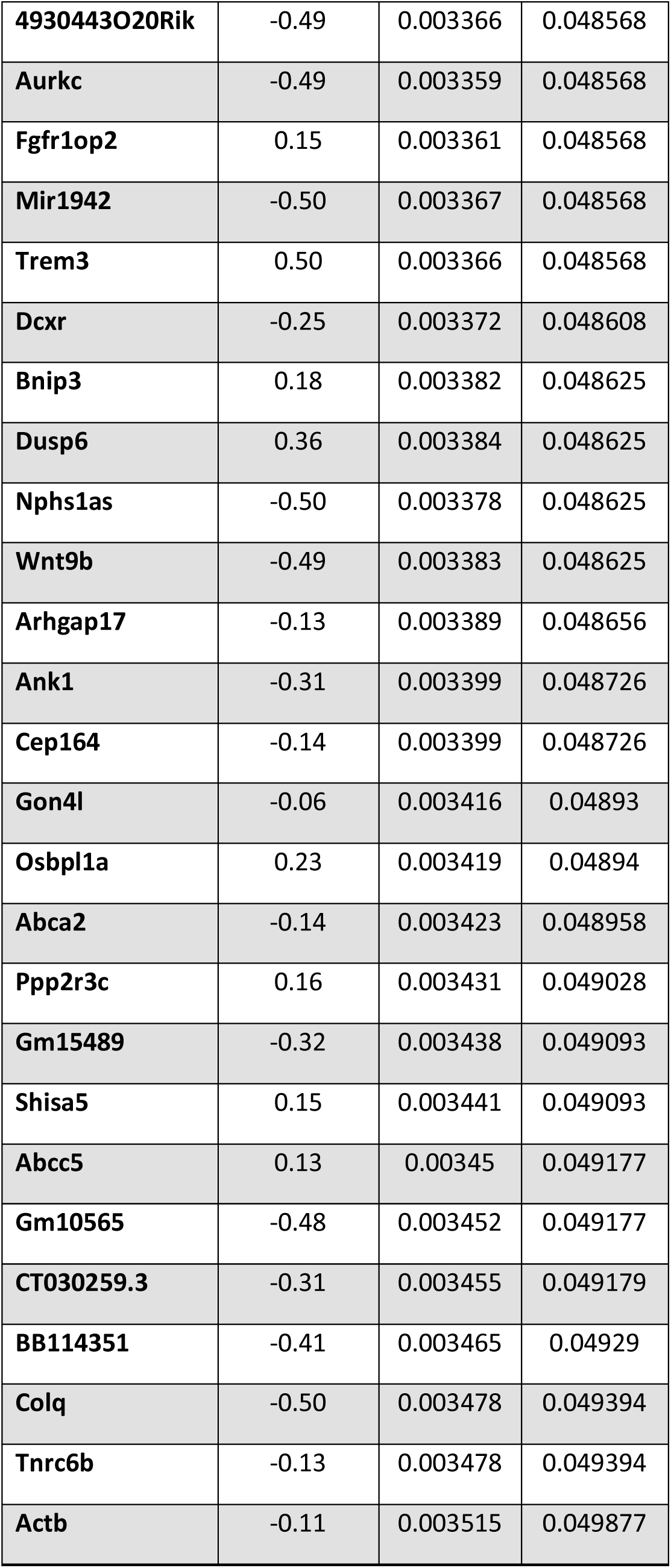
Differentially expressed genes from the EC of *APOE3/4* vs. *APOE3/3* mice

**Table S2.**
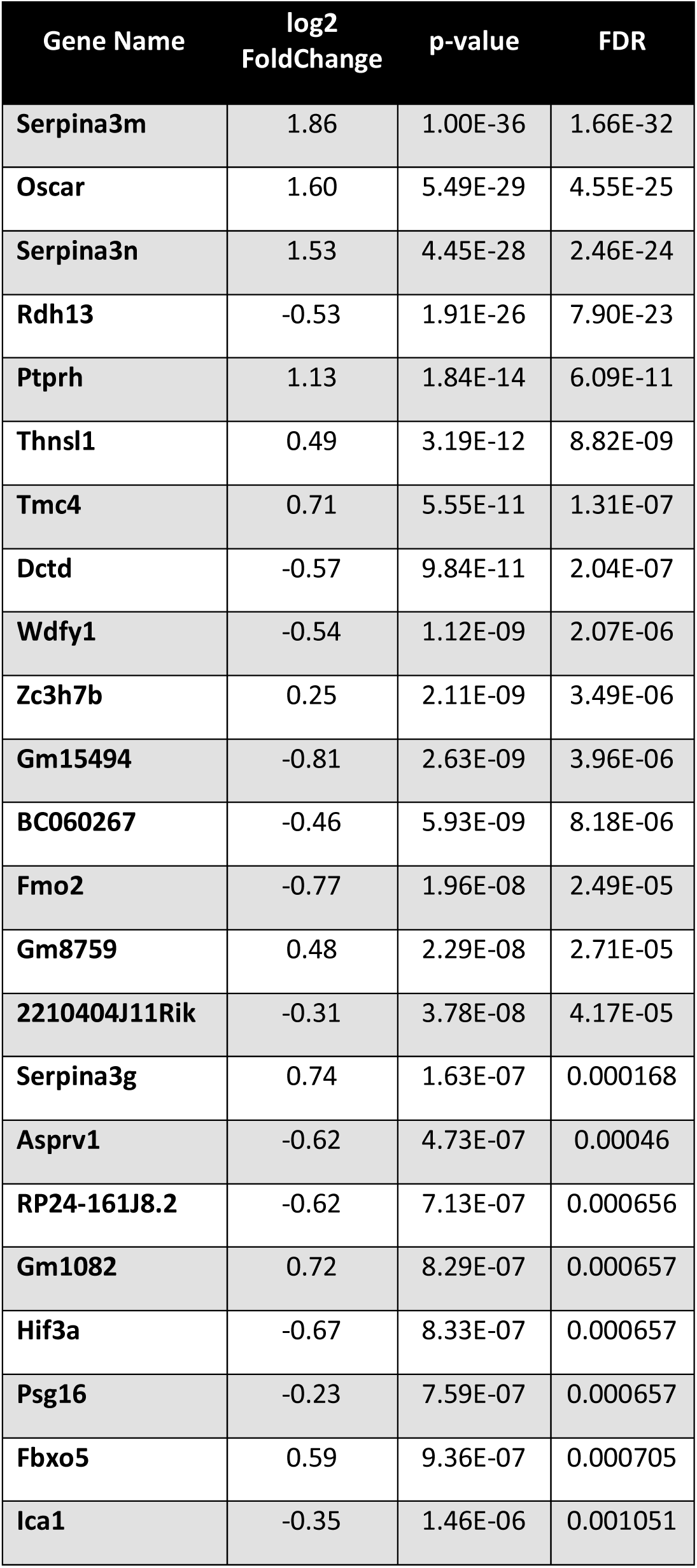
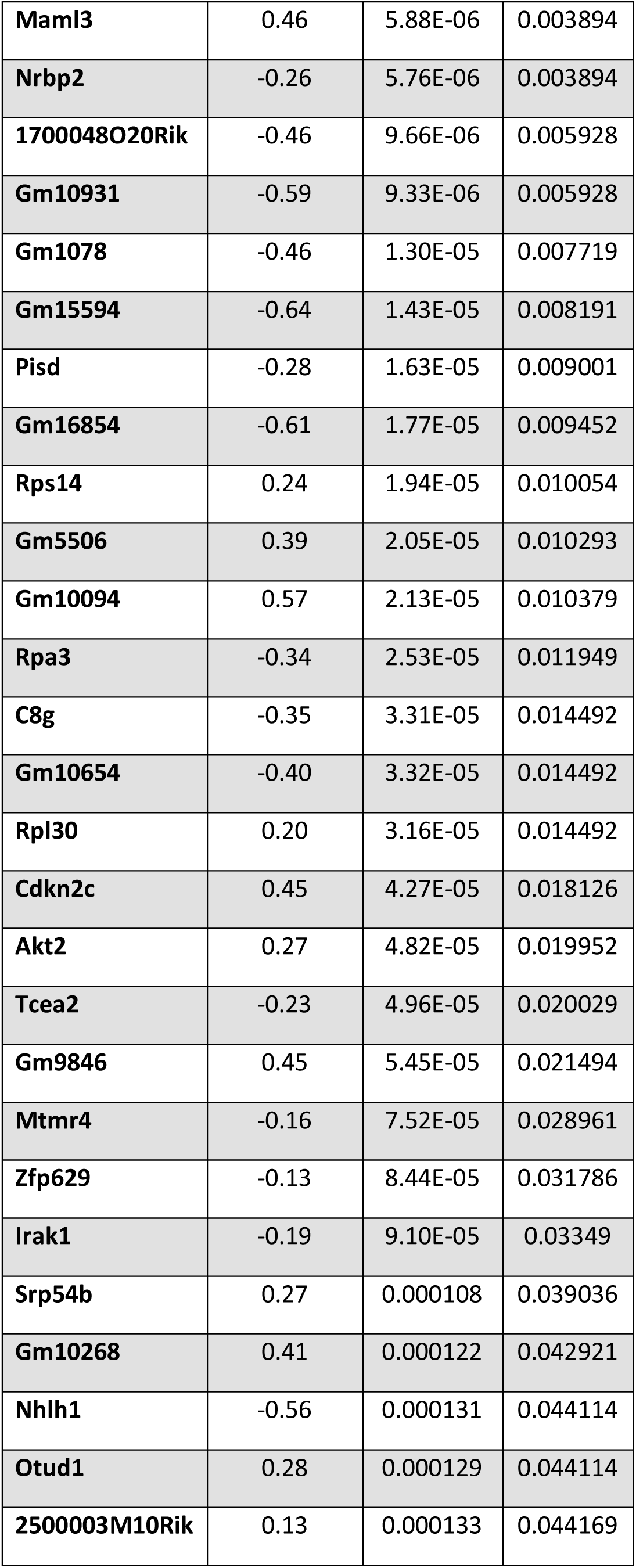
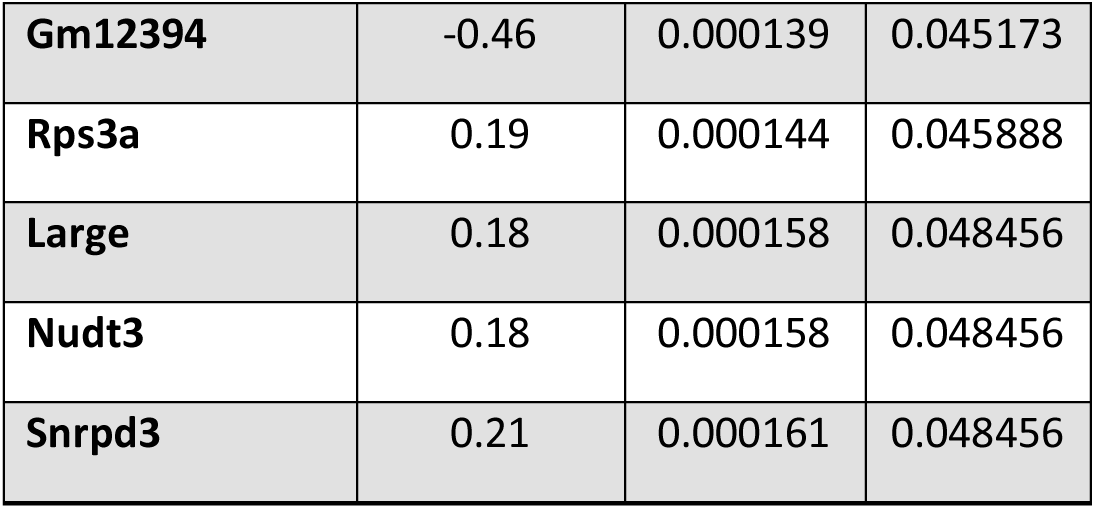
Differentially expressed genes from the PVC of *APOE3/4* vs. *APOE3/3* mice

**Table S3.**
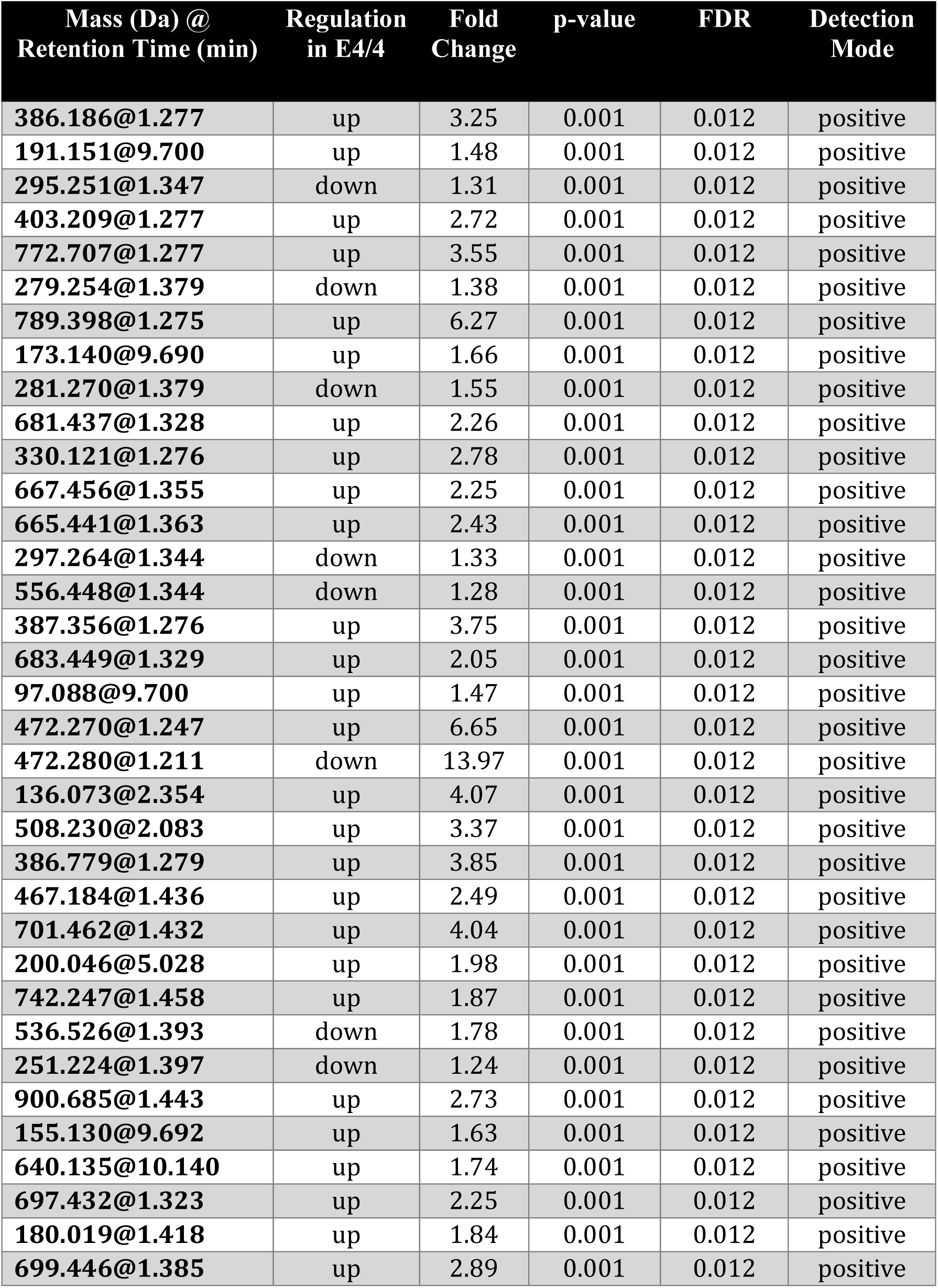
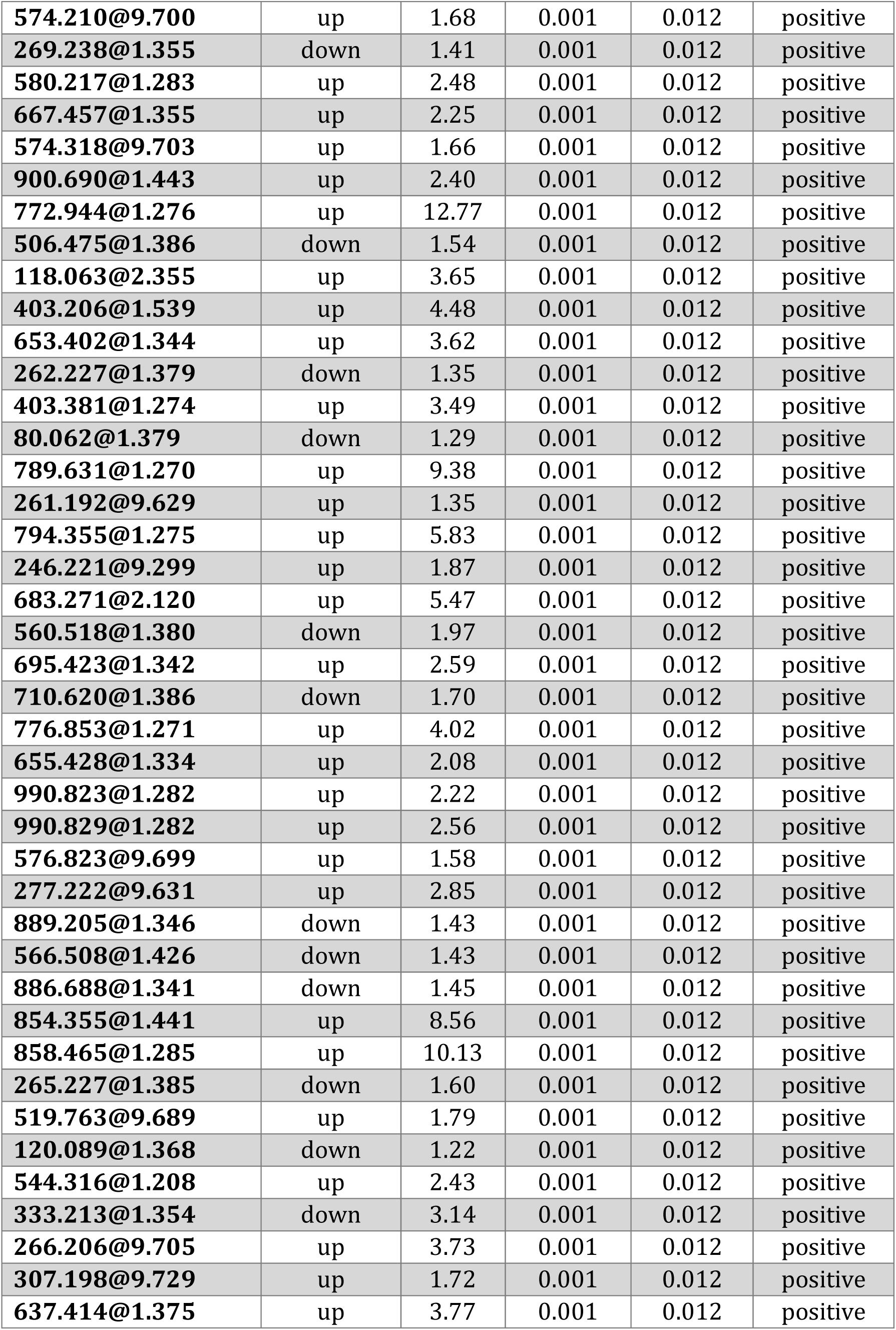
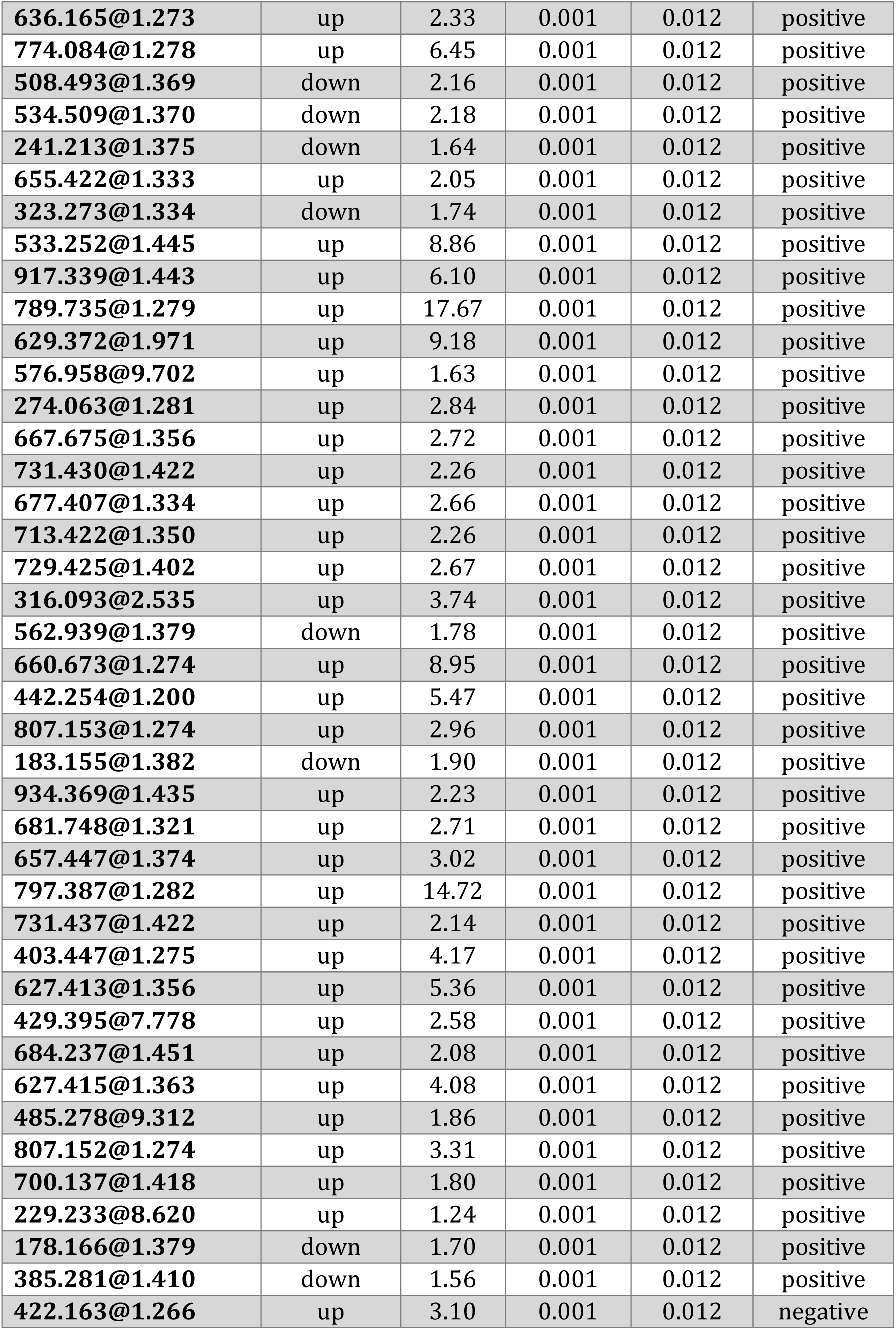
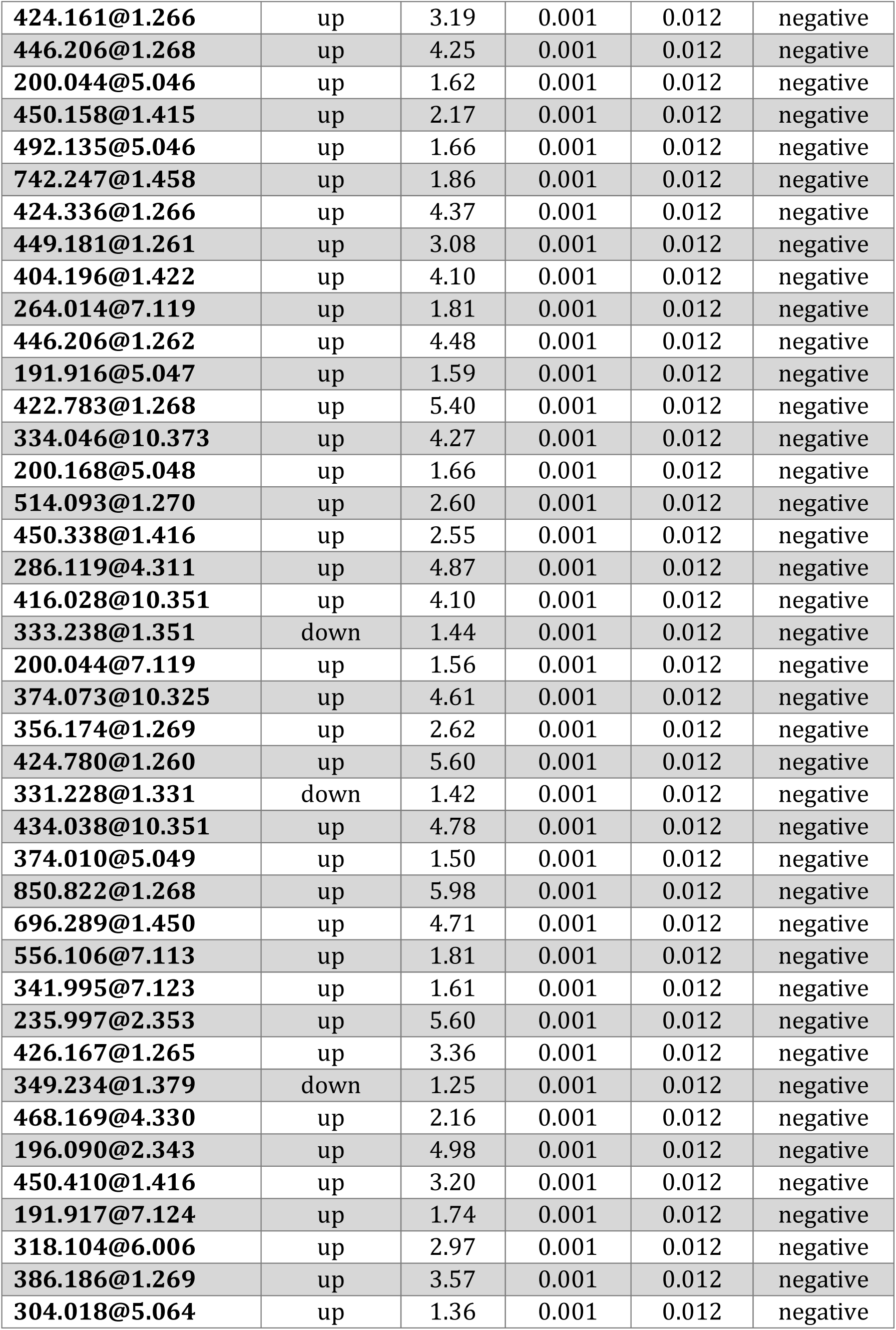
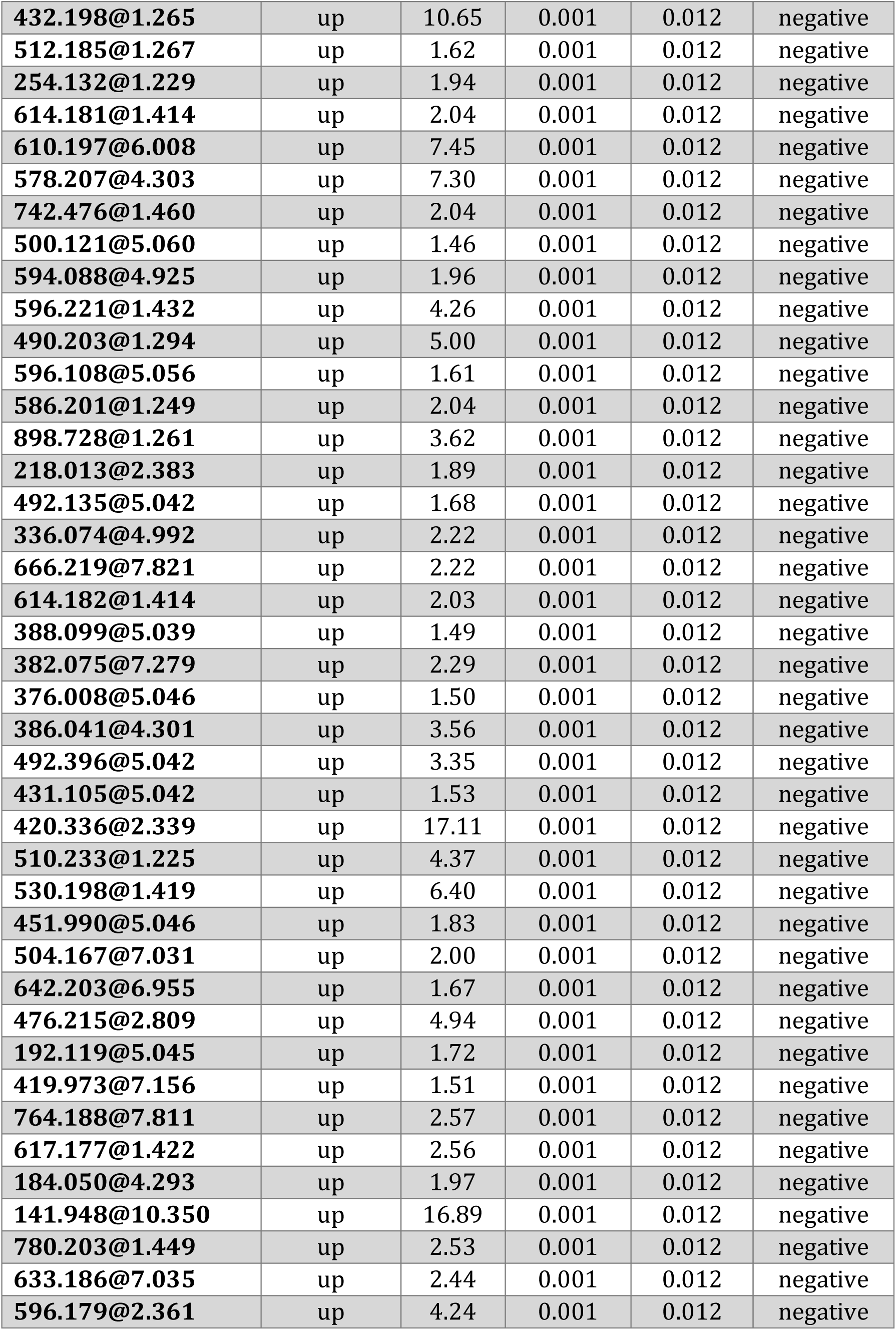
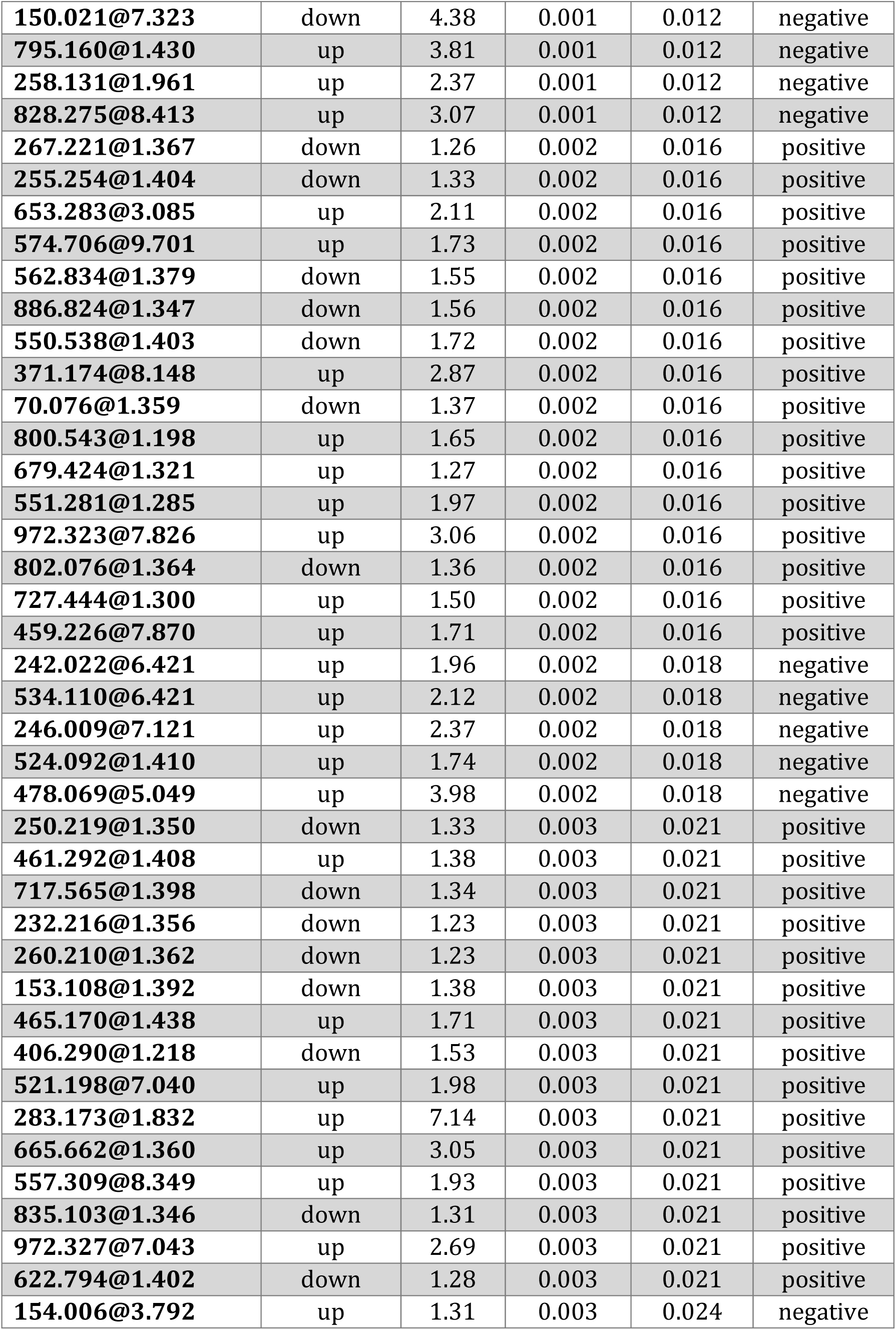
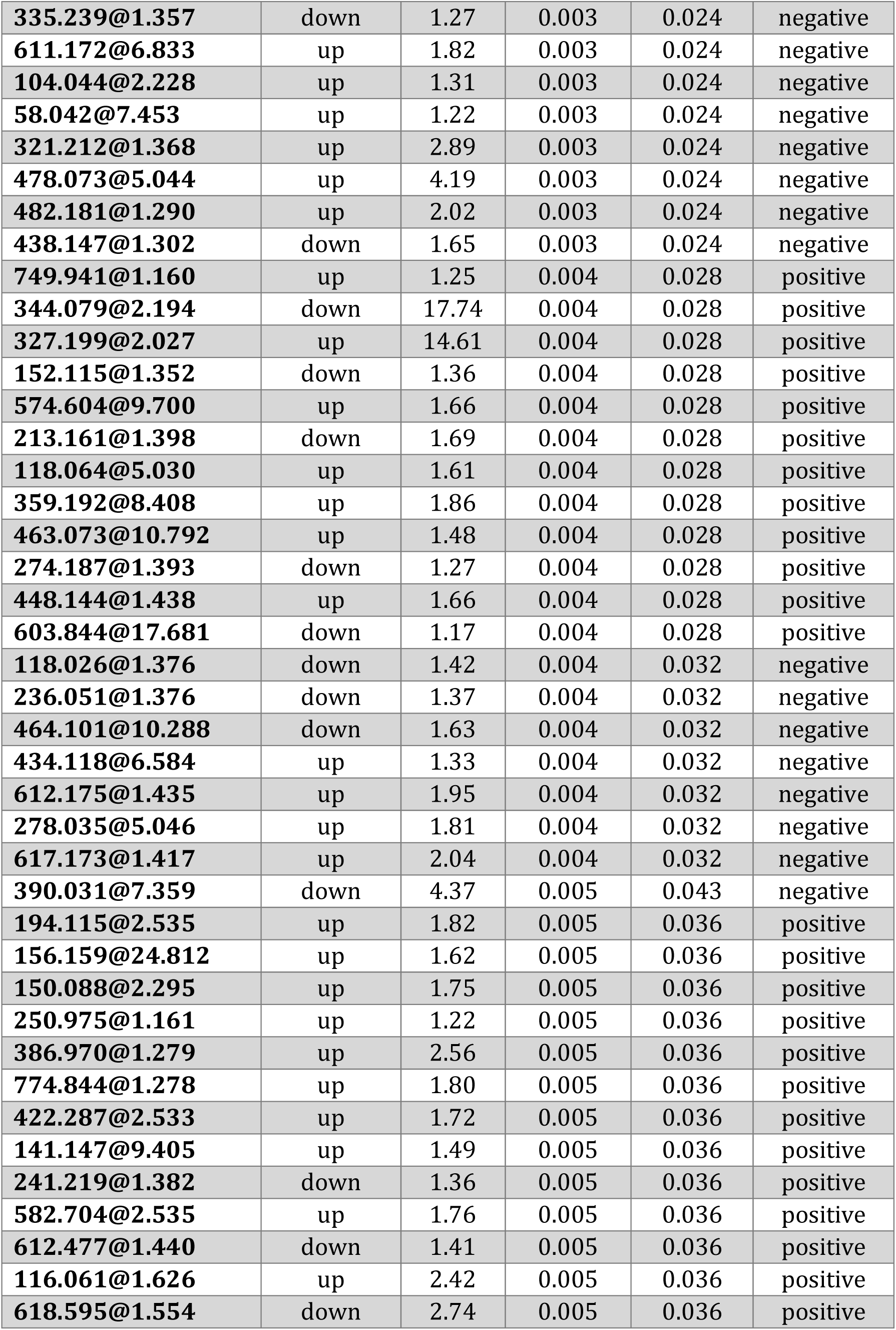
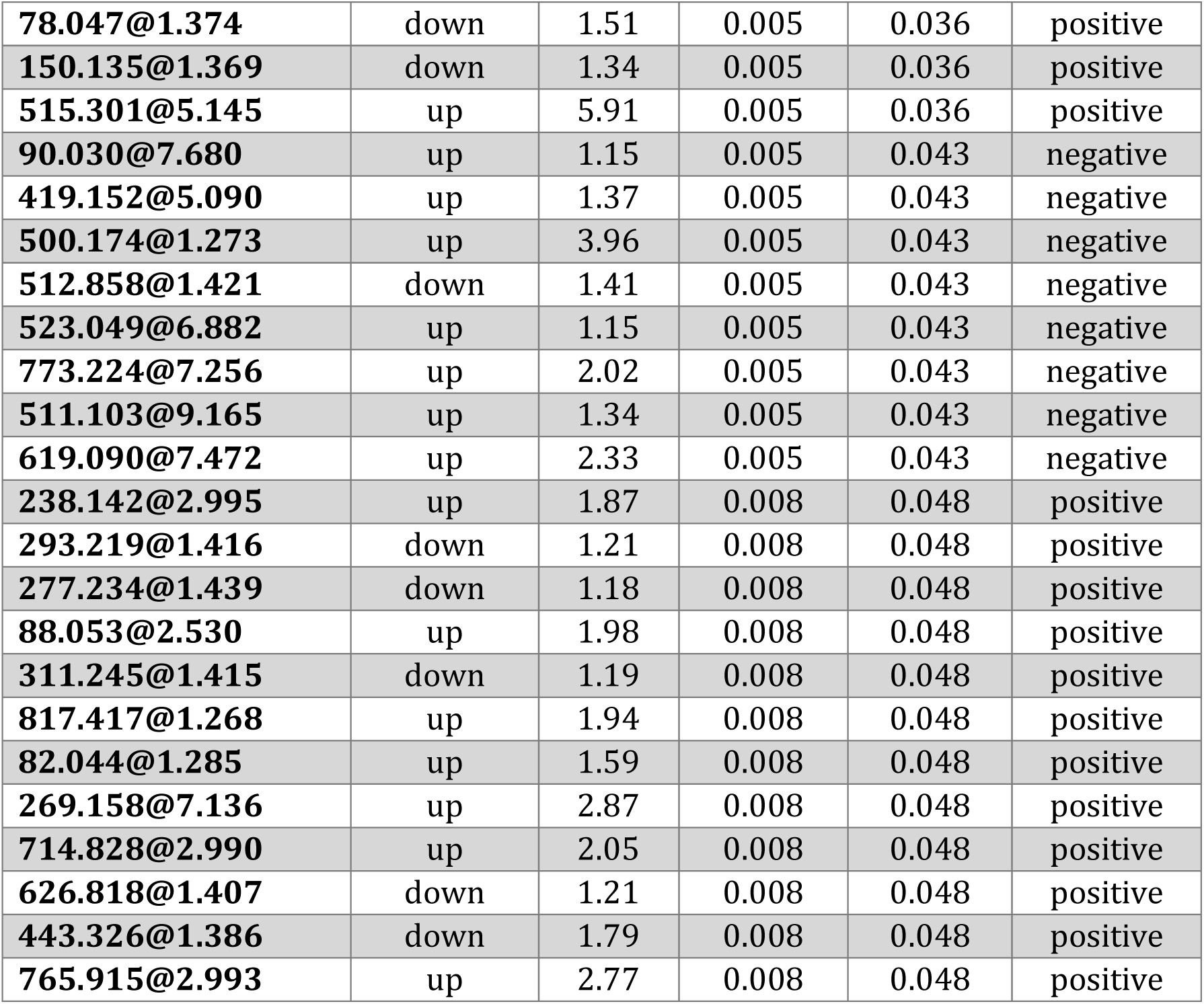
Differentially expressed untargeted metabolites from the EC of APOE3 vs. E4 mice

**Table S4.** Differentially expressed untargeted metabolites from the PVC of APOE3 vs. E4 mice

| **Mass (Da) @ Retention Time (min)** | **Regulation in E4/4** | **Fold Change** | **p-value** | **FDR** | **Detection Mode** |
| --- | --- | --- | --- | --- | --- |
| 386.186@1.277 | up | 3.11 | 0.001 | 0.014 | positive |
| 191.151@9.700 | up | 1.51 | 0.001 | 0.014 | positive |
| 403.209@1.277 | up | 2.64 | 0.001 | 0.014 | positive |
| 772.707@1.277 | up | 3.55 | 0.001 | 0.014 | positive |
| 279.254@1.379 | down | 1.37 | 0.001 | 0.014 | positive |
| 789.398@1.275 | up | 6.08 | 0.001 | 0.014 | positive |
| 173.140@9.690 | up | 1.64 | 0.001 | 0.014 | positive |
| 281.270@1.379 | down | 1.43 | 0.001 | 0.014 | positive |
| 681.437@1.328 | up | 2.29 | 0.001 | 0.014 | positive |
| 330.121@1.276 | up | 2.60 | 0.001 | 0.014 | positive |
| 667.456@1.355 | up | 2.10 | 0.001 | 0.014 | positive |
| 665.441@1.363 | up | 2.15 | 0.001 | 0.014 | positive |
| 297.264@1.344 | down | 1.30 | 0.001 | 0.014 | positive |
| 387.356@1.276 | up | 3.69 | 0.001 | 0.014 | positive |
| 683.449@1.329 | up | 2.02 | 0.001 | 0.014 | positive |
| 97.088@9.700 | up | 1.51 | 0.001 | 0.014 | positive |
| 472.270@1.247 | up | 20.46 | 0.001 | 0.014 | positive |
| 810.327@1.275 | up | 5.15 | 0.001 | 0.014 | positive |
| 472.280@1.211 | down | 12.38 | 0.001 | 0.014 | positive |
| 255.254@1.404 | down | 1.27 | 0.001 | 0.014 | positive |
| 136.073@2.354 | up | 2.90 | 0.001 | 0.014 | positive |
| 508.230@2.083 | up | 3.89 | 0.001 | 0.014 | positive |
| 386.779@1.279 | up | 4.48 | 0.001 | 0.014 | positive |
| 467.184@1.436 | up | 2.81 | 0.001 | 0.014 | positive |
| 424.141@1.280 | up | 2.13 | 0.001 | 0.014 | positive |
| 701.462@1.432 | up | 3.64 | 0.001 | 0.014 | positive |
| 174.008@1.273 | up | 1.94 | 0.001 | 0.014 | positive |
| 536.526@1.393 | down | 1.59 | 0.001 | 0.014 | positive |
| 251.224@1.397 | down | 1.28 | 0.001 | 0.014 | positive |
| 900.685@1.443 | up | 8.45 | 0.001 | 0.014 | positive |
| 155.130@9.692 | up | 1.60 | 0.001 | 0.014 | positive |
| 697.432@1.323 | up | 2.17 | 0.001 | 0.014 | positive |
| 180.019@1.418 | up | 1.64 | 0.001 | 0.014 | positive |
| 699.446@1.385 | up | 2.57 | 0.001 | 0.014 | positive |
| 574.210@9.700 | up | 1.74 | 0.001 | 0.014 | positive |
| 269.238@1.355 | down | 1.36 | 0.001 | 0.014 | positive |
| 580.217@1.283 | up | 2.50 | 0.001 | 0.014 | positive |
| 250.975@1.161 | up | 1.14 | 0.001 | 0.014 | positive |
| 667.457@1.355 | up | 2.10 | 0.001 | 0.014 | positive |
| 574.318@9.703 | up | 1.68 | 0.001 | 0.014 | positive |
| 900.690@1.443 | up | 5.28 | 0.001 | 0.014 | positive |
| 772.944@1.276 | up | 10.49 | 0.001 | 0.014 | positive |
| 506.475@1.386 | down | 1.51 | 0.001 | 0.014 | positive |
| 118.063@2.355 | up | 2.75 | 0.001 | 0.014 | positive |
| 262.227@1.379 | down | 1.29 | 0.001 | 0.014 | positive |
| 403.381@1.274 | up | 3.62 | 0.001 | 0.014 | positive |
| 789.631@1.270 | up | 5.61 | 0.001 | 0.014 | positive |
| 246.221@9.299 | up | 1.76 | 0.001 | 0.014 | positive |
| 683.271@2.120 | up | 4.25 | 0.001 | 0.014 | positive |
| 560.518@1.380 | down | 1.96 | 0.001 | 0.014 | positive |
| 695.423@1.342 | up | 1.72 | 0.001 | 0.014 | positive |
| 710.620@1.386 | down | 2.26 | 0.001 | 0.014 | positive |
| 776.853@1.271 | up | 4.51 | 0.001 | 0.014 | positive |
| 655.428@1.334 | up | 1.81 | 0.001 | 0.014 | positive |
| 990.823@1.282 | up | 2.14 | 0.001 | 0.014 | positive |
| 574.706@9.701 | up | 1.70 | 0.001 | 0.014 | positive |
| 990.829@1.282 | up | 2.29 | 0.001 | 0.014 | positive |
| 576.823@9.699 | up | 1.61 | 0.001 | 0.014 | positive |
| 277.222@9.631 | up | 2.74 | 0.001 | 0.014 | positive |
| 566.508@1.426 | down | 1.53 | 0.001 | 0.014 | positive |
| 886.688@1.341 | down | 1.40 | 0.001 | 0.014 | positive |
| 854.355@1.441 | up | 6.49 | 0.001 | 0.014 | positive |
| 858.465@1.285 | up | 10.18 | 0.001 | 0.014 | positive |
| 562.834@1.379 | down | 1.54 | 0.001 | 0.014 | positive |
| 544.316@1.208 | up | 2.70 | 0.001 | 0.014 | positive |
| 266.206@9.705 | up | 2.65 | 0.001 | 0.014 | positive |
| 636.165@1.273 | up | 2.09 | 0.001 | 0.014 | positive |
| 241.219@1.382 | down | 1.27 | 0.001 | 0.014 | positive |
| 774.084@1.278 | up | 7.49 | 0.001 | 0.014 | positive |
| 508.493@1.369 | down | 2.10 | 0.001 | 0.014 | positive |
| 534.509@1.370 | down | 2.07 | 0.001 | 0.014 | positive |
| 655.422@1.333 | up | 1.80 | 0.001 | 0.014 | positive |
| 917.339@1.443 | up | 8.14 | 0.001 | 0.014 | positive |
| 789.735@1.279 | up | 14.76 | 0.001 | 0.014 | positive |
| 723.519@1.621 | up | 3.58 | 0.001 | 0.014 | positive |
| 576.958@9.702 | up | 1.68 | 0.001 | 0.014 | positive |
| 274.063@1.281 | up | 3.15 | 0.001 | 0.014 | positive |
| 800.543@1.198 | up | 1.72 | 0.001 | 0.014 | positive |
| 713.422@1.350 | up | 2.12 | 0.001 | 0.014 | positive |
| 729.425@1.402 | up | 2.37 | 0.001 | 0.014 | positive |
| 316.093@2.535 | up | 2.96 | 0.001 | 0.014 | positive |
| 679.424@1.321 | up | 1.38 | 0.001 | 0.014 | positive |
| 562.939@1.379 | down | 1.69 | 0.001 | 0.014 | positive |
| 660.673@1.274 | up | 11.80 | 0.001 | 0.014 | positive |
| 442.254@1.200 | up | 3.63 | 0.001 | 0.014 | positive |
| 807.153@1.274 | up | 4.24 | 0.001 | 0.014 | positive |
| 934.369@1.435 | up | 3.52 | 0.001 | 0.014 | positive |
| 938.270@1.439 | up | 5.64 | 0.001 | 0.014 | positive |
| 681.748@1.321 | up | 3.15 | 0.001 | 0.014 | positive |
| 146.105@1.305 | up | 1.42 | 0.001 | 0.014 | positive |
| 797.387@1.282 | up | 13.54 | 0.001 | 0.014 | positive |
| 731.437@1.422 | up | 2.44 | 0.001 | 0.014 | positive |
| 403.447@1.275 | up | 3.55 | 0.001 | 0.014 | positive |
| 627.413@1.356 | up | 4.23 | 0.001 | 0.014 | positive |
| 429.395@7.778 | up | 3.77 | 0.001 | 0.014 | positive |
| 485.278@9.312 | up | 1.88 | 0.001 | 0.014 | positive |
| 745.417@1.429 | up | 2.29 | 0.001 | 0.014 | positive |
| 807.152@1.274 | up | 4.64 | 0.001 | 0.014 | positive |
| 203.114@13.469 | up | 1.40 | 0.002 | 0.018 | positive |
| 742.247@1.458 | up | 2.34 | 0.002 | 0.018 | positive |
| 84.021@13.470 | up | 1.36 | 0.002 | 0.018 | positive |
| 261.192@9.629 | up | 1.49 | 0.002 | 0.018 | positive |
| 241.270@8.482 | down | 9.41 | 0.002 | 0.018 | positive |
| 886.824@1.347 | down | 1.31 | 0.002 | 0.018 | positive |
| 369.296@1.310 | down | 1.97 | 0.002 | 0.018 | positive |
| 203.238@13.469 | up | 1.48 | 0.002 | 0.018 | positive |
| 241.213@1.375 | down | 1.33 | 0.002 | 0.018 | positive |
| 550.538@1.403 | down | 1.39 | 0.002 | 0.018 | positive |
| 371.174@8.148 | up | 2.81 | 0.002 | 0.018 | positive |
| 592.920@1.348 | down | 1.44 | 0.002 | 0.018 | positive |
| 118.064@5.030 | up | 2.10 | 0.002 | 0.018 | positive |
| 677.407@1.334 | up | 1.94 | 0.002 | 0.018 | positive |
| 835.103@1.346 | down | 1.26 | 0.002 | 0.018 | positive |
| 684.237@1.451 | up | 2.80 | 0.002 | 0.018 | positive |
| 609.848@13.474 | up | 1.53 | 0.002 | 0.018 | positive |
| 295.251@1.347 | down | 1.22 | 0.003 | 0.023 | positive |
| 311.245@1.402 | down | 1.15 | 0.003 | 0.023 | positive |
| 200.046@5.028 | up | 2.27 | 0.003 | 0.023 | positive |
| 250.219@1.350 | down | 1.18 | 0.003 | 0.023 | positive |
| 243.186@9.691 | up | 1.16 | 0.003 | 0.023 | positive |
| 312.274@8.872 | down | 1.26 | 0.003 | 0.023 | positive |
| 575.489@1.357 | up | 1.22 | 0.003 | 0.023 | positive |
| 889.205@1.346 | down | 1.27 | 0.003 | 0.023 | positive |
| 637.414@1.375 | up | 3.05 | 0.003 | 0.023 | positive |
| 82.044@1.285 | up | 2.03 | 0.003 | 0.023 | positive |
| 144.043@13.472 | up | 1.39 | 0.003 | 0.023 | positive |
| 335.222@1.371 | down | 1.55 | 0.003 | 0.023 | positive |
| 731.430@1.422 | up | 1.87 | 0.003 | 0.023 | positive |
| 665.662@1.360 | up | 2.73 | 0.003 | 0.023 | positive |
| 557.309@8.349 | up | 2.01 | 0.003 | 0.023 | positive |
| 627.415@1.363 | up | 3.49 | 0.003 | 0.023 | positive |
| 422.163@1.266 | up | 3.31 | 0.001 | 0.026 | negative |
| 424.161@1.266 | up | 3.50 | 0.001 | 0.026 | negative |
| 446.206@1.268 | up | 4.12 | 0.001 | 0.026 | negative |
| 450.158@1.415 | up | 2.73 | 0.001 | 0.026 | negative |
| 424.336@1.266 | up | 4.79 | 0.001 | 0.026 | negative |
| 449.181@1.261 | up | 3.06 | 0.001 | 0.026 | negative |
| 446.206@1.262 | up | 4.46 | 0.001 | 0.026 | negative |
| 422.783@1.268 | up | 7.22 | 0.001 | 0.026 | negative |
| 422.730@1.268 | up | 5.75 | 0.001 | 0.026 | negative |
| 642.183@1.464 | up | 4.91 | 0.001 | 0.026 | negative |
| 450.338@1.416 | up | 3.42 | 0.001 | 0.026 | negative |
| 286.119@4.311 | up | 4.02 | 0.001 | 0.026 | negative |
| 416.028@10.351 | up | 3.24 | 0.001 | 0.026 | negative |
| 374.073@10.325 | up | 3.76 | 0.001 | 0.026 | negative |
| 356.174@1.269 | up | 4.59 | 0.001 | 0.026 | negative |
| 424.780@1.260 | up | 6.11 | 0.001 | 0.026 | negative |
| 434.038@10.351 | up | 3.60 | 0.001 | 0.026 | negative |
| 850.822@1.268 | up | 7.68 | 0.001 | 0.026 | negative |
| 235.997@2.353 | up | 4.04 | 0.001 | 0.026 | negative |
| 426.167@1.265 | up | 3.61 | 0.001 | 0.026 | negative |
| 468.169@4.330 | up | 1.90 | 0.001 | 0.026 | negative |
| 450.410@1.416 | up | 4.53 | 0.001 | 0.026 | negative |
| 386.186@1.269 | up | 4.89 | 0.001 | 0.026 | negative |
| 432.198@1.265 | up | 11.42 | 0.001 | 0.026 | negative |
| 500.174@1.273 | up | 4.45 | 0.001 | 0.026 | negative |
| 246.009@7.121 | up | 2.28 | 0.001 | 0.026 | negative |
| 479.888@10.357 | up | 3.39 | 0.001 | 0.026 | negative |
| 898.728@1.261 | up | 4.25 | 0.001 | 0.026 | negative |
| 460.091@4.301 | up | 14.40 | 0.001 | 0.026 | negative |
| 386.041@4.301 | up | 3.49 | 0.001 | 0.026 | negative |
| 478.073@5.044 | up | 6.00 | 0.001 | 0.026 | negative |
| 510.233@1.225 | up | 4.11 | 0.001 | 0.026 | negative |
| 210.004@3.861 | down | 3.46 | 0.001 | 0.026 | negative |
| 476.215@2.809 | up | 5.05 | 0.001 | 0.026 | negative |
| 305.992@3.872 | down | 2.58 | 0.001 | 0.026 | negative |
| 780.203@1.449 | up | 3.19 | 0.001 | 0.026 | negative |
| 466.138@7.514 | up | 4.02 | 0.001 | 0.026 | negative |
| 498.171@1.246 | up | 3.58 | 0.001 | 0.026 | negative |
| 478.069@5.049 | up | 4.88 | 0.001 | 0.026 | negative |
| 530.090@5.032 | up | 2.26 | 0.001 | 0.026 | negative |
| 795.160@1.430 | up | 3.44 | 0.001 | 0.026 | negative |
| 374.010@5.049 | up | 1.72 | 0.002 | 0.027 | negative |
| 110.998@2.919 | down | 1.53 | 0.002 | 0.027 | negative |
| 556.106@7.113 | up | 2.03 | 0.002 | 0.027 | negative |
| 403.089@2.912 | down | 1.56 | 0.002 | 0.027 | negative |
| 191.917@7.124 | up | 1.93 | 0.002 | 0.027 | negative |
| 318.104@6.006 | up | 2.95 | 0.002 | 0.027 | negative |
| 445.102@1.444 | down | 5.15 | 0.002 | 0.027 | negative |
| 614.181@1.414 | up | 2.28 | 0.002 | 0.027 | negative |
| 610.197@6.008 | up | 7.49 | 0.002 | 0.027 | negative |
| 58.042@7.453 | up | 1.31 | 0.002 | 0.027 | negative |
| 596.221@1.432 | up | 3.11 | 0.002 | 0.027 | negative |
| 492.135@5.042 | up | 1.93 | 0.002 | 0.027 | negative |
| 614.182@1.414 | up | 2.29 | 0.002 | 0.027 | negative |
| 388.099@5.039 | up | 1.78 | 0.002 | 0.027 | negative |
| 382.075@7.279 | up | 2.50 | 0.002 | 0.027 | negative |
| 514.127@4.986 | up | 2.70 | 0.002 | 0.027 | negative |
| 451.990@5.046 | up | 1.96 | 0.002 | 0.027 | negative |
| 524.092@1.410 | up | 1.92 | 0.002 | 0.027 | negative |
| 184.050@4.293 | up | 2.75 | 0.002 | 0.027 | negative |
| 258.131@1.961 | up | 1.95 | 0.002 | 0.027 | negative |
| 200.044@5.046 | up | 1.99 | 0.003 | 0.030 | negative |
| 492.135@5.046 | up | 1.95 | 0.003 | 0.030 | negative |
| 742.247@1.458 | up | 1.88 | 0.003 | 0.030 | negative |
| 88.015@2.780 | up | 2.43 | 0.003 | 0.030 | negative |
| 191.916@5.047 | up | 1.96 | 0.003 | 0.030 | negative |
| 413.833@17.826 | up | 1.05 | 0.003 | 0.030 | negative |
| 334.046@10.373 | up | 3.07 | 0.003 | 0.030 | negative |
| 112.015@1.554 | up | 2.45 | 0.003 | 0.030 | negative |
| 200.168@5.048 | up | 1.99 | 0.003 | 0.030 | negative |
| 106.026@3.856 | down | 2.55 | 0.003 | 0.030 | negative |
| 386.068@2.750 | up | 19.03 | 0.003 | 0.030 | negative |
| 304.018@5.064 | up | 1.79 | 0.003 | 0.030 | negative |
| 594.088@4.925 | up | 1.69 | 0.003 | 0.030 | negative |
| 218.013@2.383 | up | 2.12 | 0.003 | 0.030 | negative |
| 431.105@5.042 | up | 1.77 | 0.003 | 0.030 | negative |
| 419.973@7.156 | up | 1.77 | 0.003 | 0.030 | negative |
| 629.473@1.341 | down | 1.39 | 0.003 | 0.030 | negative |
| 278.035@5.046 | up | 2.70 | 0.003 | 0.030 | negative |
| 315.277@1.439 | down | 1.10 | 0.004 | 0.031 | positive |
| 431.232@1.255 | down | 1.72 | 0.004 | 0.031 | positive |
| 414.204@1.258 | down | 1.78 | 0.004 | 0.031 | positive |
| 556.448@1.344 | down | 1.18 | 0.004 | 0.031 | positive |
| 574.604@9.700 | up | 1.73 | 0.004 | 0.031 | positive |
| 774.844@1.278 | up | 2.39 | 0.004 | 0.031 | positive |
| 265.227@1.385 | down | 1.45 | 0.004 | 0.031 | positive |
| 134.075@1.259 | down | 1.42 | 0.004 | 0.031 | positive |
| 307.198@9.729 | up | 1.50 | 0.004 | 0.031 | positive |
| 465.170@1.438 | up | 1.31 | 0.004 | 0.031 | positive |
| 216.110@10.420 | down | 1.39 | 0.004 | 0.031 | positive |
| 183.155@1.382 | down | 1.59 | 0.004 | 0.031 | positive |
| 462.476@1.131 | down | 1.16 | 0.004 | 0.031 | positive |
| 264.014@7.119 | up | 1.95 | 0.004 | 0.039 | negative |
| 514.093@1.270 | up | 2.38 | 0.004 | 0.039 | negative |
| 509.820@17.860 | up | 1.05 | 0.004 | 0.039 | negative |
| 333.238@1.351 | down | 1.21 | 0.004 | 0.039 | negative |
| 200.044@7.119 | up | 1.87 | 0.004 | 0.039 | negative |
| 341.995@7.123 | up | 1.81 | 0.004 | 0.039 | negative |
| 272.001@1.128 | down | 1.07 | 0.004 | 0.039 | negative |
| 500.121@5.060 | up | 1.64 | 0.004 | 0.039 | negative |
| 596.108@5.056 | up | 1.95 | 0.004 | 0.039 | negative |
| 376.008@5.046 | up | 1.68 | 0.004 | 0.039 | negative |
| 398.996@8.771 | down | 1.17 | 0.004 | 0.039 | negative |
| 582.407@1.339 | down | 1.36 | 0.005 | 0.041 | positive |
| 280.137@1.253 | down | 1.77 | 0.005 | 0.041 | positive |
| 488.113@1.435 | up | 1.80 | 0.005 | 0.041 | positive |
| 532.242@9.096 | up | 1.07 | 0.005 | 0.041 | positive |
| 375.256@9.354 | up | 1.18 | 0.005 | 0.041 | positive |
| 231.143@11.714 | up | 1.26 | 0.005 | 0.041 | positive |
| **117.040@8.488** | down | 1.29 | 0.005 | 0.041 | positive |
| **90.047@1.263** | down | 2.69 | 0.005 | 0.041 | positive |
| **150.135@1.369** | down | 1.33 | 0.005 | 0.041 | positive |
| **700.137@1.418** | up | 1.67 | 0.005 | 0.041 | positive |
| **176.031@2.785** | up | 39.98 | 0.005 | 0.046 | negative |
| **684.009@1.538** | up | 62.14 | 0.005 | 0.046 | negative |
| **404.196@1.422** | up | 2.57 | 0.005 | 0.046 | negative |
| **100.015@1.534** | up | 1.80 | 0.005 | 0.046 | negative |
| **210.037@1.998** | up | 2.09 | 0.005 | 0.046 | negative |
| **275.956@2.798** | up | 21.38 | 0.005 | 0.046 | negative |
| **419.152@5.090** | up | 1.90 | 0.005 | 0.046 | negative |
| **726.066@1.393** | up | 6.12 | 0.005 | 0.046 | negative |
| **257.946@2.783** | up | 16.93 | 0.005 | 0.046 | negative |
| **271.986@2.887** | down | 1.59 | 0.005 | 0.046 | negative |
| **364.982@2.818** | up | 12.87 | 0.005 | 0.046 | negative |
| **586.201@1.249** | up | 2.15 | 0.005 | 0.046 | negative |
| **726.063@1.393** | up | 6.03 | 0.005 | 0.046 | negative |
| **337.993@4.729** | down | 1.50 | 0.005 | 0.046 | negative |
| **530.198@1.419** | up | 4.61 | 0.005 | 0.046 | negative |
| **642.203@6.955** | up | 1.91 | 0.005 | 0.046 | negative |
| **482.181@1.290** | up | 2.47 | 0.005 | 0.046 | negative |
| **630.182@1.478** | up | 3.10 | 0.005 | 0.046 | negative |
| **618.325@4.233** | up | 4.49 | 0.005 | 0.046 | negative |

**Table S5.**
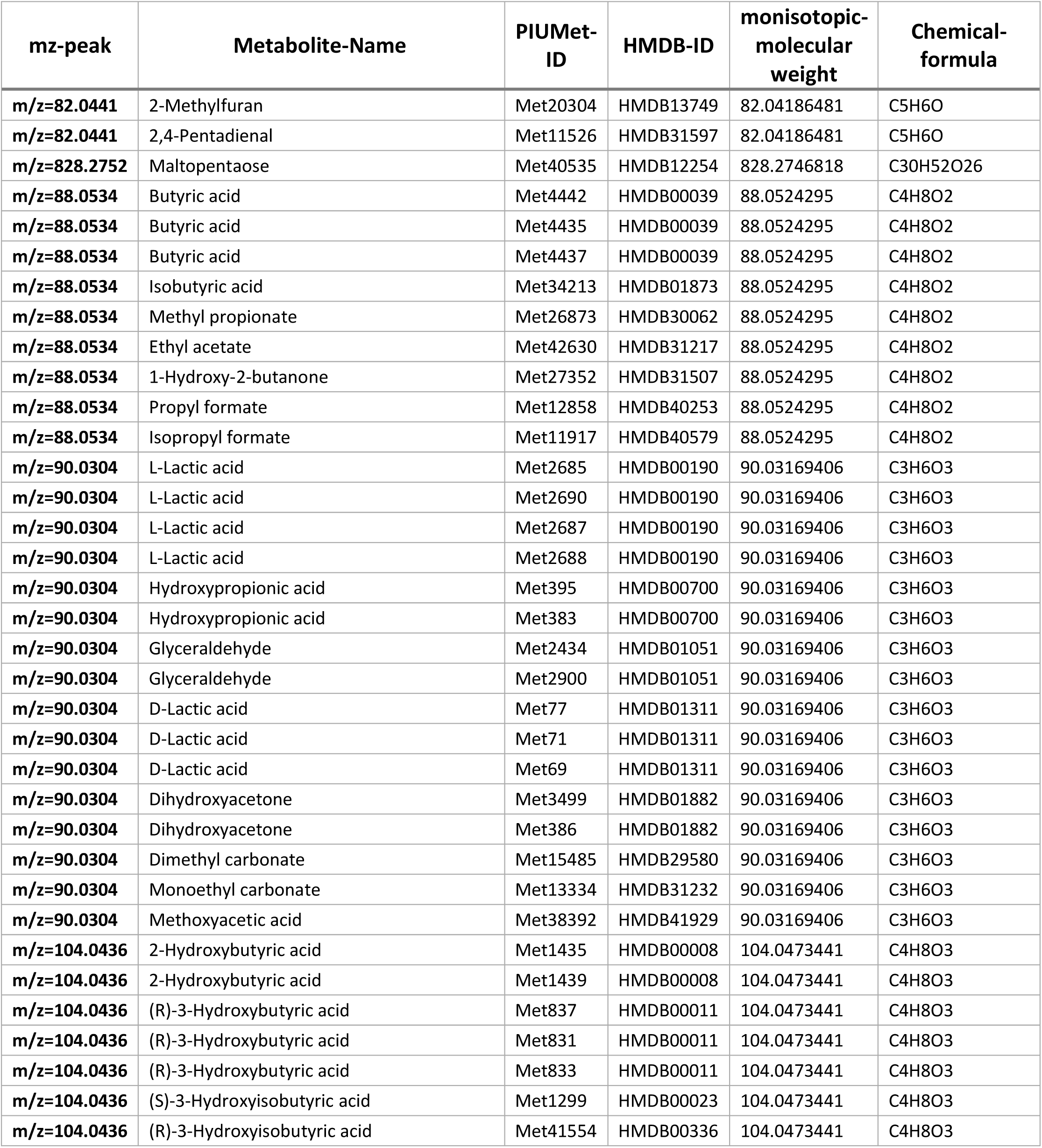
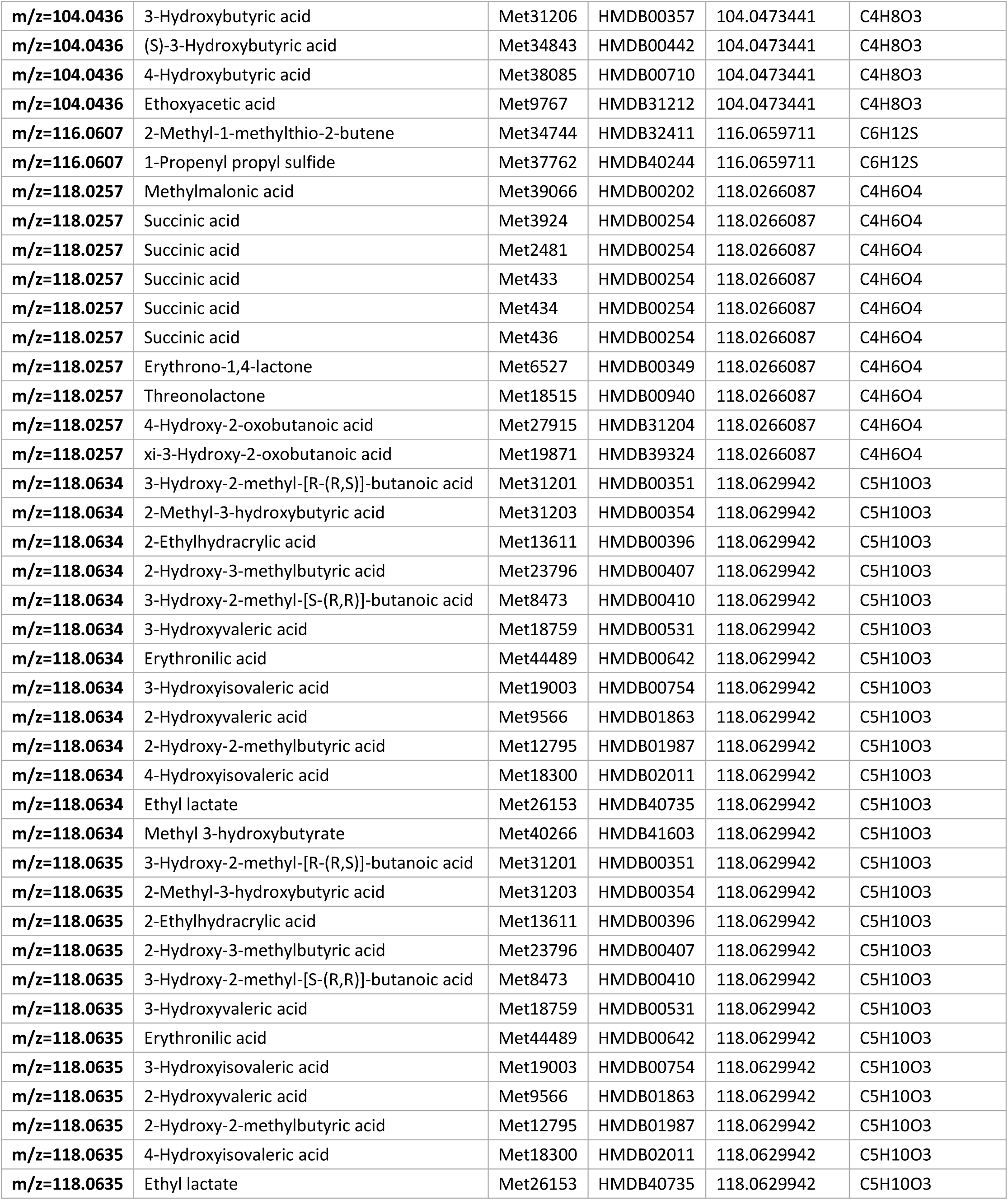
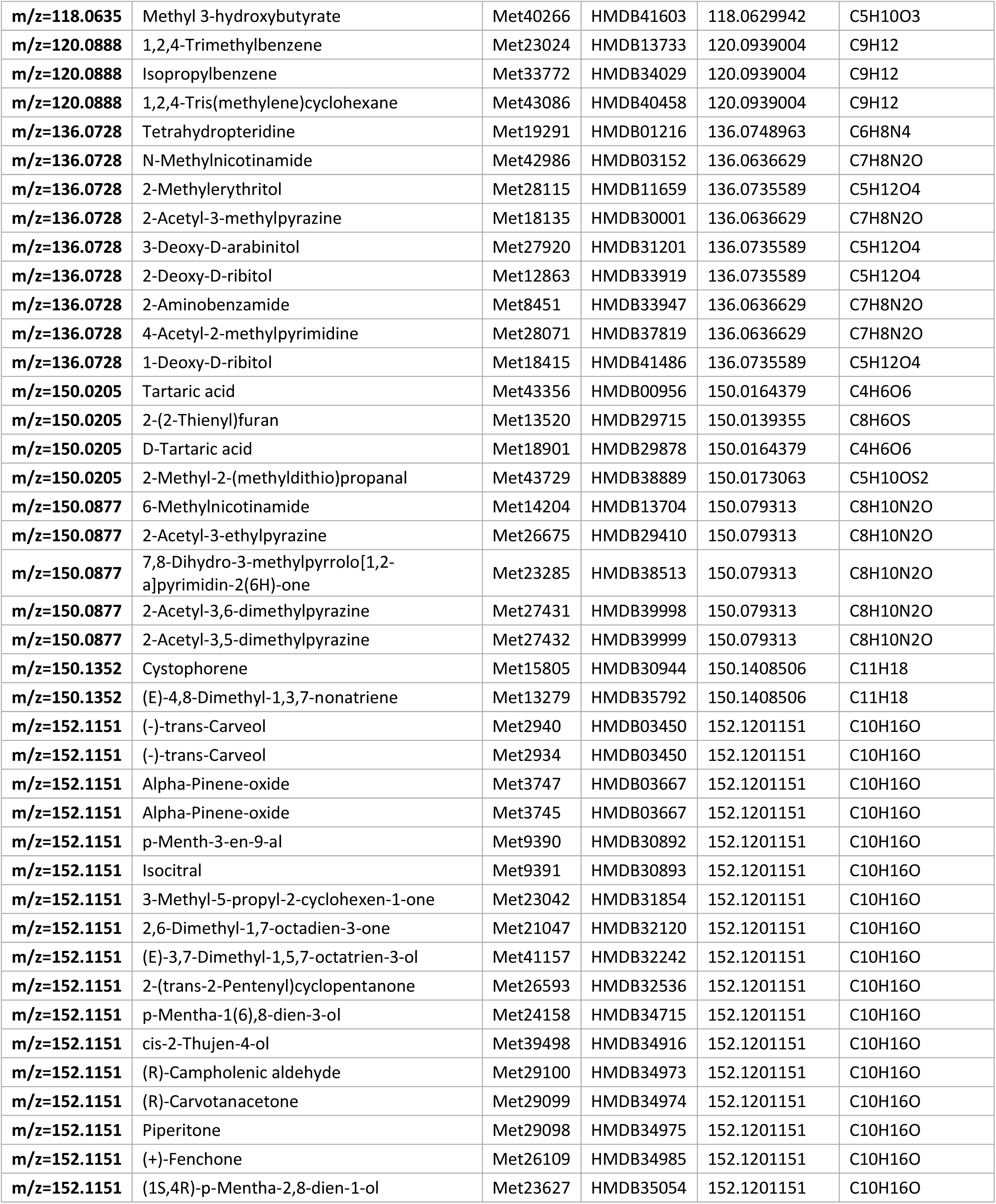
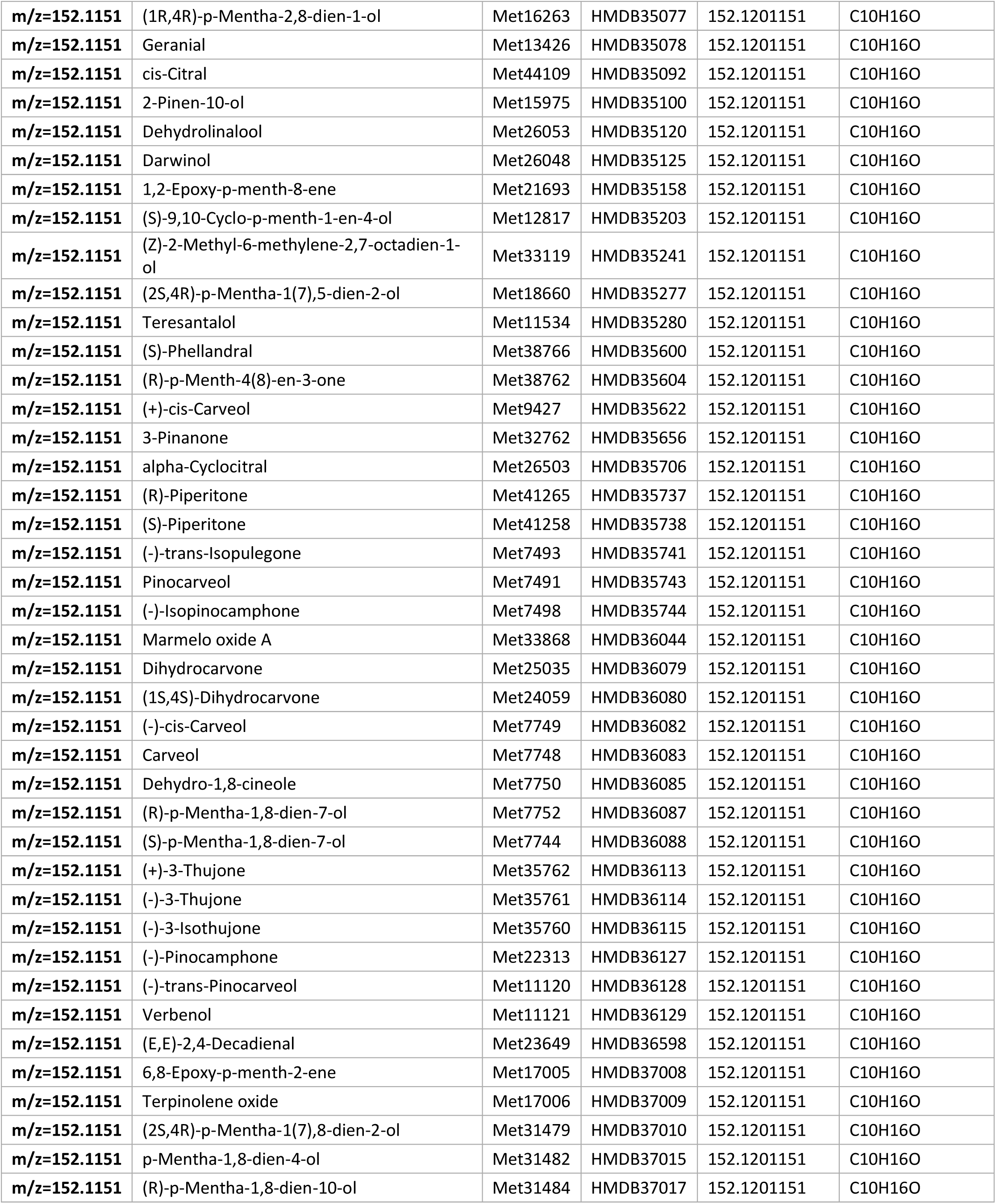
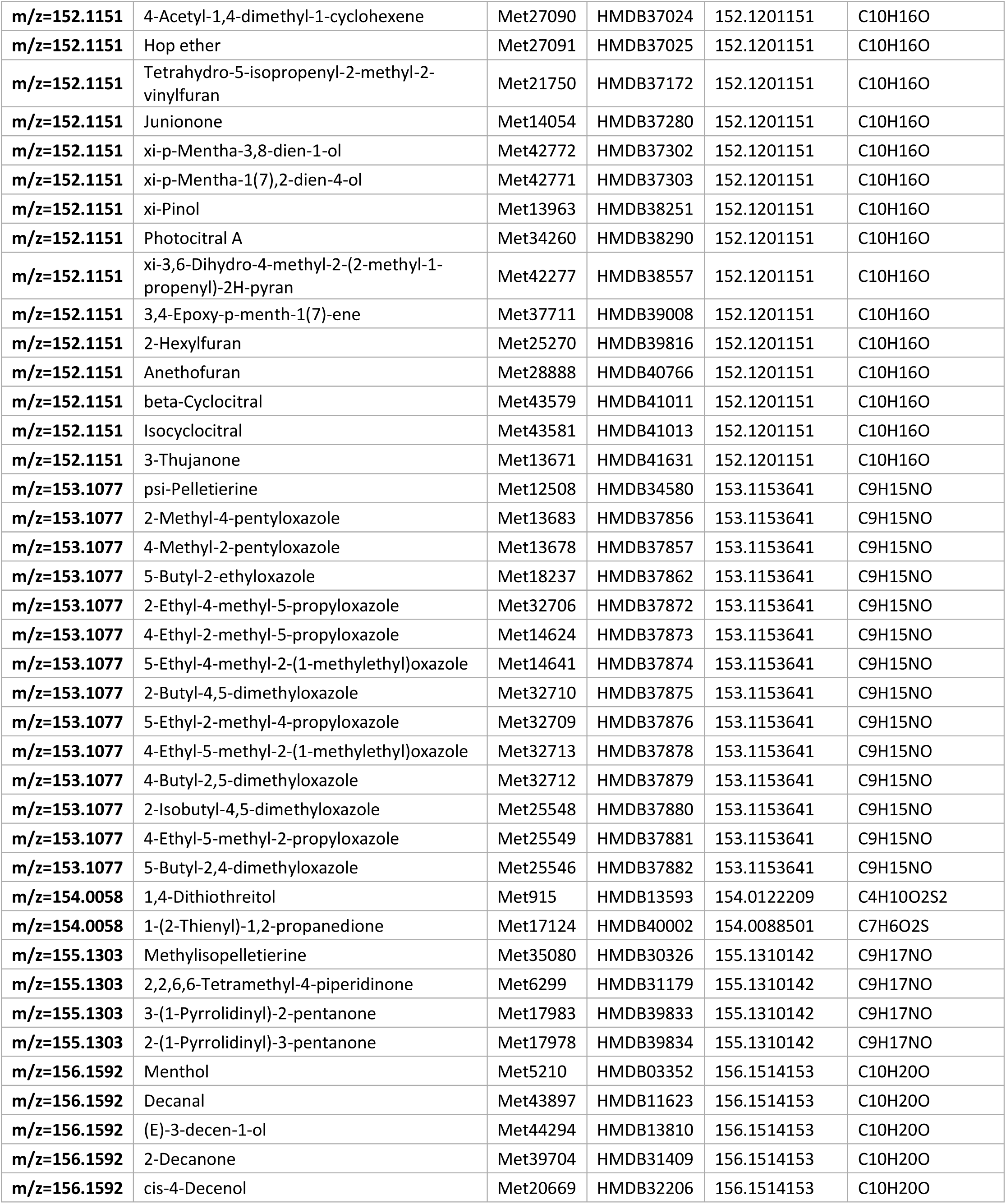
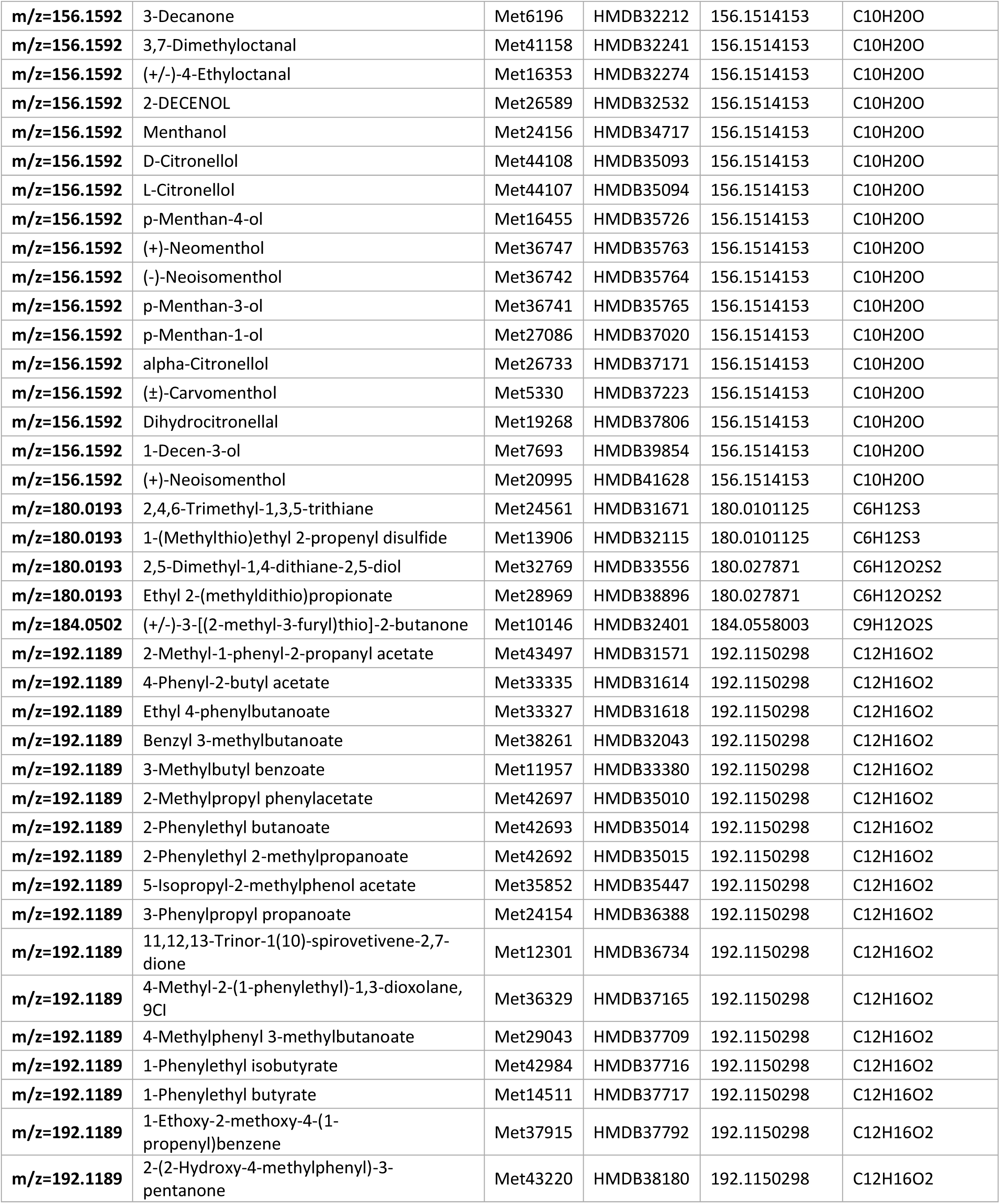
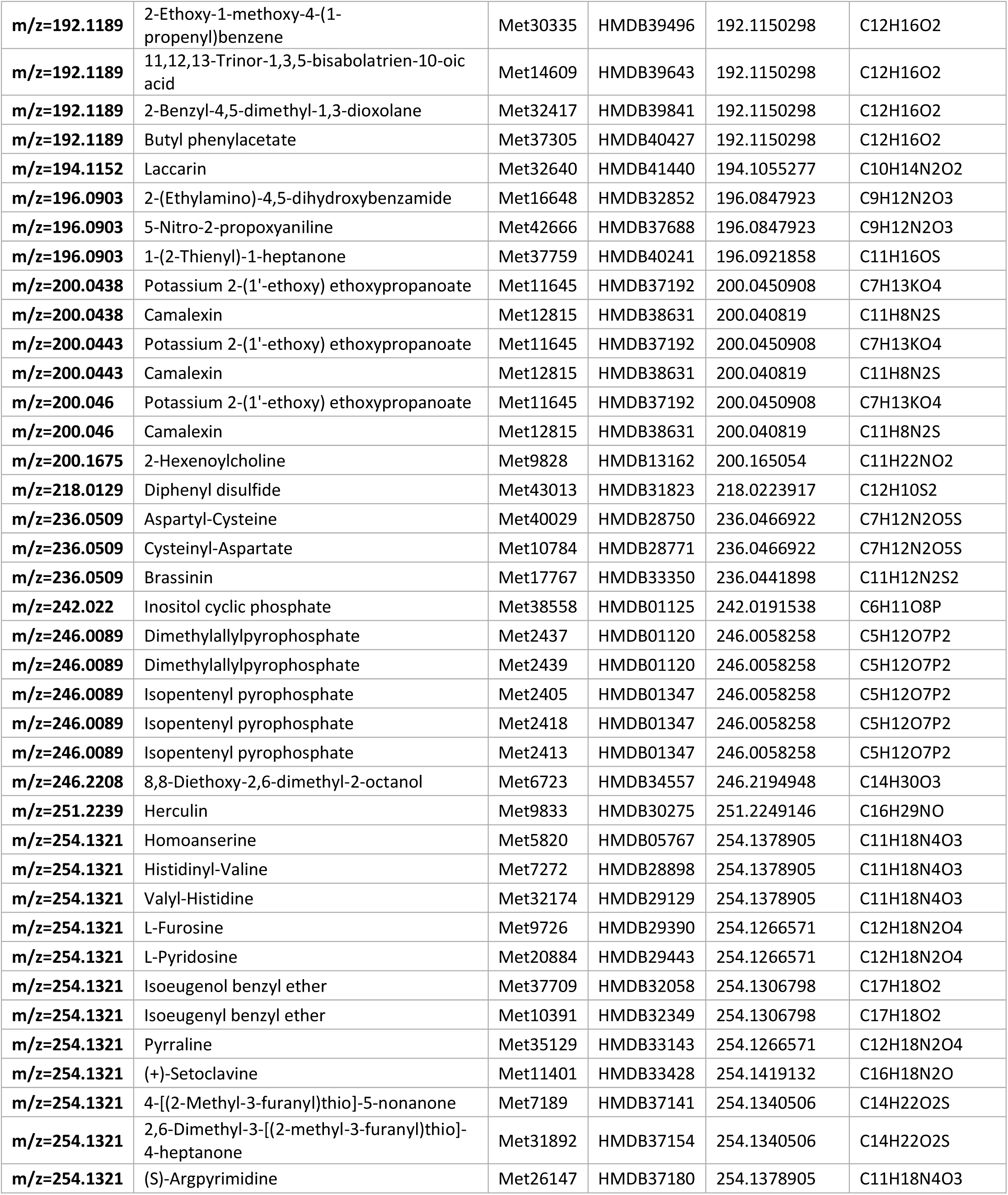
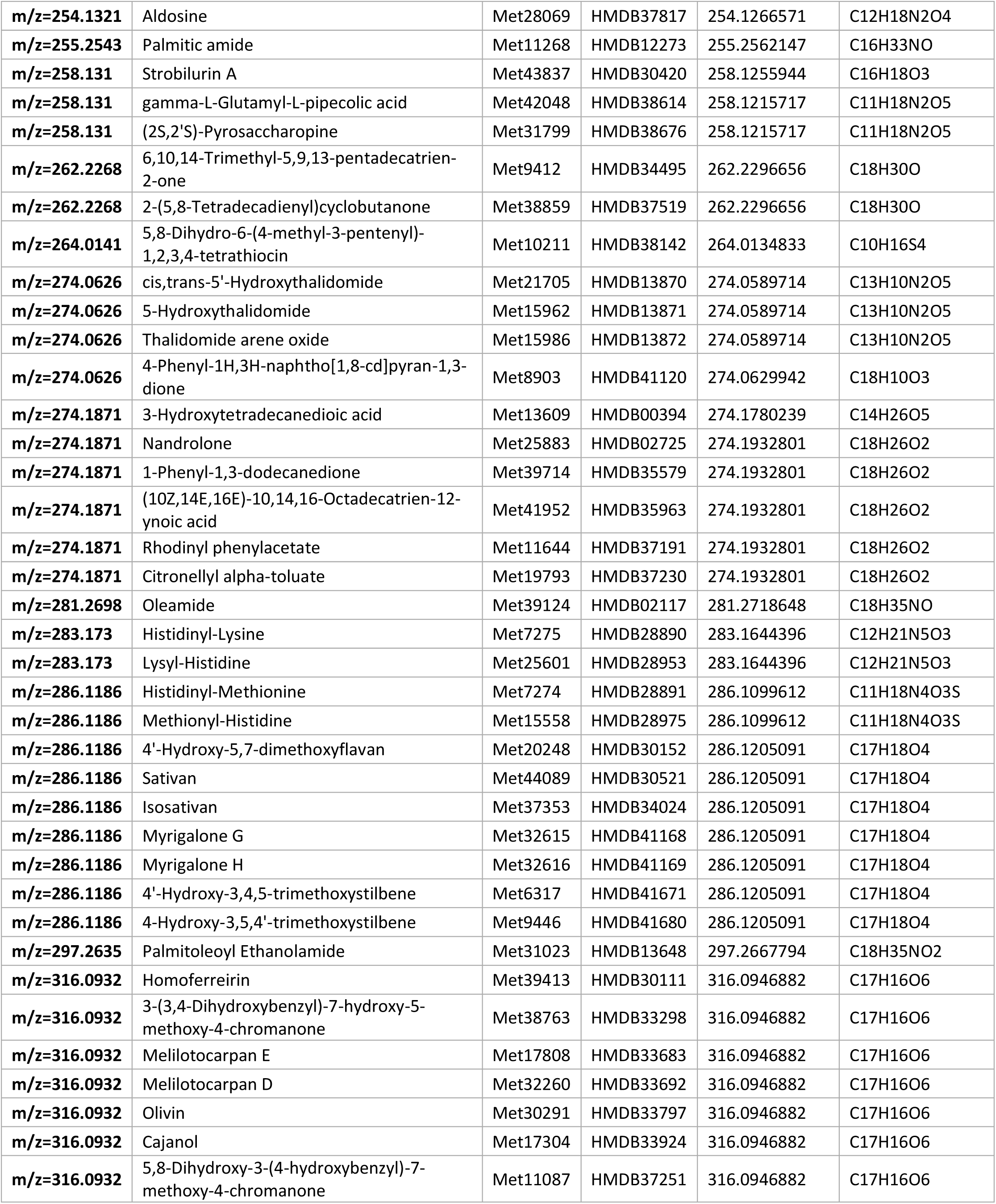
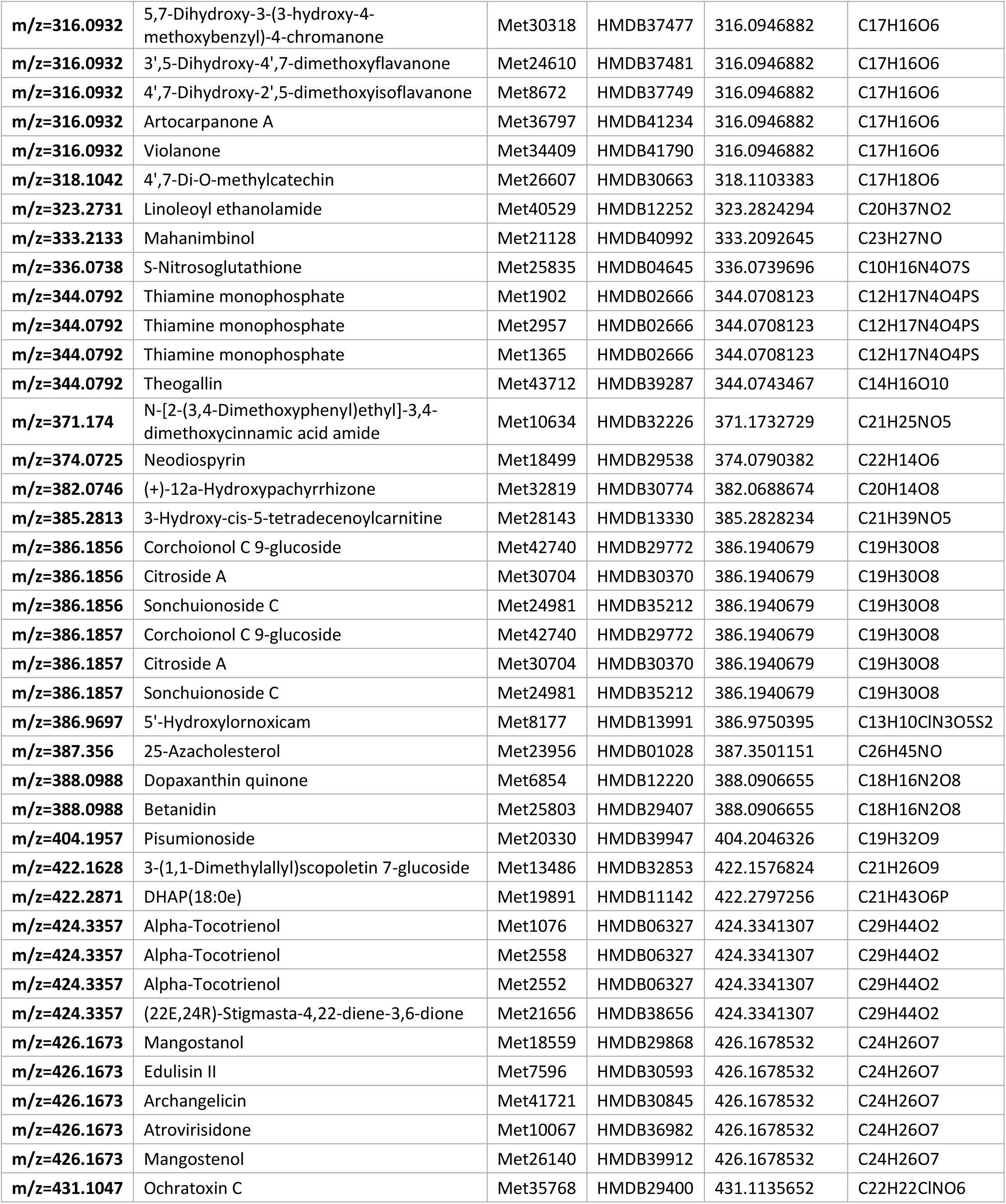
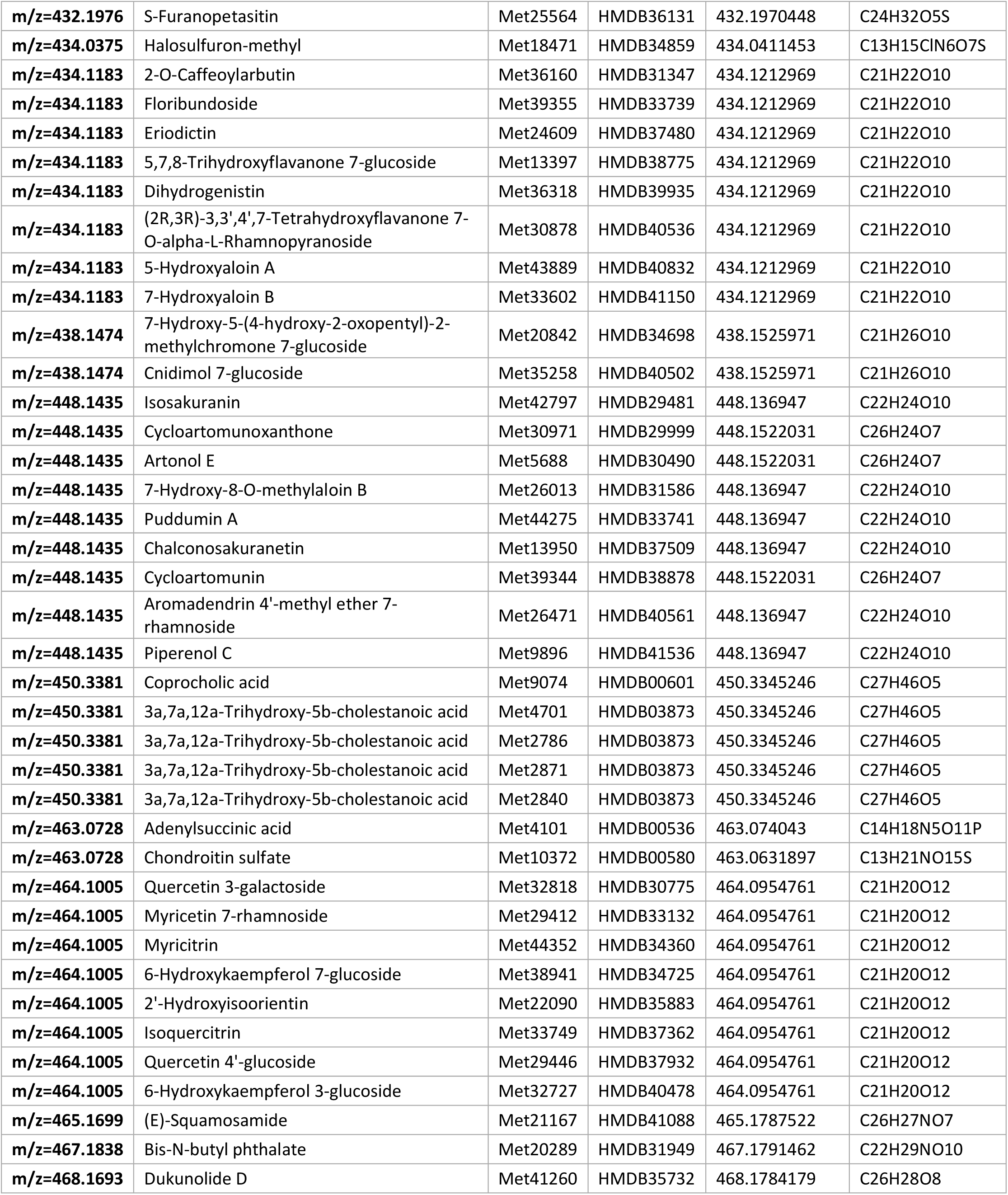
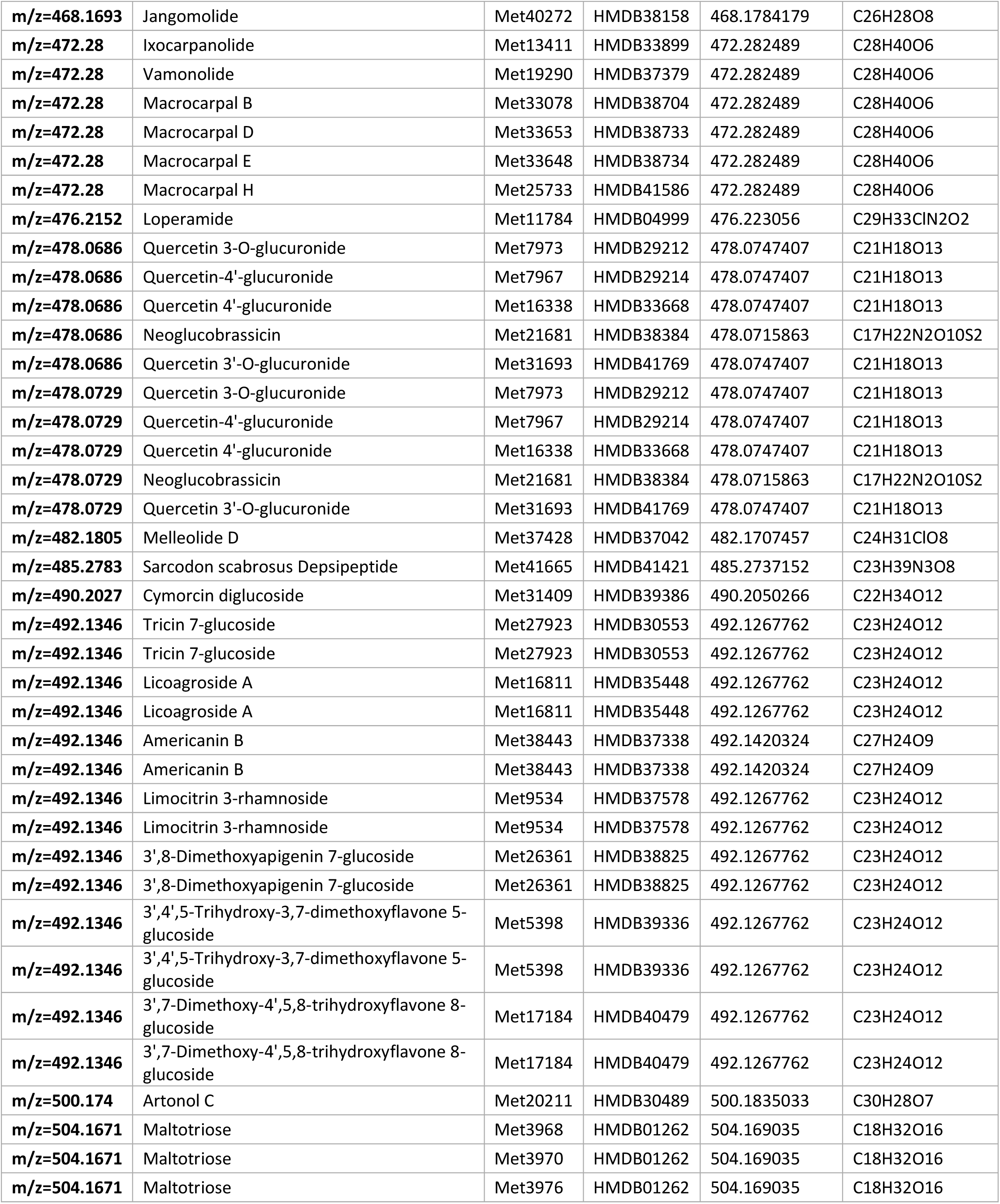
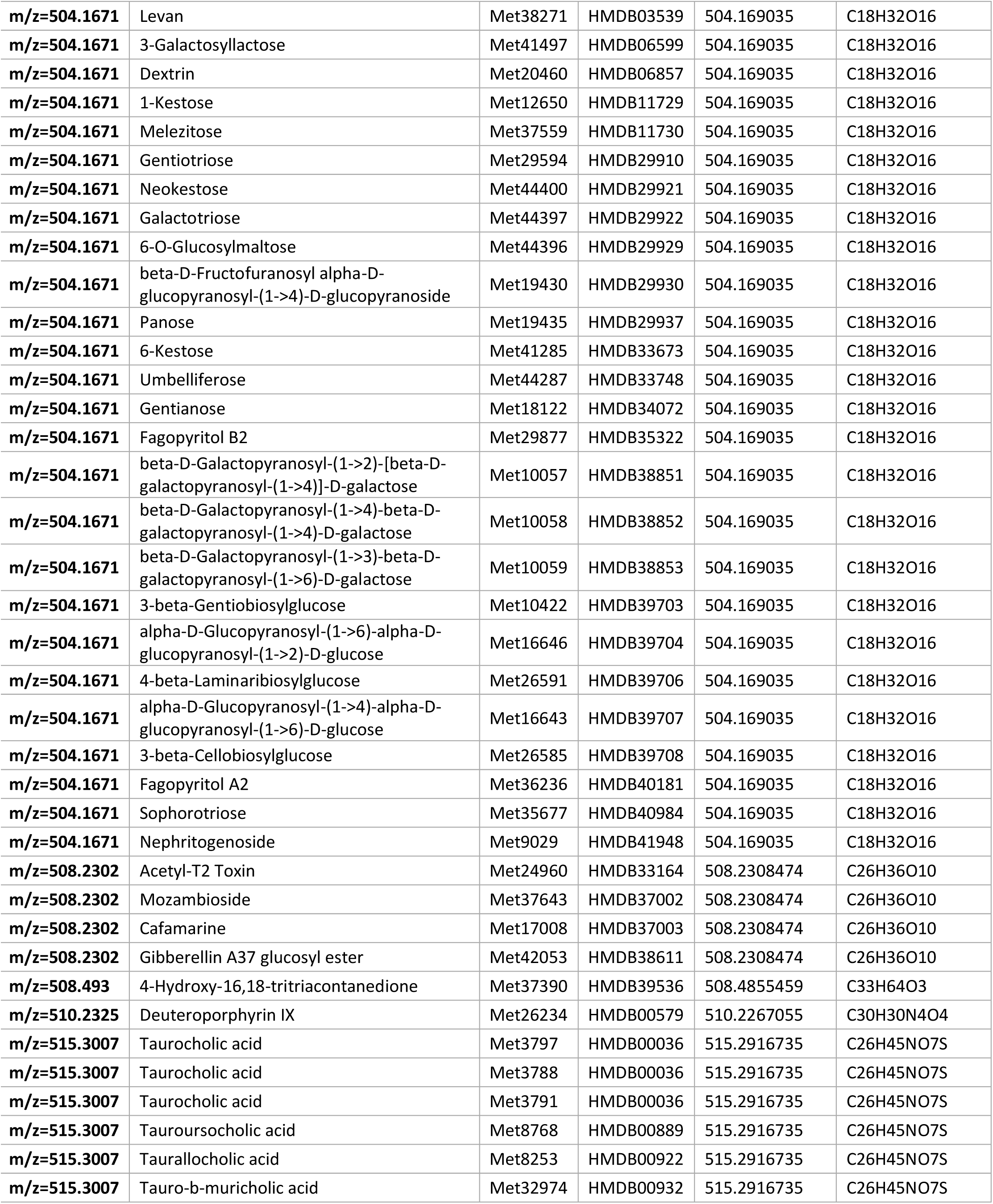
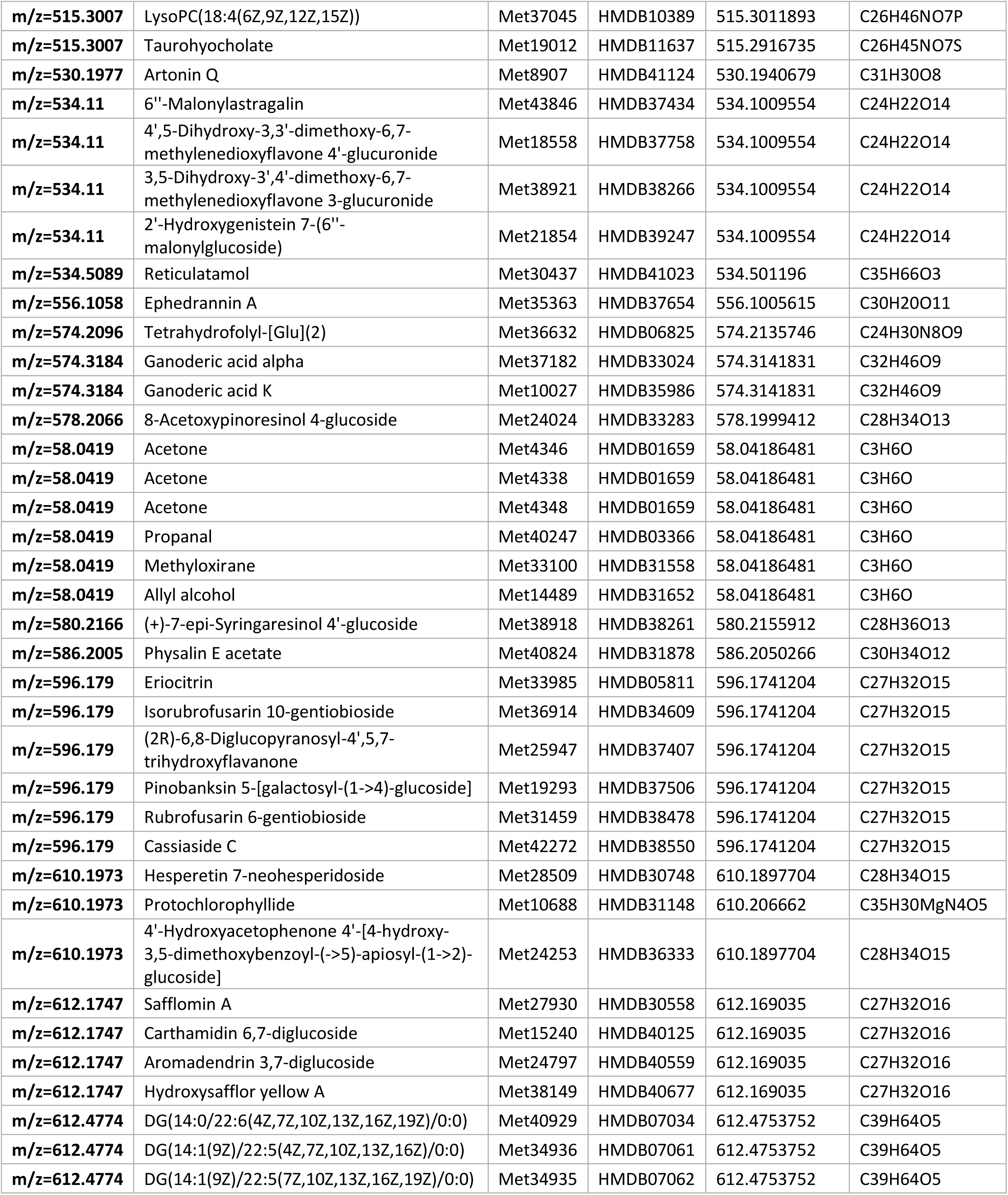
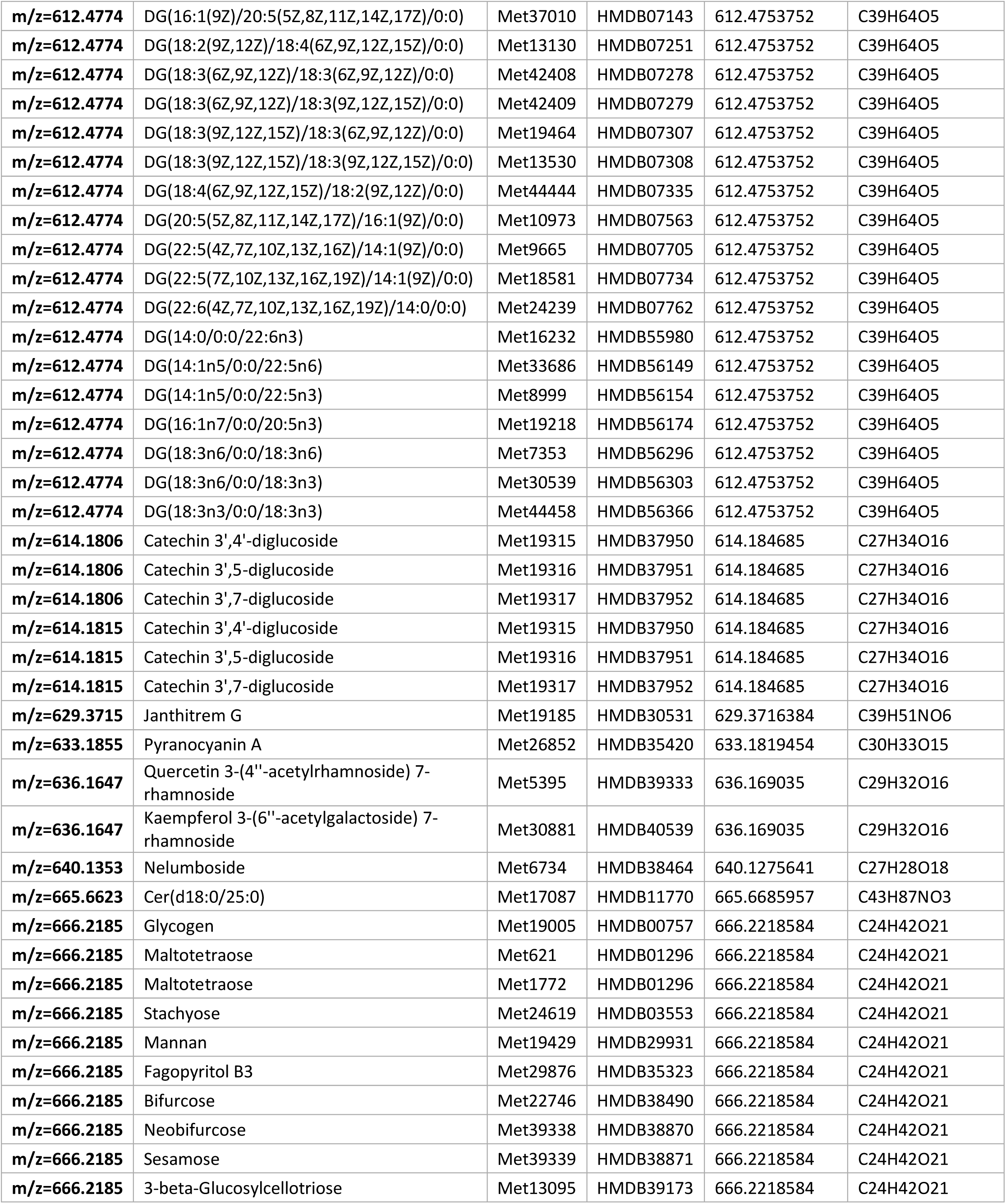
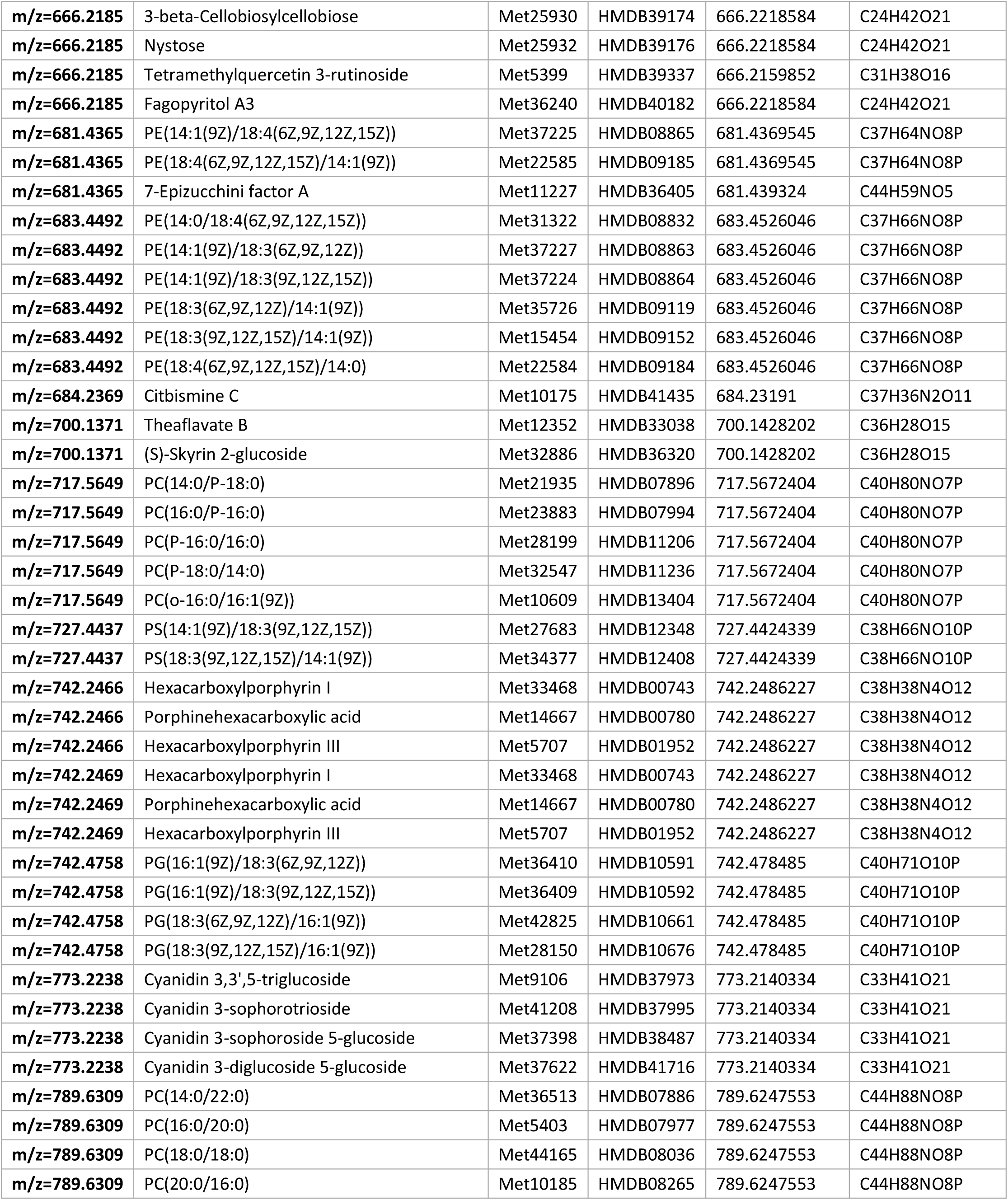
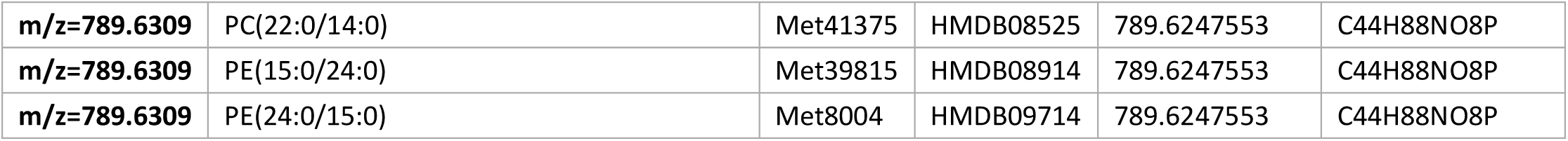
Putative identities of untargeted metabolites from the EC of APOE3 vs. E4 mice by PIUMet analysis

