## Supplementary figures and images for "*APOE4* is Associated with Differential Regional Vulnerability to Bioenergetic Deficits in Aged *APOE* Mice"

### Supplementary file 1

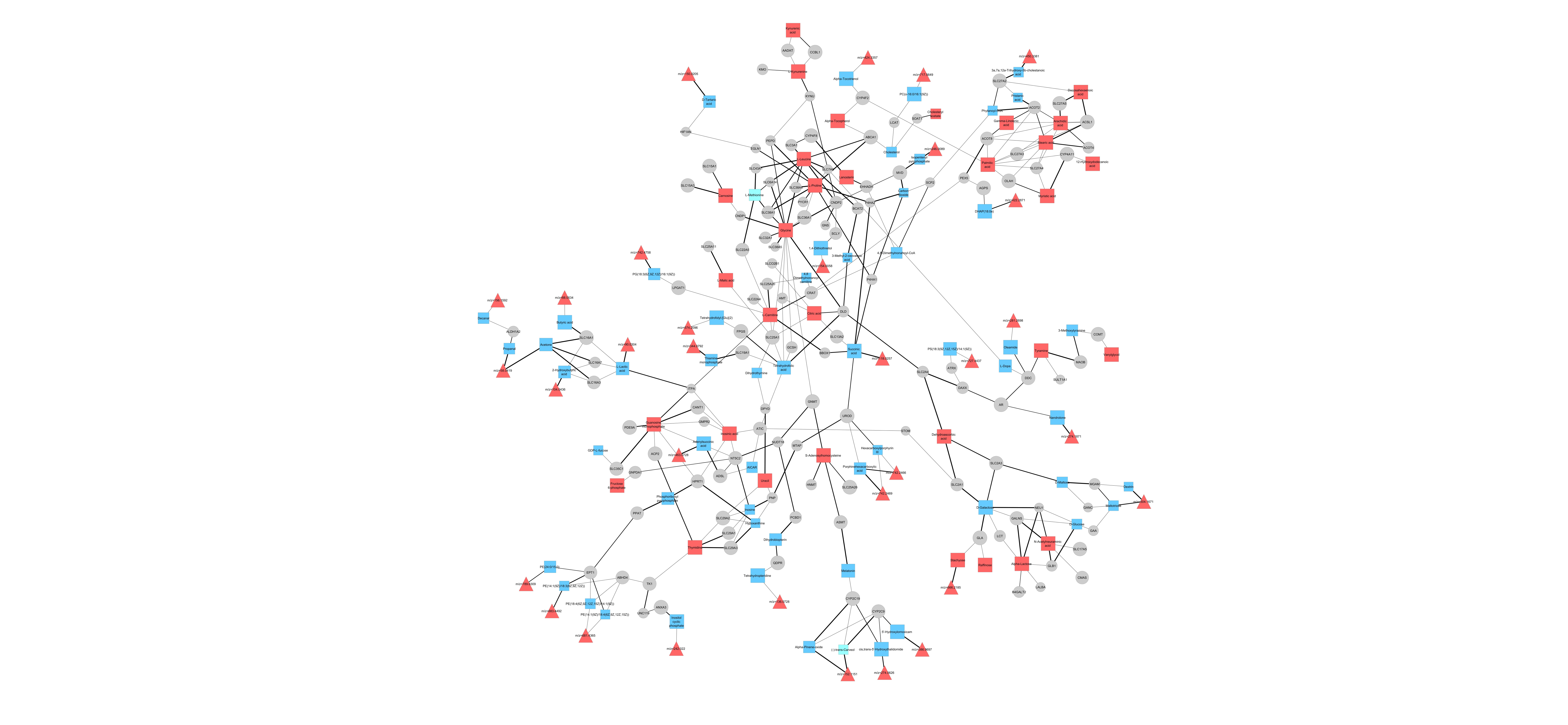
